# Dual Carbohydrate Recognition by the Chitinase-like Protein CHI3L1 Through Distinct Glycosaminoglycan and Chitin-Binding Interfaces

**DOI:** 10.64898/2026.06.23.733983

**Authors:** Önder Kurç, Nick Rähse, Mohanraj Gopalswamy, Alexandra Großdorf, Christian Gorzelanny, Jonathan Cramer, Holger Gohlke

## Abstract

CHI3L1 (YKL-40) is a chitinase-like glycoprotein involved in immune regulation, tissue remodeling, and cancer, yet the molecular principles governing its glycan interactions remain incompletely defined. Previous reports suggested that CHI3L1 can bind to chitin oligosaccharides (COS) and glycosaminoglycan (GAG) ligands, however, the molecular basis and binding sites underlying these interactions remain controversial. Here, a combination of biophysical and computational methods is employed to shed light on carbohydrate interactions of the protein and delineate a potential crosstalk between its glycan-binding interfaces. Our results demonstrate that COS and GAGs bind to distinct, non-overlapping sites on CHI3L1. Both ligand classes exhibit a strong dependence of binding affinity on the degree of polymerization. Molecular dynamics simulations, supported by mutational analysis, identify a GAG-binding site centered on residues R144, R145, and K147 and reveal an additional distal interaction site for longer GAG ligands. Biophysical and biochemical assays fail to confirm a previously proposed allo-or orthosteric interaction between both binding sites. However, physiologically relevant protein–protein interactions mediated by the chitin binding site of CHI3L1 are differentially regulated by GAG and COS ligands. COS inhibit binding of galectin-3 to CHI3L1, whereas GAG ligands enhance the affinity between the proteins by ca. 14-fold. Together, these findings establish CHI3L1 as a dual carbohydrate-binding protein with distinct recognition interfaces and reveal a previously unrecognized role for GAGs in modulating CHI3L1-mediated signaling interactions.

## Introduction

Human chitinase-3-like 1 (CHI3L1), also known as YKL-40 or human cartilage glycoprotein 39 (HC-gp39), is a secreted glycoprotein belonging to the glycoside hydrolase family 18 (GH 18). Alongside its homolog CHI3L2 (YKL-39), it was first identified in cultures of MG-63 human osteosarcoma cells.(1) Unlike true chitinases, CHI3L1 and other chitinase-like proteins lack enzymatic activity due to the absence of essential catalytic residues in their chitin-binding site. However, recent work has suggested that the inactivity of CHI3L1 is not solely attributable to the mutated catalytic motif but also to variations in non-catalytic residues that affect substrate positioning.(2) In humans, CHI3L1 is expressed by multiple immune cell types, most prominently macrophages, as well as endothelial cells, smooth muscle cells, neutrophils, synoviocytes, chondrocytes, and various tumor cells.(3) Functionally, the protein acts as a mediator of inflammation, while contributing to cell proliferation and tissue remodeling. Dysregulated CHI3L1 expression has been implicated in numerous pathological conditions, including inflammatory diseases,(4–8) infectious diseases,(9–12), and cancer, where it can support immune evasion of cancer cells and promote tumor progression.(13–17) Notably, most physiological functions of CHI3L1 are generally considered to be independent of chitin.

The high diversity of physiological functions attributed to CHI3L1 is often linked to the diverse nature of its binding partners. It has been established that CHI3L1 can bind carbohydrate ligands, such as chitin oligosaccharides (COS) and various glycosaminoglycans (GAGs), including heparin, heparan sulfate, and hyaluronan.(18) In addition, it interacts with protein components of the extracellular matrix (collagen I, II, and III) and excreted proteins, such as galectin-3 and galectin-3 binding protein.(19, 20) However, its interactions with cell surface receptors, including syndecans, CD44v3, TMEM219, and IL-13Rα2, are primarily assumed to mediate its effector functions and pathway activation.(3, 21, 22) A particularly interesting mechanism involves the interplay among CHI3L1, galectin-3, TMEM219, and IL-13Rα2. CHI3L1 is known to promote anti-apoptotic signaling in epithelial cells through the formation of a CHI3L1–IL-13Rα2–TMEM219 complex, the so-called chitosome complex, which activates PI3K/AKT and ERK/MAPK pathways.(20) It has been demonstrated that galectin-3 interacts with CHI3L1 and IL-13Rα2 in a manner that interferes with TMEM219 recruitment to IL-13Rα2. This competitive interaction disrupts formation of the canonical CHI3L1–IL-13Rα2–TMEM219 complex, thereby reducing CHI3L1-mediated anti-apoptotic signaling in epithelial cells.(20) In the context of Hermansky-Pudlak Syndrome (HPS), elevated galectin-3 expression alters this balance, contributing to exaggerated epithelial injury and fibroproliferative repair by modifying the anti-apoptotic and profibrotic functions of CHI3L1 in a tissue compartment-specific manner.

Due to the central role of CHI3L1 in cell signaling and disease, the protein has attracted significant attention as a potential therapeutic target. Considerable efforts have been made to discover small molecule-based modulators of CHI3L1.(23–25) In this context, the interaction between the canonical chitin binding site and distal regions interacting with GAGs has sparked significant interest. Based on the available data, GAGs and COS are assumed to bind to CHI3L1 at distinct binding sites. The chitin binding site in GH18 chitinase proteins is generally well known, and multiple crystal structures of CHI3L1 in complex with COS are available.(26, 27) The nature and location of the GAG binding site(s), however, are less clear and still a subject of ongoing discussion. Previous research has identified two potential binding sites for GAGs, a consensus heparin-binding motif in region 144–147 and a KR-rich domain in region 334–345 (Figure 1A).(27–29) In addition, some claims of a direct interaction of GAGs with the chitin binding site have been put forward.(18) It has also been postulated that GAG ligands and COS bind CHI3L1 in a competitive manner.(30) However, direct evidence of an orthosteric or allosteric competition mechanism is currently lacking. Recent work has suggested that COS can partially modulate CHI3L1-GAG interactions,(30) raising the possibility of allosteric regulation. However, these conclusions were largely based on indirect binding assays, and the molecular basis of this cross-talk remains largely unresolved. We recently proposed that simultaneous interaction with COS and GAG ligands has a functional relevance in a pro-inflammatory setting apart from the canonical chitosome complex.(31) Accordingly, COS are bound by soluble CHI3L1 and recruited to the glycocalyx of immune cells via GAG interactions. After concentration on the cell membrane, the immunogenic chitin fragments are subsequently released in the vicinity of cognate immune receptors, such as TLR2.

**Figure 1.**
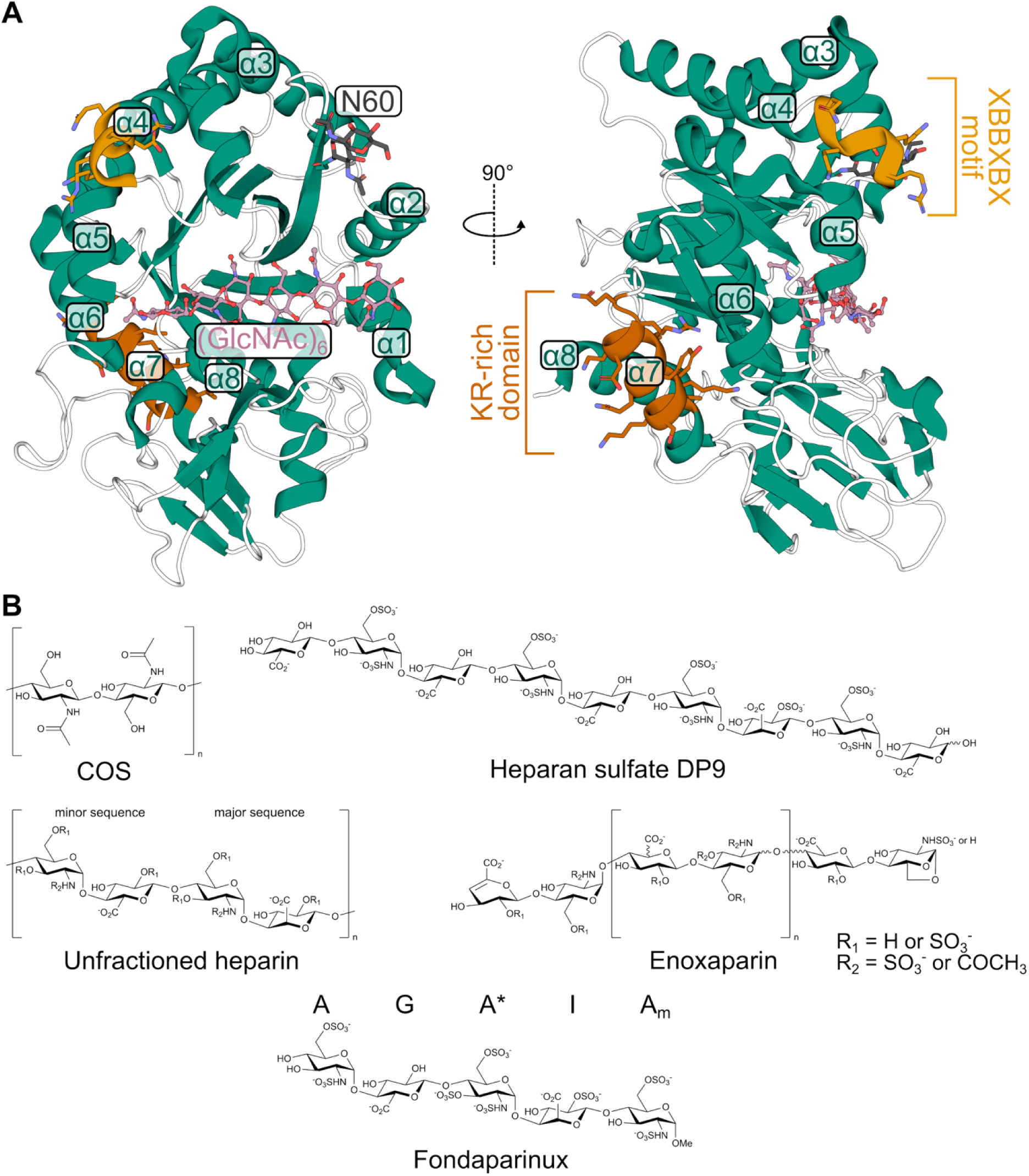
CHI3L1 structure and chemical structures of ligands characterized for binding affinity. (A) Crystal structure (PDB ID: 1HJW)(26) of CHI3L1 in complex with (GlcNAc)_6_ (violet ball-and-stick model). TIM-barrel helices are labeled α1-α8. The N-linked (N60) glycosylation site is indicated ((GlcNAc)_2_; black sticks). The proposed heparin-binding regions are highlighted in bright (^143^GRRDKQ^148^)(27, 28) and dark orange (residues 334-345)(29), respectively. (B) Structures of the studied COS and heparin/HS ligands.(32, 33) The sugar moieties of Fondaparinux are labeled according to ref.(34)

In this study, we investigate the interactions between CHI3L1 and its carbohydrate ligands. COS of different DP ((GlcNAc)_4–6_) are considered in addition to mono-and polydisperse GAGs: the chemically-defined heparin pentasaccharide fragment Fondaparinux, a chemically-defined heparan sulfate nonamer (HS9), the low-molecular weight heparin preparation Enoxaparin, and full-length unfractioned heparin (UFH, Figure 1B). By combining molecular modeling and simulations, as well as biophysical and biochemical methods, we aim to dissect the nature of the individual binding sites and delineate the dynamic interplay between their ligands. These findings contribute to a deeper understanding of how GAG and COS ligands modulate CHI3L1 function at the molecular level and establish a fundamental baseline for further investigations of CHI3L1 as a drug target.

## MATERIALS AND METHODS

### Plasmid Design and Protein Expression

The CHI3L1 gene was amplified from cDNA of human macrophages and cloned into the pIRES neo2 vector (Takara Bio, San Jose, USA). Point mutations were used to prevent chitin (W31A; W99A; W352A) or HS (R144A; R145A; K147A) binding or to activate (A138D; L140E) the protein. Mutagenesis for the chitin-binding mutant (CBmut CHI3L1) and the catalytically active CHI3L1 (Cat CHI3L1) was performed using the QuikChange Multi Site-Directed Mutagenesis Kit (Agilent, Santa Clara, USA), while Q5® Site-Directed Mutagenesis Kit (NewEngland Biolabs, Ipswich, USA) was used to produce the heparin-binding mutant (HepBmut CHI3L1). Mutagenesis was performed according to the kit instructions. After transformation into *E. coli* Mach1 T1 cells, all plasmids were extracted and point mutations were validated by Sanger sequencing.

For expression, wild-type and mutant CHI3L1 gene-containing plasmids were transfected into HEK293F cells and protein expressing cells were selected using geneticin. Cells were grown in DMEM (1% penicillin-streptomycin, 1% L-glycine, 10% FCS) medium; for recombinant protein production, culture medium was replaced with serum-reduced Opti-MEM medium (Gibco) and cells were incubated at 37 °C with 5% CO_2_ for 24 h. The cell supernatant was centrifuged for 10 min at 4000 g and passed over a Ni-NTA column (5 mL HiTrap FF (Cytiva)) with the help of a liquid chromatography system (ÄKTA go, Cytiva) at room temperature using a constant flow rate of 1 mL/min and a linear imidazole gradient (20 CV, 0-500 mM). Fractions containing the His-tagged recombinant protein were desalted using Zeba Spin desalting columns 7K MWCO 5 mL (Thermo Fisher) and buffered in PBS according to the manufacturer’s instructions. Protein purity was determined by polyacrylamide gel electrophoresis (NuPAGE™ 4 to 12%, Bis-Tris, 1.0-1.5 mm) mini protein gels. Gels were stained with Coomassie Brilliant Blue (CBB, Wako).

### Saturation Transfer Difference Nuclear Magnetic Resonance

STD-NMR experiments were performed as previously described.(35) NMR experiments were performed in a 3 mm NMR tube at 25 °C on a Bruker Avance III HD spectrometer operating at 750 MHz, equipped with a 5 mm triple resonance TCI (^1^H, ^13^C, ^15^N) cryoprobe and a shielded z-gradient. Samples were prepared in 200 µL volume with a final concentration of 7.5 µM of wild-type CHI3L1 (6.5 µM in the case of CBmut CHI3L1) and 750 µM of Fondaparinux in PBS buffer (90% (v/v)), 10% (v/v) D_2_O. Selective saturation of protein resonances (on-resonance spectrum) was performed by irradiating at 0.467 ppm for a total saturation time of 3 s with a relaxation delay of 3 s. For the reference spectrum (off-resonance), the samples were irradiated at −40 ppm. The STD-NMR spectra were acquired with 4928 scans, and the subtraction of the reference spectrum was performed internally via phase cycling after every scan to obtain the STD effect.(36) The ^1^H NMR spectrum of Fondaparinux was assigned by using previously published data by Hricovíni et al.(34) The STD intensity of the A*1 proton was set to 100% as a reference, and the relative intensities were determined. Based on these STD effects, the binding epitope was mapped on the structure of Fondaparinux. Data was processed and analyzed by the TopSpin 4.5.0 (Bruker BioSpin) software. Sodium 2,2-dimethyl-2-silapentane-5-sulfonate (DSS) was used for chemical shift referencing.

### Thermal Unfolding Profile

Nano-Differential Scanning Fluorimetry (nanoDSF) measurements were performed with a Nanotemper Prometheus instrument (Nanotemper, München, Germany). For thermal stability assessments, wild-type or mutant CHI3L1 was loaded into capillaries at a concentration of 2.5-5 µM. To evaluate ligand binding effects, CHI3L1 was briefly incubated with 400 µM (GlcNAc)_6_ or 500 µM Fondaparinux prior to being loaded into the capillaries. The data was collected and evaluated with the Prometheus Control software. Unfolding transitions were measured by tracking the fluorescence ratio (350 nm / 330 nm) as a function of temperature, and the corresponding first derivatives were used to determine the melting temperatures (*T*_m_). Thermal unfolding curves were plotted with Graphpad Prism v. 8.4.1

### Surface Plasmon Resonance (SPR)

SPR experiments were performed on a Biacore X100 instrument at 25 °C. For small molecule ligands (M < 2000 g mol^−1^), CHI3L1 (40 kDa, >90% pure based on SDS-PAGE) was immobilized at a concentration of 25 μg/mL in 10 mM sodium acetate, pH 5.0 at low density (2000-4000 RU) on a HC200M hydrogel chip (Xantec, Düsseldorf, Germany) by using amine-coupling chemistry with a 1:1 mixture of 0.1 M *N*-hydroxy succinimide and 0.1 M 3-(*N*,*N*-dimethylamino)propyl-*N*-ethylcarbodiimide. Unreacted active sites were blocked with a 7 min injection of 1 M ethanolamine at pH 8.0. A second flow cell was not modified and used as a reference channel. Analytes were injected in phosphate-buffered saline (10 mM Na_2_HPO_4_, 1.8 mM KH_2_PO_4_, 137 mM NaCl, 2.7 mM KCl) at pH 7.4 at a flow rate of 50 μL min^−1^. Triplicate injections (in random order) of each sample and a buffer blank were passed over the two surfaces. The sensorgrams were corrected by double referencing. Equilibrium dissociation constants (*K*_D_) were determined by globally fitting the equilibrium response values of the processed sensorgrams to a steady-state affinity model. All data processing and curve fitting was performed with Graphpad Prism v. 8.4.1.

### Microscale Thermophoresis (MST)

Protein–ligand interactions were quantified using microscale thermophoresis (MST). Experiments were performed on a NanoTemper Monolith NT.115 (Nanotemper, München, Germany) instrument. Cyanine 5-maleimide (Lumiprobe, Hannover, Germany) was employed for labeling of CHI3L1 by incubating the protein with a 10-fold molar excess of dye in phosphate-buffered saline at pH 7.4 according to the manufacturer’s recommendations. The excess dye was removed by size exclusion chromatography. A two-fold serial dilution of the unlabeled binding partner (ligand) was prepared in phosphate-buffered saline with the addition of 0.05% Tween-20 at pH 7.4 and incubated with the labeled protein at a final concentration of 40 nM. All samples were incubated for 15-25 min at room temperature to allow binding equilibration before loading into capillaries. For the galectin-3 binding study, labeled CHI3L1 or CBmut CHI3L1 was incubated with serial twofold dilutions of galectin-3 in PBS buffer containing 0.05% Tween-20 in the presence or absence of 500 µM (GlcNAc)_6_, or 500 µM Fondaparinux as additive. The resulting data was analyzed with the instrument software suite or by fitting to a 1:1 binding model based on the law of mass action with Graphpad Prism v. 8.4.1.

### Molecular Dynamics Simulations

The crystal structure of CHI3L1 in the *apo* state (PDB ID: 1HJX) (26) was prepared using the Protein Preparation Workflow (37) (default settings) in Maestro (Schrödinger Release 2024-4: Maestro, Schrödinger, LLC, New York, NY, 2024.). To avoid artificially charged termini, N-and C-terminal residues were capped with ACE and NMA residues, respectively. The p*K*_a_ values and protonation states were predicted at pH 7.4. The predominant protonation state was assigned to each titratable residue. The N-linked glycan, at N60, visible in the crystal structure, was removed during system preparation. The model of the triple mutant was obtained by introducing the substitutions R144A, R145A, and K147A into CHI3L1 using Maestro. 3D models of Fondaparinux and HS9 were generated using the GLYCAM-Web GAG Builder,(38) with HS9 modeled both with and without 6*O*-and 2*O*-sulfation. For each system, one ligand molecule was placed at an arbitrary position using PACKMOL,(39) which resulted in a concentration of ∼1.3 mM. To minimize bias introduced by the starting position of the ligand, we generated 20 independent simulations with different, randomized initial coordinates, in which the initial distance between the ligand and CHI3L1 ranged from 13 Å to 28 Å (see Table S1). The systems were placed in a cubic box and solvated with OPC water,(40) such that the distance between the edge of the box and the closest solute atom was at least 12 Å. To obtain a neutral system, we added explicit Na^+^ and Cl^−^ counter ions, which replaced solvent molecules, with the ionic strength set to 0.15 M, following the SPLIT method.(41) Topology files were built using the tLEaP module in AmberTools24.(42) The ff19SB force field for proteins(43) was used in combination with the GLYCAM06 force field for the carbohydrates (44) and the Li and Merz 12-6 ions parameters for Na^+^ and Cl^−^ ions. (45, 46) For *N*-sulfated glucosamine (GlcNS), the corresponding prep file available in the GLYCAM prep file database was used. Charge adjustments for each *O*-sulfation position were performed according to the documentation on GLYCAM-Web.

All molecular dynamics (MD) simulations were performed with the Amber24 suite of molecular simulation programs (42) using the GPU-accelerated implementation of PMEMD.(47, 48) Covalent bonds involving hydrogen atoms were constrained with the SHAKE algorithm,(49) and application of the hydrogen mass repartitioning strategy(50) allowed the use of a 4 fs integration time step. Periodic boundary conditions were applied in all directions. The Particle Mesh Ewald (PME) method(51) was used to compute long-range electrostatic interactions, while a non-bonded cutoff of 9 Å was employed for both short-range electrostatics and van der Waals interactions. The simulations were prepared following the protocol described in ref.,(52) adapted for the ligand systems investigated here. The systems were initially minimized by relieving unfavorable contacts of the water molecules. Applying harmonic positional restraints with force constants of 2.5-10.0 kcal mol^−1^ Å^−2^ to structural elements of the protein and all ligand atoms (see Table S2), solvent molecules were first minimized for 2,500 steps using the steepest descent algorithm, followed by 2,500 steps of minimization with the conjugate gradient algorithm. Subsequently, the system was heated over 50 ps from 100 K to 300 K in the NVT ensemble. Finally, the system was simulated in the NPT ensemble to adjust the density to 1 g cm^-3^ and to gradually reduce positional restraints (seven steps for a total simulation time of 950 ps; see Table S2). Pressure was maintained at 1 bar using isotropic position scaling with the Berendsen barostat(53) with a pressure relaxation time of 1 ps. Temperature control at a target temperature of *T* = 300 K was maintained using the Langevin thermostat(54) with a collision frequency of γ = 1.0 ps^−1^. 20 independent production runs initialized from different starting positions of the ligand and different initial velocity distributions were performed with a length of 1 µs each. The simulations were not biased by any prior knowledge of potential binding epitopes. This resulted in a total cumulative MD sampling time of 1 µs × 20 runs × 4 ligands = 80 µs. Coordinates were saved in a trajectory file at 100 ps intervals. For the subsequent analysis, snapshots were extracted from the trajectories at regular intervals of 1 ns, resulting in 1,000 configurations per production run.

### Analysis of MD Trajectories

MD trajectories were post-processed and analyzed using CPPTRAJ(55) of AmberTools24 (42) and MDAnalysis (v2.10.0).(56) Trajectories were visually inspected with VMD.(57) All figures showing molecular representations were generated using PyMOL. (58)

We computed the root mean square deviation (RMSD) as a measure of structural variability and the root mean square fluctuation (RMSF) as a measure of mobility for the protein backbone C_α_ atoms relative to the crystal structure (PDB ID: 1HJX) during the MD simulations. The end-to-end distance of the ligands was calculated as the distance from the C1 ring carbon of the reducing end to the C4 ring carbon of the non-reducing end. For calculating the time series for binding of the ligand’s anionic groups to cationic residues of CHI3L1, the distance between the sulfo and carboxyl oxygens and the side chain nitrogens was calculated using the “nativecontacts*”* command as implemented in CPPTRAJ.(55) We employed a cutoff distance of 4 Å to define the formation of a salt bridge between the charged heavy atoms.

Upon visual inspection of the MD trajectories, multiple binding and unbinding events of the ligands to and from CHI3L1 were observed. To visualize preferred binding regions, ligand heavy atom positions from the 20 production runs (20 × 1 µs) were binned onto a grid of 0.5 Å spacing and normalized by the total number of snapshots (20,000) using the “grid” command of CPPTRAJ, yielding spatial occupancy densities of the ligand relative to CHI3L1. For visualization purposes in Figures 4, S20, and S21, we only show regions exceeding the 95^th^ percentile of nonzero grid occupancy values, corresponding to the top 5% of visited grid points across all simulations.

To identify how the ligands interact with CHI3L1, we calculated for each frame if the heavy-atom distance of the ligand to any CHI3L1 residue was below 4 Å and from this constructed a binary contact matrix in which a residue was assigned a value of 1 if the distance criteria was true. Another filtering step was then applied to retain only those frames with five or more interacting residues.(35, 59) Filtered contact fingerprints were clustered using agglomerative hierarchical clustering with average linkage and Jaccard distance as a metric, implemented via SciPy; (60) clusters were defined by applying a Jaccard distance cutoff of 0.6, allowing for transient contact fluctuations between frames of the same binding mode. Subsequently, per-residue contact frequencies were calculated as the proportion of frames within a given cluster in which the residue was in contact with the ligand.

### Adaptive Steered Molecular Dynamics Simulations

To estimate the binding free energy of Fondaparinux and HS9 (6*O*-and 2*O*-sulfated) to CHI3L1, we simulated the unbinding process of the ligands with respect to the binding modes sampled by the fldMD simulations, using adaptive-steered molecular dynamics (ASMD) simulations.(61, 62) In standard steered molecular dynamics (SMD), (63) the steering force is applied through a pseudo particle that is attached to a specified atom (or set of atoms) by a harmonic spring to transverse a reaction coordinate at a particular velocity. The free energy difference between two states A and B (Δ*G* = *G*_B_ − *G*_A_) can be calculated by an appropriately weighted average of the nonequilibrium work *W*_A→B_ over the SMD trajectories using Jarzynski’s equality (eq. 1 and 2): (64)

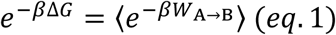

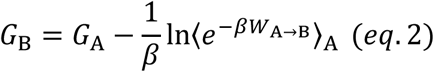

yielding the potential of mean force (PMF) along the selected pathway (*β* = (*k_B_T*)^−1^: *k_B_* and *T* are the Boltzmann constant and temperature, respectively). However, SMD requires many nonequilibrium trajectories to converge the PMF. The ASMD method addresses this limitation by splitting the overall reaction coordinate into several stages with each stage performed as a conventional SMD simulation. At the end of each stage, a contraction of the sampled nonequilibrium configurations is performed to discard trajectories that contribute very little to eq. (2) and, thus, to sample the most important ones more efficiently.(61) We selected the medoids of the two most populated clusters (c1 and c2) as starting conformations for the nonequilibrium pulling simulations. Each system was prepared for the ASMD simulations following the minimization and equilibration protocol described previously (see Table S1), whereas in the last density adaptation step, the distance between the heavy-atom center of mass (COM) of the ligand and that of the protein was restrained with a harmonic force constant of 100 kcal mol^−1^ Å^−2^. (65, 66) ASMD simulations were performed to pull the ligands away from the protein,(67, 68) increasing the distance between the COMs by 30 Å, using the aforementioned force constant and a constant pulling velocity of *v* = 1.0 Å ns^−1^. All ASMD simulations were partitioned into 15 stages, and for each stage 48 simulations with different randomized initial velocities were simulated for 2 ns, such that the distance was perturbed 2 Å/stage. After completion of each pulling stage, the coordinates at the end of the trajectory whose nonequilibrium work is the closest to the Jarzynski average served as the starting point for the next pulling stage. ASMD has been shown to significantly reduce the number of trajectories necessary for convergence of the PMF compared to conventional SMD simulations, hence lowering the computational cost.(62)

To estimate the statistical uncertainty of the pulling simulations, we calculated the Jarzynski-weighted cumulative error at the end of each stage according to ref.(69) For a given stage (*n*) consisting of *N* trajectories, the Jarzynski-weighted averages of the work (⟨*w*⟩*_JA_*) (eq. 3) and the squared work (⟨*w*^2^⟩*_JA_*) (eq. 4) were computed using the exponential Boltzmann factor as a statistical weight:

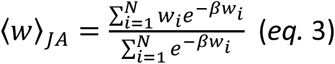

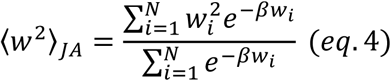

where *w_i_* denotes work of the trajectory *i* during the stage. These weighted averages were used to define the statistical variance (σ_n_ ^2^) of a stage *n* (eq. 5):

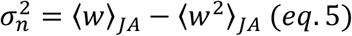

To account for the accumulation of uncertainty over the stages, the variances of all preceding stages were summed. The final reported error (*E*) was calculated as the square root of this cumulative variance (eq. 6):

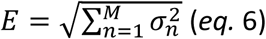

where *M* is the number of stages at a given point along the reaction path.

### Fluorogenic Chitinase Activity and Inhibition Assays

A dilution series of 4-Methylumbelliferyl β-D-*N*,*N*ʹ,*N*ʹʹ-triacetylchitotrioside was added to a solution containing 70 nM of a catalytically active CHI3L1 mutant (Cat CHI3L1) in PBS buffer in the presence or absence of 1.5 mg mL^−1^ UFH. Reaction kinetics were continuously tracked over 120 minutes to capture the progress curves by measuring fluorescence intensity of the reaction product 4-Methylumbelliferone at an excitation wavelength of 360 nm and an emission wavelength of 450 nm. Raw relative fluorescence units (RFU) were corrected by subtracting the background signal from substrate-blank control wells. The initial velocity values were calculated from the slope of a regression line fitted to the linear phase of the progress curve (*t* = 0 s to 2000 s) for each condition. A plot of the resulting initial velocities against substrate concentration was analyzed by fitting to a Michaelis-Menten model.

## Results

### The Affinity of CHI3L1 to GAG or COS Ligands Is Dependent on the Degree of Polymerization and Is Differentially Affected by Mutations in the GAG and Chitin Binding Sites

To dissect the influence of DP on the interactions with carbohydrate-based ligands, the affinities of a series of COS (DP4-6) and polydisperse (Enoxaparin (DP ca. 2-24), UFH (DP ca. 40-60)), as well as monodisperse (Fondaparinux (DP5); heparan-sulfate nonamer HS9 (DP9)) GAG preparations were determined. SPR revealed a progressively increasing affinity for COS ligands with increasing DP, ranging from 83.5 µM for (GlcNAc)_4_ to 2.8 µM for (GlcNAc)_6_ (Table 1). We observed a pronounced ca. 7-fold improvement in affinity upon elongation of the chitin chain from (GlcNAc)_4_ to (GlcNAc)_5_ and a further ca. 4-fold affinity enhancement for (GlcNAc)_6_. This indicates the progressive occupancy of six canonical subsites within the CHI3L1 binding cleft, with each additional GlcNAc unit providing a significant contribution to the binding energy.

**Table 1.** Binding affinities of COS and GAG ligands of different DP to CHI3L1 and its mutant variants. Raw data is displayed in Figures S1 and S2.

| Ligand | Protein | $K_D$ [μM] | Method |
| --- | --- | --- | --- |
| (GlcNAc)<sub>4</sub> | CHI3L1-Wild type | 83.5 (70.5-98.9) | SPR |
| (GlcNAc)<sub>5</sub> | CHI3L1-Wild type | 11.9 (11.0-12.9) | SPR |
| (GlcNAc)<sub>6</sub> | CHI3L1-Wild type | 2.8 (2.5-3.1) | SPR |
| (GlcNAc)<sub>6</sub> | CBmut CHI3L1 | No binding detected | SPR |
| Fondaparinux | CHI3L1-Wild type | 172.4 (76.4-369.1) | MST |
| Fondaparinux | HepBmut CHI3L1 | No binding detected | MST |
| HS9 | CHI3L1-Wild type | 6.7 (4.6-9.7) | MST |
| HS9 | HepBmut CHI3L1 | No binding detected | MST |
| Enoxaparin | CHI3L1-Wild type | 5.0 (2.1-12.0) | MST |
| UFH | CHI3L1-Wild type | 5.2 (3.2-9.8) | SPR |
| UFH | CBmut CHI3L1 | 5.5 (3.6-7.4) | SPR |

GAG derivatives display a similar dependence on DP. Whereas the chemically defined pentasaccharide Fondaparinux only showed a weak affinity (*K*_D_ = 172.4 µM by MST), an approximately 26-fold increase in affinity was observed for HS9 (*K*_D_ = 6.7 µM). UFH yielded an approximately 33-fold increase in affinity compared with Fondaparinux. Note that the interaction with UFH likely involves marked contributions of multivalency and stoichiometries differing from a classical 1:1 interaction. A direct comparison with shorter monodisperse GAG ligands should, thus, be viewed with caution. To account for this fact, the affinity was characterized by SPR employing a heparin-coated surface in combination with a Hill binding model, which yielded a Hill coefficient of ca. *n* = 2, potentially indicating a deviation from 1:1 stoichiometry. Notably, the observed affinity values for GAG ligands of different DP show a steep increase from DP5 to DP9 and plateau for longer GAG ligands, indicating that a further increase in chain length does not meaningfully contribute to the interaction, potentially discounting primarily entropic contributions from multivalency. This finding was corroborated by a glycan array analysis, in which defined GAG fragments of different DP were probed for their affinity to CHI3L1 (Figure 2).(31) In this experiment, a steep increase of fluorescence intensity, indicating binding of labeled CHI3L1, was recorded between DP9 and DP12. The high affinity of HS9 in MST experiments might, therefore, indicate a contributing role of the diverging sulfation pattern to CHI3L1 affinity. Overall, the general picture that emerges from both experiments is that GAG affinity for the DPs investigated reveals two states with a steep transition between low (ca. DP 5-10) and high (ca. DP > 10) DP.

**Figure 2.**
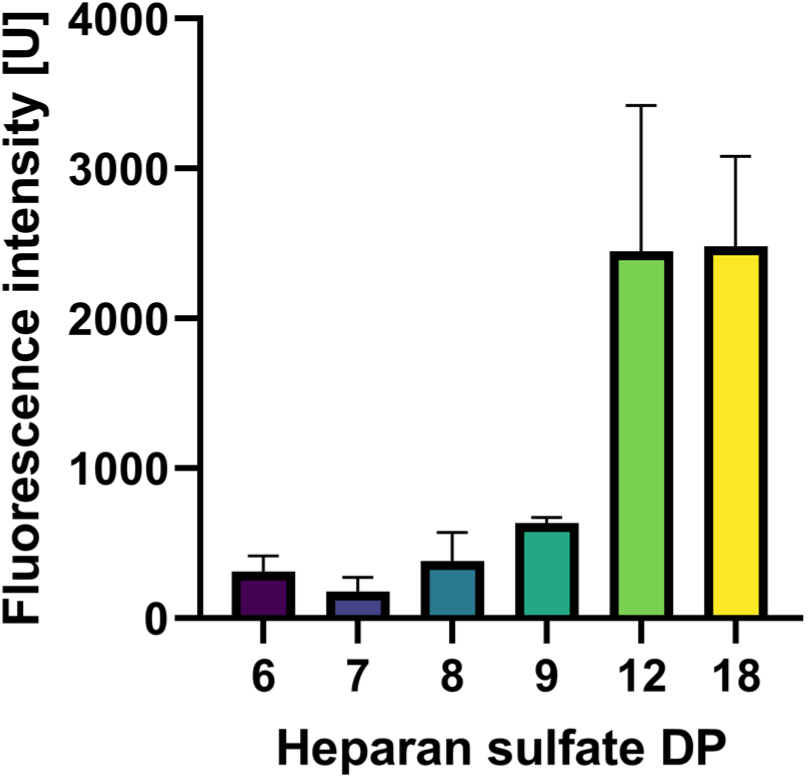
CHI3L1 binding to heparan sulfate fragments of different DPs. The fluorescence intensity of fluorescently labeled CHI3L1 bound to the glass surface increases as a function of DP.

To differentiate the binding sites for both ligand classes, two protein constructs containing a mutation in either the canonical GAG binding site (HepBmut CHI3L1: R144A; R145A; K147A) or the chitin binding site (CBmut CHI3L1: W31A; W99A; W352A) were produced and their affinity towards GAG and COS ligands was determined (Table 1). Whereas COS binding affinity of CBmut CHI3L1 was fully abrogated, the mutant protein fully retained its ability to bind to a heparin-modified surface in SPR experiments. Conversely, no binding affinity of HepBmut CHI3L1 to GAG ligands could be determined by MST in the measured concentration range.

### The Fondaparinux Binding Mode is Conserved Independently of the Canonical Chitin-Binding Site

The interaction of Fondaparinux with CHI3L1 was further investigated using Saturation Transfer Difference (STD) NMR spectroscopy to map the ligand-binding epitope (Figure 3). The ^1^H NMR spectrum of Fondaparinux was assigned (reference spectra at the top in Figure 3A, red) by using previously published data from Hricovíni et al..(34) In the presence of wild type (WT) CHI3L1, strong STD signals were observed for multiple residues (blue spectrum in Figure 3A), with the highest relative intensity detected for the protons colored in blue (≥100%) in the chemical structure of Fondaparinux (Figure 3C), indicating their close proximity to the protein surface. Additional contributions from protons within the central saccharide units further suggest an extended binding interface involving several regions of the ligand (yellow in Figure 3C, 76%-99%). Protons present in the terminal saccharides showed weak STD signals (green in Fig 3C, ≤75%), indicating that these residues are located further away from the binding interface. In the CBmut CHI3L1 variant, which carries mutations in the chitin-binding site, a comparable STD pattern was observed, with similar relative intensities across the Fondaparinux structure (Figure 3B, D). Epitope mapping projected onto the molecular structure revealed that the same regions of the ligand are predominantly involved in binding to both protein variants. Notably, the distribution of high-intensity STD signals remained largely unchanged, indicating preservation of the binding mode.

**Figure 3.**
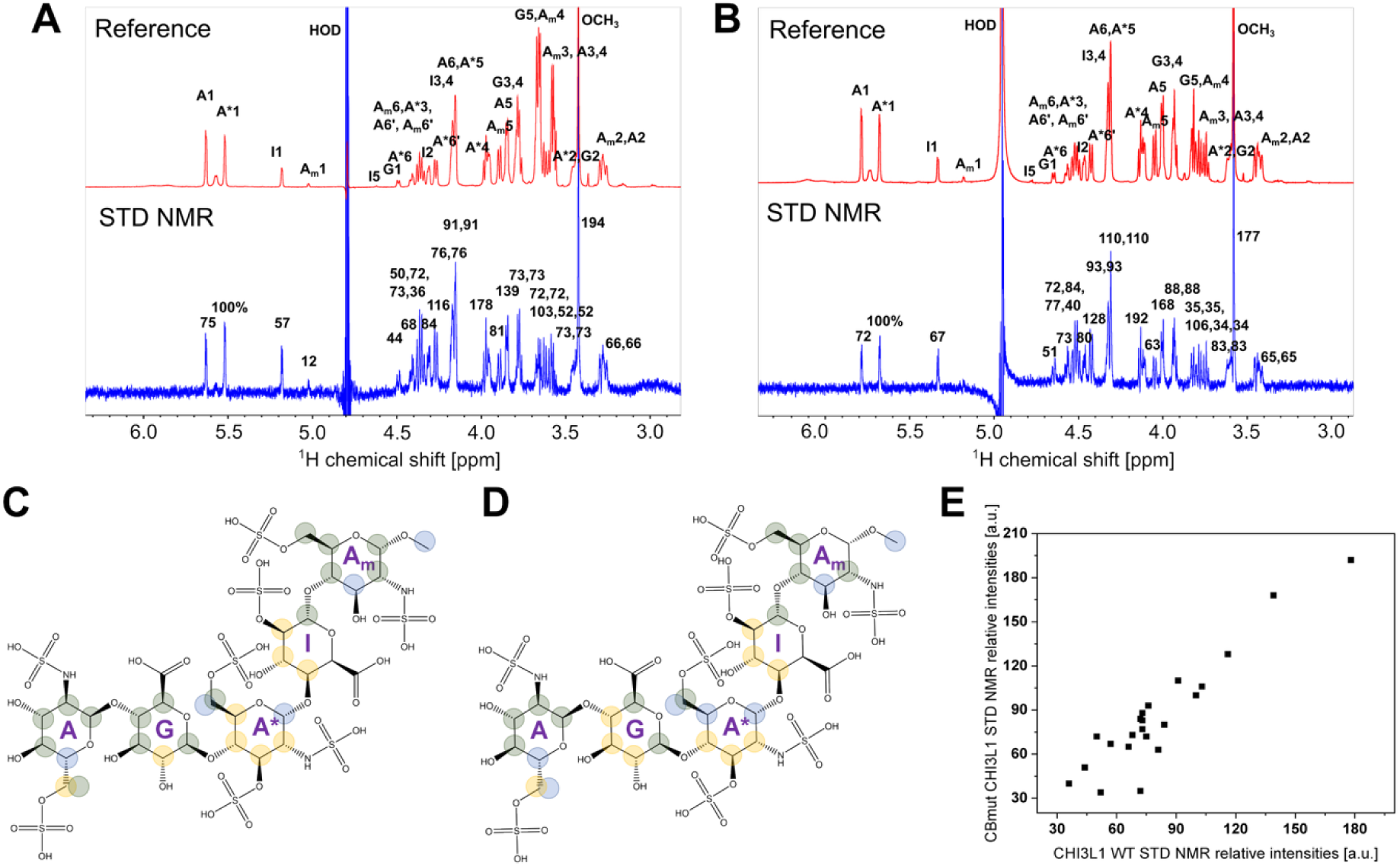
Mapping of the CHI3L1 binding epitope of Fondaparinux by STD NMR. (A, B) ^1^H NMR reference spectra (red, top) and Saturation Transfer Difference (STD) NMR spectra (blue, bottom) of Fondaparinux in the presence of WT (A) and CBmut CHI3L1 (B). The reference spectra show the assignment of proton resonances for the pentasaccharide. (C, D) Binding epitope mapping projected onto the chemical structure of Fondaparinux, associated with panels A and B, respectively. The relative STD intensities are indicated by colored circles, where the STD intensity of the A*1 proton was set to 100% as a reference. Similar and higher percentages (yellow: 76%-96% or blue: 100%-200% circles) signify closer proximity to the protein surface, while lower percentages (green: 35%-75%) indicate residues further away from the binding interface. (E) Correlation plot of STD intensities of Fondaparinux in the presence of WT (A) and CBmut CHI3L1 (B), indicating that mutation of the chitin binding site does not significantly affect heparin binding.

Quantitative comparison of STD intensities WT and CBmut CHI3L1 demonstrated a strong correlation (Figure 3E), supporting the conclusion that mutation of the chitin-binding site does not significantly alter Fondaparinux binding. These findings suggest that Fondaparinux interacts with CHI3L1 through a binding site distinct from the canonical chitin-binding domain, highlighting an alternative interaction interface relevant for heparin/HS recognition.

### R144, R145, and K147 Form the Primary Heparin-Binding Site in CHI3L1 and Support Multivalent Engagement of Longer Glycosaminoglycans

To investigate the molecular mechanism of Fondaparinux and heparan sulfate DP9 (HS9) binding to CHI3L1, we performed unbiased molecular dynamics (MD) simulations of free ligand diffusion (fldMD).(35, 70, 71) In these simulations, no artificial guiding or biasing force was applied to any of the molecules, allowing for the characterization of spontaneous association and dissociation events between the protein and the ligand at atomistic resolution. We performed 20 independent simulations for both Fondaparinux and HS9 (with simulation times of 1 µs each), in which one ligand molecule was arbitrarily placed in the water phase of the simulation box; this corresponds to a ligand concentration of ca. 1.3 mM. During the simulations, CHI3L1 remained structurally invariant, as demonstrated by the time courses of the root-mean-square deviation (RMSD) of the C_α_ atomic positions (Figure S3-S6). In most of the simulations, both ligands diffused extensively before interacting with CHI3L1 (Figure S8 and S9). To characterize the conformational ensemble of the ligands in solution, we calculated the end-to-end distance (*R*_ee_), which revealed a preference for the extended conformation (Figure S12 and S13),(72, 73) in line with visual observations (Figure S8 and S9). During the fldMD simulations, we observed multiple binding events of Fondaparinux and HS9 to positively charged surface locations of CHI3L1 (Figure 4A, B, S7). A binding event was recorded if the heavy-atom distance between ligand and protein was less than 4 Å, and interactions with at least five residues were formed (Table S1).(35, 59) Subsequently, ligand poses were clustered using fingerprints of interactions with CHI3L1 residues to resolve distinct binding sites (Figure S16 and S17). To qualitatively visualize the binding regions identified by the cluster analysis, we calculated occupancy densities of Fondaparinux/HS9 and mapped them around the structure of CHI3L1 (Figure 4B, D).(74)

**Figure 4.**
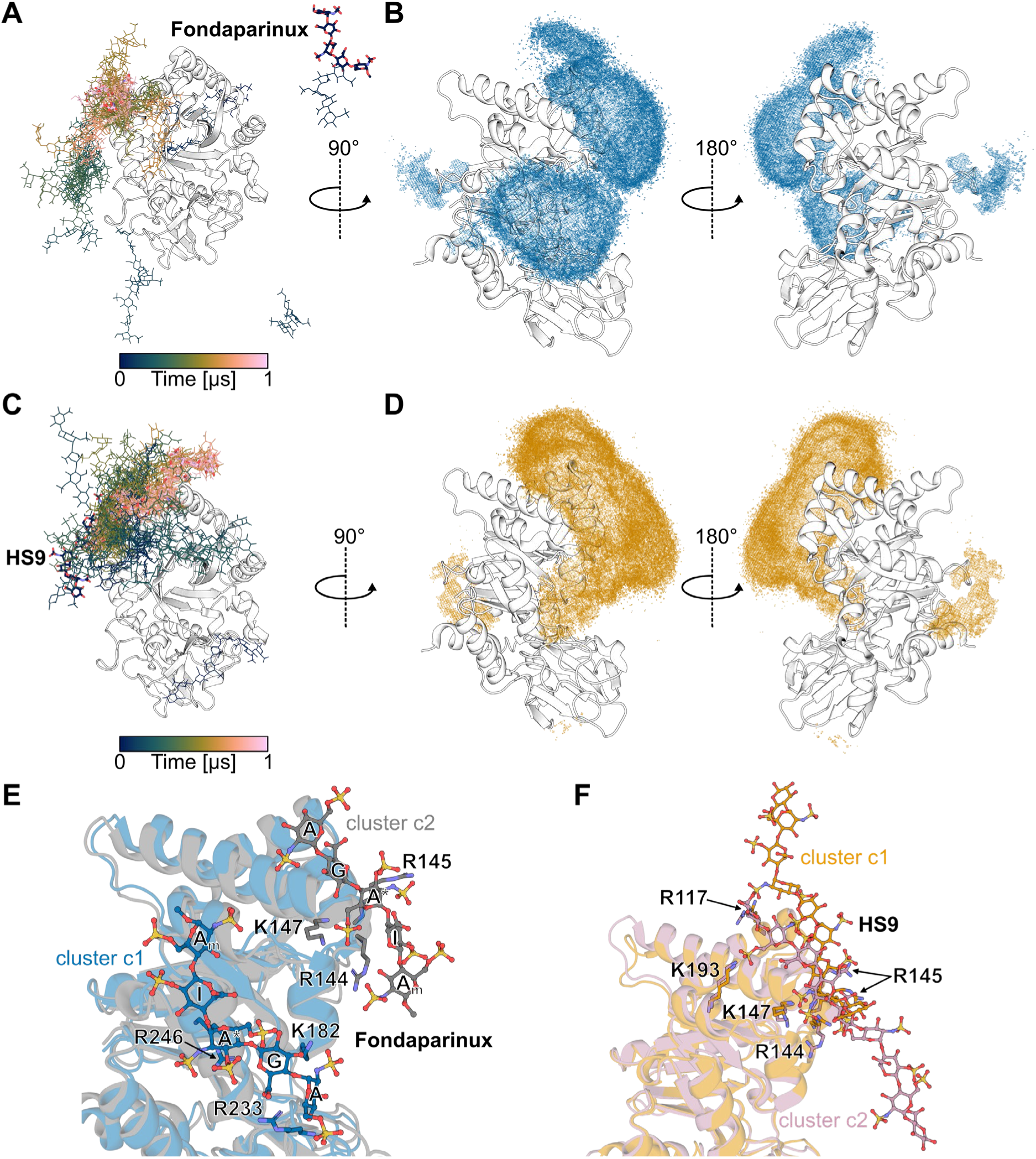
Unbiased MD simulations of CHI3L1-Fondaparinux/HS9 binding. (A-D) The Fondaparinux (A) and HS9 (C) molecules moving from the free state (stick representation) to the bound state (lines representation) in selected fldMD simulations. The derived occupancy densities from all fldMD simulations for Fondaparinux (B; blue mesh) and HS9 (D; orange mesh) are mapped around the CHI3L1 crystal structure (white cartoon; PDB ID: 1HJW). For the occupancy density contour level, see the Materials and Methods section. The orientation of the protein in the left structures is identical to the one in Figure 1A, right. (E) Predominant CHI3L1/Fondaparinux binding pose (cluster medoid; ball-and-stick model) from the c1 (blue) and c2 (grey) cluster (same orientation as the left structure in B), with amino acids (stick representation) binding Fondaparinux. The sugar moieties of Fondaparinux are labeled according to ref.(34) (F) Predominant CHI3L1/HS9 binding pose (cluster medoid; ball-and-stick model) from the c1 (orange) and c2 (violet) cluster (same orientation as the left structure in D), with amino acids (stick representation) binding HS9.

Inspection of the Fondaparinux densities revealed that the ligand primarily interacted with CHI3L1 at or adjacent to two regions: (1) a cluster of positively charged residues (K182, R233, R246) located near the reducing end of the carbohydrate-binding groove between the interface of helices 5 and 6 (cluster 1 (c1), population 35.0%); and (2) the previously described (27, 28, 30) heparin-binding site ^143^GRRDKQ^148^, following the consensus sequence [-X-B-B-X-B-X-],(75) where B is a basic and X is a nonbasic residue (cluster 2 (c2), population 20.1%) (Figure 4B and S16). Intriguingly, in some simulations, we observed that Fondaparinux initially bound to this nonspecific secondary heparin-binding site, then moved and rearranged to the consensus binding region while remaining associated with the protein surface, and vice versa (Figure 4A and S8). Rather than adopting a single defined conformation, Fondaparinux interacts in a highly dynamic fashion, forming transient salt bridges between its sulfate or carboxylate groups and arginine and lysine side chains (Figure 5A). This is further supported by the time-series data, which show that the anionic groups of Fondaparinux bind intermittently to the previously described interacting cationic residues of CHI3L1 (Figure S26).

**Figure 5.**
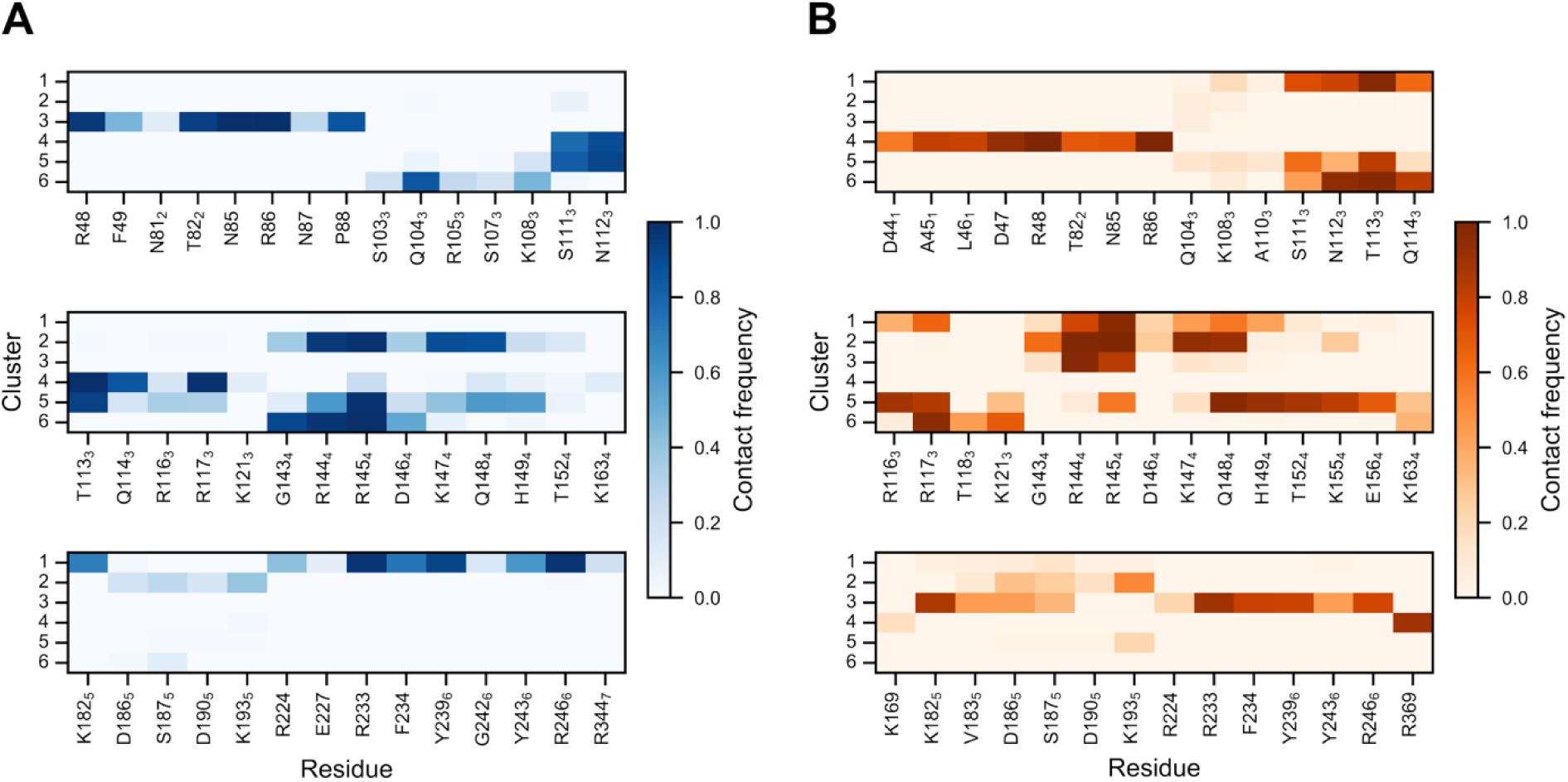
Contacts between CHI3L1 and Fondaparinux/HS9. (A, B) Heatmap illustrating the residue-wise contact frequencies between CHI3L1 with Fondaparinux (A) and HS9 (B). Data are shown for the six most populated clusters, ranked by size (1 = highest population) (see also Figures S16 and S17). Only residues exhibiting a contact frequency >10% in at least one cluster are shown. Subscript numbers indicate the α-helix (see Figure 1A) to which the residue belongs; the absence of subscripts indicates a residue location in a loop.

The STD NMR experiments revealed that each sugar residue of Fondaparinux is in contact with CHI3L1, while the highest STD percentages were observed for the protons of the central A* residue (Figure 3), indicating it as the interacting core. The representative conformations, identified as the cluster’s medoid, of clusters c1 and c2 show binding modes that agree with the observed STD signals, i.e., the central sugar A* is preferentially in contact with CHI3L1 (see Figure 4E). Due to the observed fluctuations in the binding mode with attachment and detachment events, pointing towards an entropically-dominated binding mode,(76, 77) the terminal units A and A_m_ are frequently solvent-exposed (Figure S26), which could account for their weaker STD signals. Taken together, these results indicate predominantly electrostatic, low-specificity interactions, consistent with the low affinity observed experimentally (Table 1) and aligning with previous observations.(28)

Unlike Fondaparinux, HS9 exhibited its highest occupancy density around the ^143^GRRDKQ^148^ heparin-binding site (Figure 4D). In the observed binding modes, part of the HS9 chain was anchored to residues R144, R145, and K147, while the other units of the chain formed extended interactions with adjacent basic residues, engaging R116 and R117 (cluster 1 (c1), population 28.6%) on one side, or K155 and K193 (cluster 2 (c2), population 21.3%) on the other, or simultaneously engaging both the consensus and nonspecific site sampled by Fondaparinux (cluster 3 (c3), population 9.4%) (Figure 5B and S17). Obviously, the increased chain length allows HS9 to form extended interactions, thereby effectively bridging distinct cationic patches, stabilizing the ligand on the surface of CHI3L1. This is further supported by the time-series data, which show that charged groups of HS9 stay bound significantly longer to individual protein residues than those of Fondaparinux (Figure S27). In addition, to investigate the influence of the degree of sulfation of HS9 (see Figure 1B) binding to CHI3L1, we performed fldMD simulations with HS9 lacking 2*O*-and 6*O*-sulfate groups (HS9_Δ2S/6S_) (Figure S10). Interestingly, HS9_Δ2S/6S_ showed only a slight reduction in the number of binding events as the 2*O*-and 6*O*-sulfated HS9 molecule (Table S3) and interacted mainly with the ^143^GRRDKQ^148^ binding site (Figure S20 and S22). This shows a high specificity of different HS molecules for the ^143^GRRDKQ^148^ binding site and highlights important contributions of polar residues located within helices 3 and 4.

To validate the prediction that the ^143^GRRDKQ^148^ motif is the preferred heparin-binding region for GAG anchoring, we generated a CHI3L1 triple mutant (HepBmut) in which the three basic interface residues R144, R145, and K147 were mutated to alanine. Comparative fldMD simulations were performed to reassess binding of the high-affinity ligand HS9 to the mutant variant, following the same protocol used for the WT (Figure S11). The introduction of these mutations neither compromised the overall structural stability of the protein nor induced significant conformational changes, as suggested by the C_α_ root-mean-square deviation and fluctuation profiles (Figure S6, S24). In these simulations, HS9 interacted in a nonspecific and transient manner mainly at two distinct regions of the protein (Figure S21). Specifically, occurrences of HS9 were localized to the previously described sampled heparin-binding site (cluster 1 (c1), population 22.8%) and the interface between helices 3 and 4, where it engaged residues R116, R117, and K155 (cluster 2 (c2), population 21.5%) (Figure S19 and S23). Relative to the WT, the mutant exhibited a pronounced 41.7% decrease in the number of binding events (Table S3). While our MST measurements demonstrated a complete abrogation of binding affinity for this mutant, we still detected residual binding during the simulations, likely because the HS9 concentration in the simulation box (∼1.3 mM) was higher than under experimental conditions. This observation is corroborated by single molecule force spectroscopy (SMFS) measurements, in which two independent unbinding events of different tensile strengths are recorded for the CHI3L1/HS interaction.(31)

Taken together, these results suggest that R144, R145, and K147 form the primary heparin-binding site in CHI3L1. As the degree of polymerization increases, longer heparin/HS molecules can engage in simultaneous, multivalent interactions with adjacent basic patches and polar networks, contributing to the overall stability of the complex. Notably, in contrast to previous reports, we did not observe any binding for either Fondaparinux or HS9/HS9_Δ2S/6S_ at the KR-rich domain(29) or within the chitin-binding cleft.(25, 29)

### Fondaparinux Preferentially Binds to the ^143^GRRDKQ^148^ Site and HS9 Shows Distinct Unbinding Thermodynamics

To complement our findings from the structural analysis with a view on the energetic preference of binding, we conducted adaptive steered MD (ASMD) simulations (61, 62) to estimate energetic binding properties of the two ligands with respect to the sampled binding modes/sites. For each ligand, we selected the medoids of the two most populated clusters (c1 and c2) as starting conformations (Figure 6A, B). By employing a nonphysical spring to apply an external force, the ligand was pulled from the bound state identified in the fldMD simulations towards the unbound state in the bulk solvent across several stages.(65, 67, 78) The nonequilibrium work performed on the system was determined by integrating the force of the spring over each pulling trajectory within these stages. By applying Jarzynski’s equality (64) to the resulting ensemble of ASMD trajectories, this work was rigorously related to the free energy change along the reaction coordinate, yielding a unidimensional potential of mean force (PMF; Figure 6C, D). Differentiating the PMF with respect to distance yielded the computed pulling forces for the unbinding process (Figure 7), enabling a qualitative comparison with recent experimental SMFS measurements.(31)

**Figure 6.**
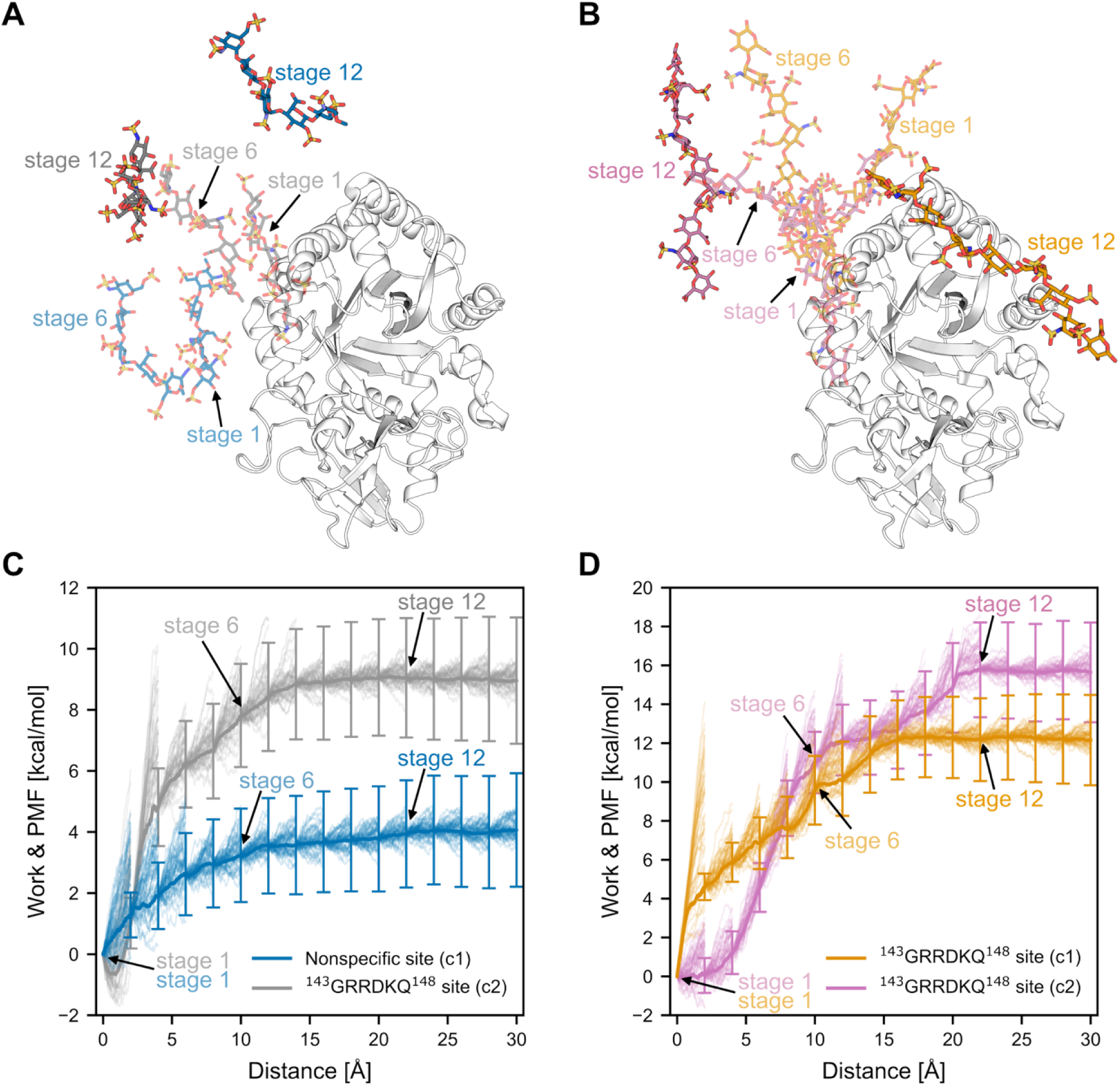
ASMD-derived PMFs for the unbinding of Fondaparinux and HS9 from CHI3L1. (A, B) Fondaparinux (A) and HS9 (B) are pulled from their initial bound conformations (cluster medoids; see Figure 4E, F; d = 0 Å) toward the fully solvated state (d = 30 Å). Snapshots depict the starting configuration (stage 1; translucent), an intermediate (stage 6; translucent), and the detached ligand (stage 12; opaque); corresponding stages are indicated in the PMF profiles. For Fondaparinux, blue (c1) represents unbinding from the nonspecific site, while gray (c2) represents the ^143^GRRDKQ^148^ heparin-binding site. For HS9, orange (c1) and violet (c2) represent different starting conformations anchored to the ^143^GRRDKQ^148^ site. (C, D) Jarzynski-averaged PMF profiles for Fondaparinux (C) and HS9 (D), respectively. Thick lines represent the PMF, while thin lines show the 48 individual work traces per stage. Error bars indicate the cumulative Jarzynski-weighted standard deviation at the end of each ASMD stage.(69)

**Figure 7.**
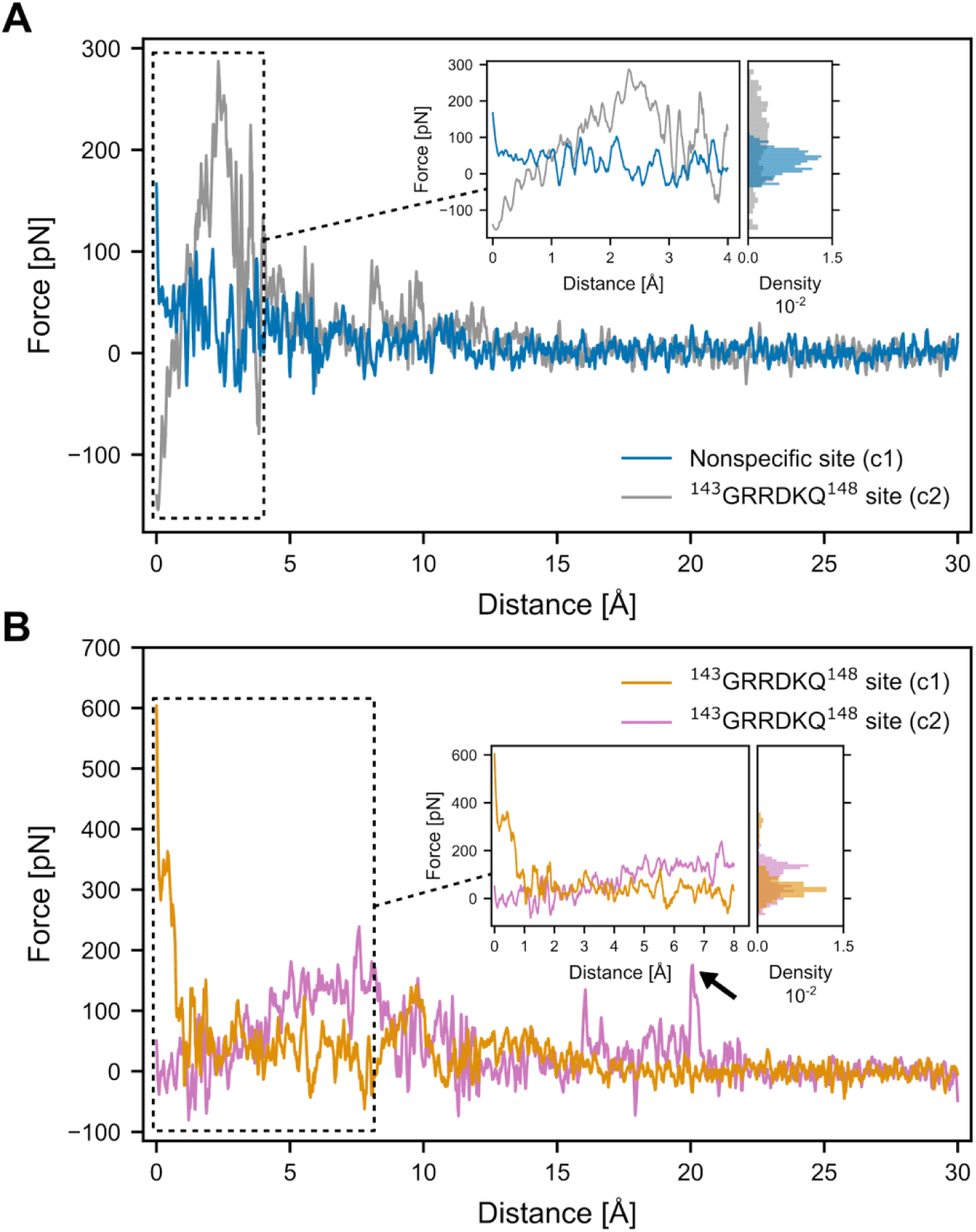
Forces of the unbinding process of Fondaparinux and HS9 from CHI3L1. (A, B) Unbinding force profiles for Fondaparinux (A) and HS9 (B), calculated as the first-order derivative of the respective PMFs. The data were smoothed with a Savitzky-Golay filter (window length = 51, degree of the smoothing polynomial = 2). The black arrow in panel B indicates the observed residual interactions. Magnification of the early unbinding phase detailing the initial force peaks for Fondaparinux (A; d = 0-4 Å) and HS9 (B; d = 0-8 Å) is shown in the insets. Color coding is the same as in Figure 4: for Fondaparinux, blue (cluster c1) represents unbinding from the nonspecific site and gray (cluster c2) from the ^143^GRRDKQ^148^ heparin-binding site; for HS9, orange (cluster c1) and violet (cluster c2) denote different starting conformations anchored at the ^143^GRRDKQ^148^ site.

The resulting Jarzynski-averaged PMF profiles for the unbinding process of Fondaparinux reveal a difference of ∼5 kcal mol^−1^ in the free energy required to pull the ligand from the nonspecific (Δ*G*_unbinding, c1_ = 4.1 ± 1.9 kcal mol^−1^) and the ^143^GRRDKQ^148^ heparin-binding site (Δ*G*_unbinding, c2_ = 9.0 ± 2.1 kcal mol^−1^) (Figure 6C). Aside from the higher energy barrier required to extract the ligand from the ^143^GRRDKQ^148^ site, which indicates a stronger binding affinity, we observe that both PMF profiles reach a plateau after complete dissociation of the pentasaccharide (at a distance of ∼13 Å).

During the forced unbinding of HS9, the cluster c1 binding mode requires Δ*G*_unbinding, c1_ = 12.2 ± 2.3 kcal mol^−1^ for complete dissociation, with the system reaching a plateau at ∼16 Å. Although the PMF profile for the cluster c2 binding mode initially follows a similar trend, the free energy for cluster c2 continues to rise beyond a pulling distance of 16 Å (Figure 6D). We attribute this secondary energetic hurdle to persistent contacts between a terminal sugar and CHI3L1 residues. Breaking these residual interactions extends the unbinding coordinate to ∼22 Å, resulting in a higher dissociation free energy of Δ*G*_unbinding, c1_ = 15.6 ± 2.6 kcal mol^−1^.

Overall, the relative trends and order of magnitude in the calculated free energy profiles for Fondaparinux and HS9 are in good agreement with the experimental values (Table 1). However, the absolute unbinding energies are likely overestimated as a consequence of nonequilibrium pulling, which, especially in the case of HS9, restricts the long and highly flexible GAG chain to sufficiently sample its conformational and configurational degrees of freedom in the unbound state.

The force-distance profiles for Fondaparinux reveal that during the initial stages of extraction (0 to 4 Å), the pulling forces required to unbind the ligand from the ^143^GRRDKQ^148^ site are higher than those for the nonspecific site (Figure 7A). This observation is in line with the steeper initial gradient of the respective PMF profile (Figure 6C). Conversely, upon reaching the plateau region at distances >13 Å, the pulling force fluctuates around zero, confirming that Fondaparinux is completely unbound. For HS9, analysis of the unbinding process reveals distinct mechanical responses depending on the engagement of additional contacts while still bound at the ^143^GRRDKQ^148^ site (Figure 7B). The cluster c1 binding mode exhibits maximum resistance immediately at the onset of pulling (0 to 2 Å), whereas the rupture force for cluster c2 steadily builds, reaching its peak just before 8 Å (Figure 7A). Notably, despite the different timescales inherent to ASMD and experimental setups,(79) the magnitudes of the reproduced force fluctuations agree well with recent SMFS measurements.(31)

To conclude, the ASMD simulations reveal a stronger binding of Fondaparinux to the specific ^143^GRRDKQ^148^ site compared to the nonspecific secondary site, and highlight distinct, conformation-dependent unbinding thermodynamics for HS9.

### Differential Thermal Stability and Tryptophan Fluorescence Signatures Support Distinct GAG and Chitin Binding Sites in CHI3L1

To explore how ligand binding affects protein stability, we examined the thermal unfolding profile of CHI3L1 in the presence of GAGs and COS utilizing nanoDSF (Figure 8). The structurally defined GAG fragment Fondaparinux produced a Δ*T*_m_ of −0.3 °C at a concentration of 1 mM. Similarly, UFH produced a Δ*T*m of −0.2 °C at a concentration of 2.5 mg/mL. Contrastingly, COS stabilized the protein, with (GlcNAc)_6_ yielding a positive Δ*T*_m_ of 3.5 °C at 50 µM concentration. In addition, it can be observed that COS binding results in a marked shift of the ratio of fluorescence at 350 nm and 330 nm. This is in line with crystallographic observations that COS engage in interactions between the apolar α-faces of the GlcNAc pyranose rings and Trp side chains, which strongly contribute to the observed fluorescence signal. The fact that a similar effect is not observed for the interaction with GAGs suggests that these ligands interact via a different binding mode or site, not involving aromatic residues. Thus, it can be assumed that the local environment of Trp sidechains is unperturbed upon GAG binding and that this class of ligands is unlikely to bind, even partially, at the Trp-rich chitin binding site. Together, these results support the hypothesis that heparin and chitin interact with CHI3L1 through distinct and functionally non-overlapping binding sites, thereby inducing opposite effects on protein stability.

**Figure 8.**
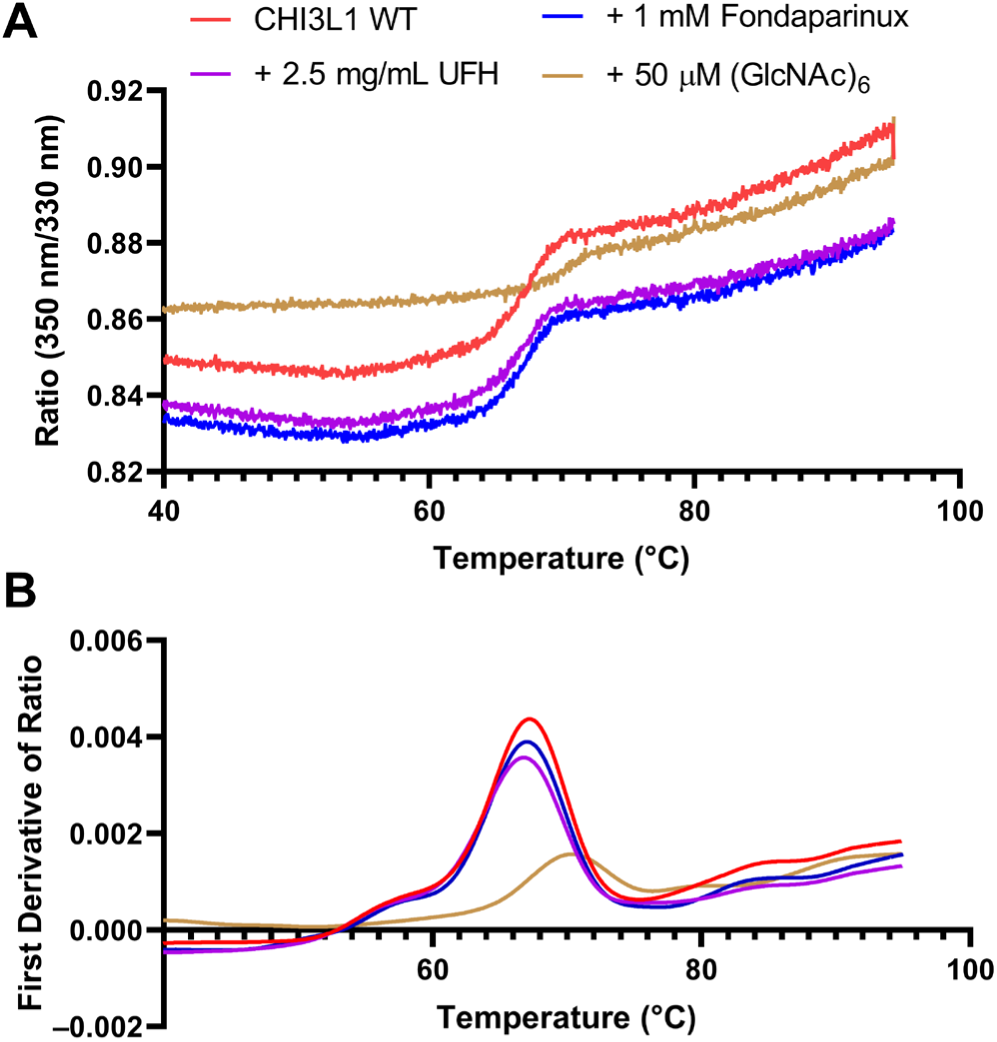
Thermal unfolding profile of CHI3L1 by nanoDSF. (A) Thermal denaturation of CHI3L1 WT was recorded without ligand (red) or in the presence of 1 mM Fondaparinux (blue), 2.5 mg/mL UFH (purple) or 50 µM (GlcNAc)_6_ (brown) by observing the ratio of intrinsic protein fluorescence at 350 nm and 330 nm. (B) First derivative calculated from data in (A).

### COS and GAG Ligand Binding to CHI3L1 is Not Competitive

To investigate a potential interaction between the binding sites of COS and GAGs, binding assays were performed in a competitive assay setup via SPR. Fondaparinux was employed as the competitor rather than UFH, because of the observation of a persistent, non-recovering increase in RU due to quasi-irreversible binding of the multivalent ligand to the sensor-chip surface. Our results revealed a *K*_D_ of 2.9 µM (2.8 to 3.1 µM) for (GlcNAc)_6_ in the presence of 250 µM Fondaparinux, which is virtually identical to the affinity of (GlcNAc)_6_ alone (*K*D = 2.8 µM, Table 1, Figure 9A). As an orthogonal method to delineate potential competition between COS and GAG ligands, an enzymatic assay employing the reactivated catalytically active mutant Cat CHI3L1 was selected for this effort. This approach was expected to distinguish between a potential allosteric or orthosteric mode of inhibition with an enzyme kinetic analysis. For this, the production of fluorescent 4-methylumbelliferone from the chitinase substrate 4-methylumbelliferyl β-D-*N*,*N*ʹ,*N*ʹʹ-triacetylchitotrioside was recorded in the absence or presence of 1.5 mg mL^−1^ UFH (Figure 9A). Contrary to the initial hypothesis, however, no significant difference could be detected between both tested conditions in the progress curves. Fitting the resulting initial velocity values for all tested concentrations yielded superimposable Michaelis-Menten curves with non-distinguishable kinetic parameters (Figure 9B). Instead of the expected effects of inhibition by heparin that would have allowed a differentiation between allosteric or orthosteric inhibition, the observed curves indicate that heparin does not modulate the catalytic activity of Cat CHI3L1. Consequently, it can be concluded that the COS and GAG binding sites of CHI3L1 do not overlap and are not functionally connected through allostery.

**Figure 9.**
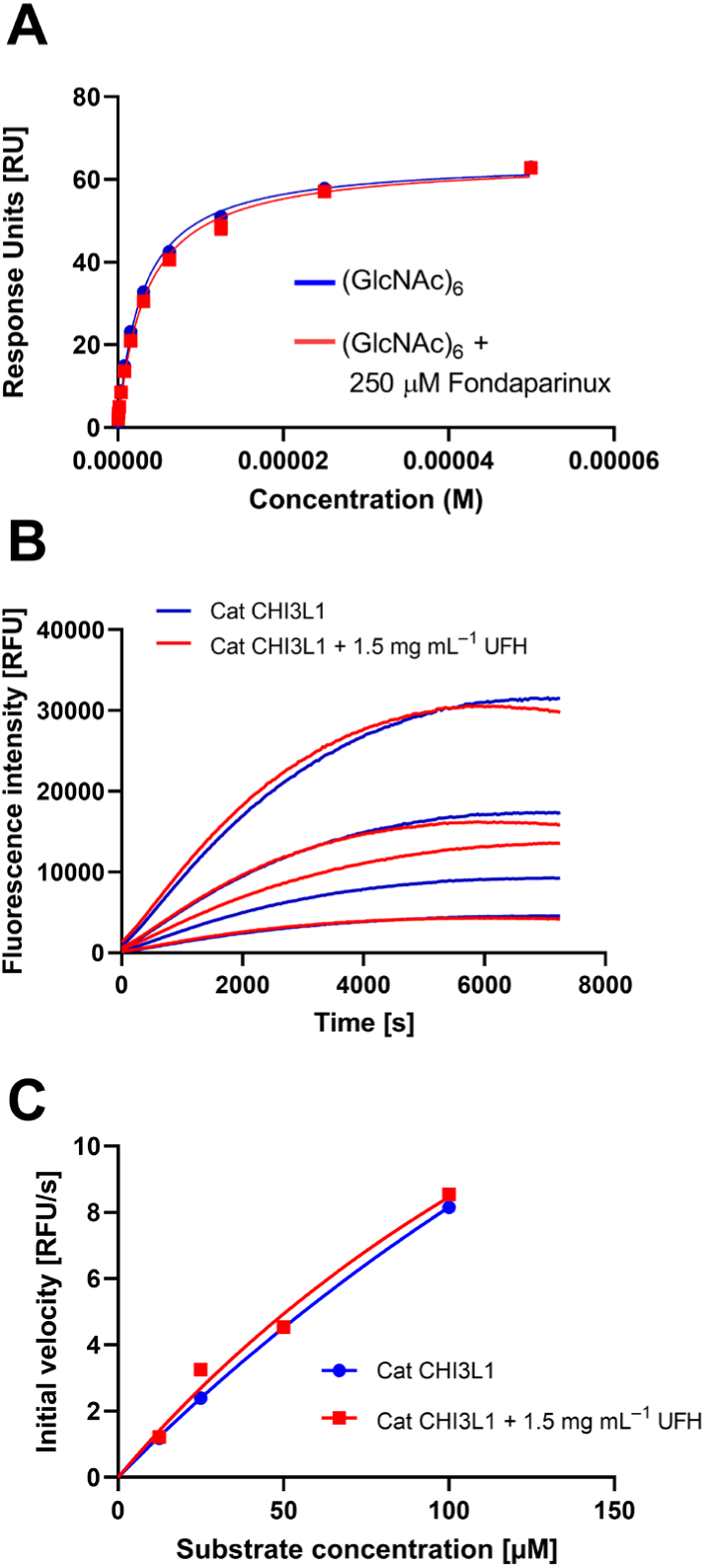
Experimental evidence for the lack of competition between COS and GAG ligands of CHI3L1. (A) SPR binding isotherm of (GlcNAc)_6_ binding to immobilized CHI3L1 in the absence or presence of 250 µM Fondaparinux. (B) Enzymatic activity of CHI3L1 in the absence or presence of UFH. CHI3L1 was incubated with a fluorogenic substrate and fluorescence intensity at 450 nm was recorded as a function of reaction time. The curves represent the mean of three independent experiments. (C) Global fitting of the initial velocities obtained in (B) to the Michaelis–Menten equation.

### COS and GAG ligands modulate physiologically relevant protein–protein interactions of CHI3L1 differently

Galectin-3 has been confirmed as a functionally relevant binding partner for CHI3L1. It has been hypothesized that the interaction between CHI3L1 and galectin-3 is mediated by the chitin-binding site. To shed light on this interaction and its interplay with canonical CHI3L1 ligands, we determined the binding affinity of the proteins in the presence and absence of carbohydrate ligands (Figure 10, Table 2). MST experiments revealed that CHI3L1 binds galectin-3 with a *K*_D_ value of 8.1 µM. With the COS-binding deficient mutant CBmut CHI3L1, binding affinity did not change significantly. This indicates that galectin-3 utilizes a fundamentally different binding mode compared with COS ligands, which renders it insensitive to the loss of Trp residues within the binding site. The addition of 500 µM (GlcNAc)6, however, fully abrogated the physiologically relevant protein-protein interaction (Figure S29). This observation is in line with previous hypotheses that galectin-3 engages CHI3L1 via its hydrophobic chitin-binding site.(80) Contrastingly, the presence of the GAG ligand Fondaparinux in a concentration of 500 µM led to a ca. 14-fold increase in the CHI3L1/galectin-3 binding affinity. This observation was unexpected, as previous experiments had not revealed any meaningful interaction between the COS and GAG binding sites. In contrast, when the protein ligand galectin-3 was bound to the chitin-binding site, marked cooperativity was observed. Although the basis for the differential recognition of COS and protein ligands at the CHI3L1 chitin-binding site remains unclear, these findings demonstrate that the physiologically relevant interaction between galectin-3 and CHI3L1 is differentially modulated by distinct classes of carbohydrate ligands.

**Figure 10.**
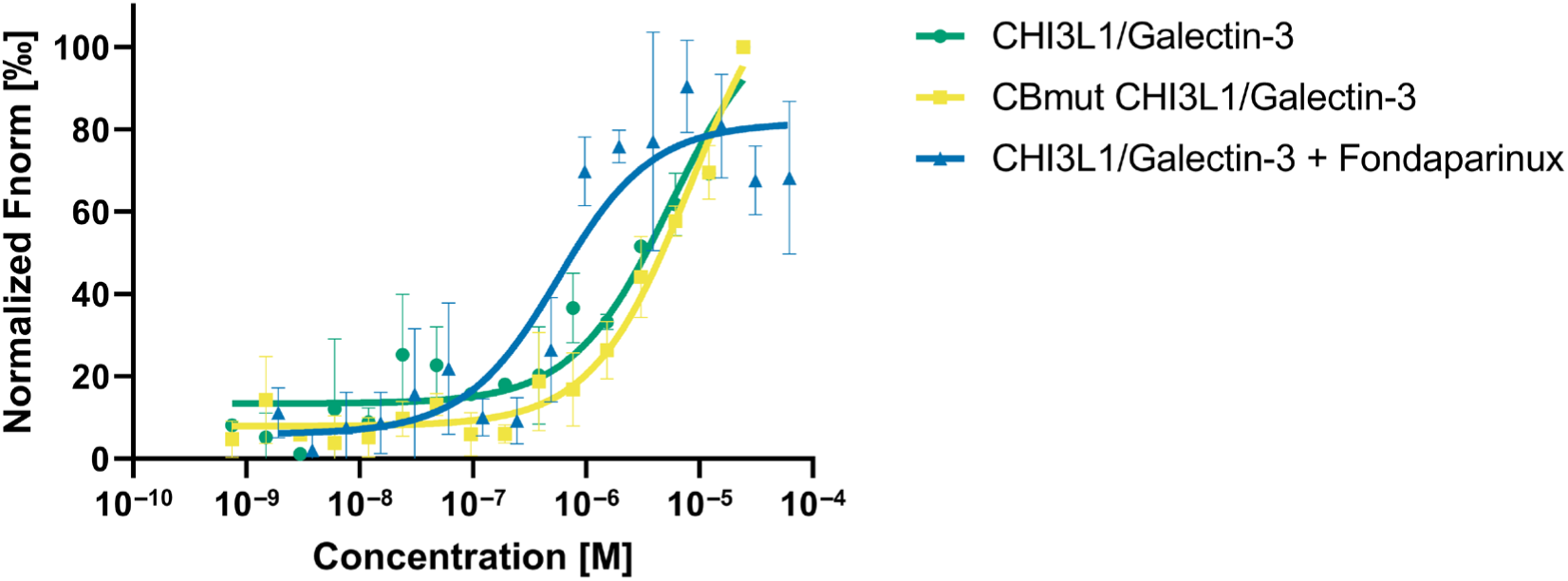
Interaction between CHI3L1 (green) or CBmut CHI3L1 (yellow) and galectin-3. The presence of 500 µM Fondaparinux increases the affinity between CHI3L1 and galectin-3 by ca. 14-fold (blue). Fnorm values were normalized to ease visual comparison between the datasets. Raw data is displayed in Figure S29.

**Table 2.** Binding affinity data for the interaction between CHI3L1 and galectin-3 determined by MST.

| Interaction | $K_D$ [μM] |
| --- | --- |
| CHI3L1 + galectin-3 | 8.1 (5.6-11.5) |
| CBmut CHI3L1 + galectin-3 | 16.3 (7.0-55.8) |
| CHI3L1 + (GlcNAc)<sub>6</sub> + galectin-3 | No Binding Detected |
| CHI3L1 + Fondaparinux + galectin-3 | 0.6 (0.4-0.7) |

Given that the CHI3L1/galectin-3 complex has been shown to regulate IL-13Rα2 signaling and to influence cell apoptosis and migration in a disease-relevant context, the results suggest that modulation of the respective binding affinities by cell-surface GAGs may contribute to the effector functions of the chitosome complex. Furthermore, this experiment provides direct evidence for a previously underappreciated immunomodulatory mechanism of COS, namely the inhibition of the CHI3L1/galectin-3 interaction.

## Discussion

CHI3L1 has emerged as an important biomarker and signaling molecule in inflammation, fibrosis, cancer, and neurodegenerative disease, yet the molecular basis of its interaction with GAGs remains incompletely understood. In the present study, we combined biophysical, structural, computational, and functional approaches to define the interaction of CHI3L1 with heparin/HS ligands and to compare this interaction with canonical COS recognition. Our data describe a previously suggested (18, 27, 28) but uncharacterized GAG-binding site centered on residues R144, R145, and K147 that is distinct from the well-established Trp-rich chitin-binding cleft. Binding of GAG ligands was found to depend strongly on the degree of polymerization, with longer heparin/HS chains engaging extended interaction networks beyond the primary binding site and binding affinity approaching a maximum at ca. DP10. Furthermore, GAG and COS ligands exhibited fundamentally different effects on CHI3L1 stability, fluorescence properties, and protein-protein interactions, supporting the existence of two functionally distinct and non-overlapping carbohydrate recognition interfaces. Finally, molecular simulations revealed that the identified R144/R145/K147 site represents the energetically preferred binding location for Fondaparinux and contributes to distinct unbinding pathways for longer HS oligomers.

A major strength of the present study is the convergence of several independent experimental and computational methods toward the same mechanistic model. Mutational analysis demonstrated that disruption of the R144/R145/K147 region abolished detectable GAG binding, whereas mutation of the canonical chitin-binding site preserved interaction with heparin ligands. Conversely, COS binding was eliminated by mutation of the Trp-rich chitin-binding cleft. STD-NMR epitope mapping further showed that Fondaparinux binds wild-type and chitin-binding-site mutants with highly similar ligand-contact patterns, indicating that the canonical chitin-binding site is not required for GAG recognition. These findings were independently supported by nanoDSF measurements, which revealed markedly different thermal and fluorescence responses upon GAG and COS binding, and by competition experiments, in which no inhibitory effect of heparin on the affinity of CHI3L1 or the enzymatic activity of the reactivated Cat CHI3L1 construct could be observed. These results demonstrate a lack of direct competition between the two ligand classes. In addition, molecular dynamics simulations, free-energy analyses, and ASMD unbinding calculations consistently identified the R144/R145/K147 region as the most favorable GAG-binding site and provided a structural rationale for the experimentally observed chain-length dependence.

While the evidence for distinct GAG and COS binding sites is compelling, some observations require further validation. The observation that Fondaparinux markedly enhances the affinity of the CHI3L1/galectin-3 interaction is intriguing but currently lacks a structural explanation. Although the effect was reproducibly observed by MST, the molecular basis of this apparent positive cooperativity remains unresolved. Likewise, the absolute free energies obtained from ASMD simulations should be interpreted cautiously because nonequilibrium pulling approaches are sensitive to sampling limitations and may overestimate dissociation energies. Nevertheless, the relative ranking of binding modes and the qualitative agreement with experimentally observed force profiles support the mechanistic conclusions drawn from the simulations.

The present findings both confirm and extend previous observations in the field. Earlier studies demonstrated that CHI3L1 binds chitin-derived oligosaccharides through a conserved aromatic groove formed by multiple tryptophan residues.(26, 27) More recently, Kognole and Payne provided the first evidence on the molecular interaction of heparin and related GAGs with CHI3L1, highlighting a broader carbohydrate-recognition repertoire beyond chitin-derived glycans.(28) The current work independently confirms GAG binding and further establishes that this interaction is mediated by a site distinct from the canonical chitin-binding cleft. In contrast to the KR-rich motif proposed previously as a functional heparin-binding domain,(29) neither molecular dynamics simulations, mutagenesis, nor ligand-binding experiments supported substantial binding at this location. Instead, our results identify the R144/R145/K147 region as the dominant GAG-recognition site. The observed dependence on GAG chain length additionally aligns with recent reports demonstrating interactions of CHI3L1 with extracellular matrix GAGs and suggesting multivalent binding mechanisms.(18) Beyond confirming GAG recognition, the present study provides several novel insights, including experimental evidence for non-overlapping GAG and COS binding sites, structural characterization of the GAG-binding epitope, quantification of ligand-dependent effects on protein stability, and the discovery that carbohydrate ligands differentially regulate the physiologically relevant interaction between CHI3L1 and galectin-3.

In summary, this study establishes CHI3L1 as a dual carbohydrate-binding protein that recognizes GAGs and chitin-derived oligosaccharides through distinct molecular interfaces. By integrating mutagenesis, biophysical characterization, NMR spectroscopy, molecular simulations, and functional interaction studies, we provide a coherent mechanistic framework for understanding how CHI3L1 engages structurally diverse glycans. These findings refine current models of CHI3L1 ligand recognition and suggest that differential engagement of the GAG-and chitin-binding sites may modulate downstream biological functions. Given the central role of CHI3L1 in inflammatory, fibrotic, and malignant diseases, the identification of a discrete GAG-binding interface may also facilitate the development of targeted molecular probes and therapeutic strategies aimed at selectively modulating CHI3L1 signaling pathways.

## Data availability

All files necessary to reproduce the simulations, including protein and ligand starting structures, AMBER topology and force-field parameter files, as well as initial configurations, restart files, and AMBER input files (PDB, PARM7, RST7, and text formats), are available through researchdata.hhu.de.

## Acknowledgements

The authors are grateful for financial support by the German Research Foundation (DFG SPP 2416 (GO 1367/7-1 (525826480), CR 811/3-1 (525826480), GO 2528/10-1 (525894428)): “CodeChi - Chitin, chitosan and chito-oligosaccharides and their interaction with proteins of the extracellular matrix and cellular signaling”). We thank Prof. Dr. Hakon Leffler and Prof. Dr. Ulf Nilsson from Lund University for providing recombinant galectin-3 samples for interaction studies. HG is grateful for the computational infrastructure and support provided by the “Zentrum für Informations-und Medientechnologie” (ZIM) at Heinrich Heine University Düsseldorf and the computing time provided by the John von Neumann Institute for Computing (NIC) on the supercomputers JUWELS and JUPITER at Jülich Supercomputing Centre (JSC) (user ID: chitin). We acknowledge access to the Jülich-Düsseldorf Biomolecular NMR Center.

## Author contributions

Ö.K. performed biophysical experiments and analyzed the resulting data, N.R. performed molecular dynamics simulations and analyzed the resulting data, M.G. conducted STD-NMR experiments, A.G. provided recombinant CHI3L1 samples for this study and conducted the glycan array experiment, C.G. supervised experiments and acquired funding, J.C. and H.G. conceptualized the study, supervised experiments, and acquired funding. All authors contributed to manuscript preparation and document revision.

## Conflicts of interest

There are no conflicts to declare.

## Notes

### Competing Interest Statement

The authors have declared no competing interest.

### Summary of Updates

The revised manuscript substantially extends the previous version by adding and integrating several new experimental and computational analyses: * A glycan-array analysis was added, providing independent evidence for a sharp increase in CHI3L1 binding for longer GAGs (around DP9-12) and strengthening the chain-length dependence. * The MD analysis was expanded to include protein RMSD/RMSF, ligand end-to-end distances, and time-resolved salt-bridge analyses, providing additional structural characterization of GAG binding. * The ASMD simulations were extended and analyzed in greater detail, including force-distance profiles and conformationally distinct unbinding pathways, with comparison to experimental force measurements. * Additional nanoDSF data with unfractionated heparin further support the distinct effects of GAGs versus COS on CHI3L1 stability and fluorescence. * The galectin-3 interaction experiments and their interpretation were expanded, including the reproducible ~14-fold enhancement by Fondaparinux and inhibition by COS. * The Discussion was substantially revised and broadened, placing the findings in the context of previous GAG-binding models, explicitly discussing limitations of the ASMD free energies and the unresolved mechanism of the Fondaparinux effect. Overall, the new version provides a more comprehensive, independently supported mechanistic model for distinct GAG and chitin-binding interfaces of CHI3L1.

## References

1. Johansen, J. S., Williamson, M. K., Rice, J. S., and Price, P. A. (1992) Identification of proteins secreted by human osteoblastic cells in culture. Journal of Bone and Mineral Research. 7, 501–512

2. Suzuki, K., Okawa, K., Ohkura, M., Kanaizumi, T., Kobayashi, T., Takahashi, K., et al. (2024) Evolutionary insights into sequence modifications governing chitin recognition and chitinase inactivity in YKL-40 (HC-gp39, CHI3L1). Journal of Biological Chemistry. 300, 107365

3. He, C. H., Lee, C. G., Dela Cruz, C. S., Lee, C.-M., Zhou, Y., Ahangari, F., et al (2013) Chitinase 3-like 1 Regulates Cellular and Tissue Responses via IL-13 Receptor α2. Cell Rep. 4, 830–841

4. Bakula, D., Vrkljan, N., Ratkajec, V., Glavcic, G., Miler, M., Pelajic, S., et al. (2024) YKL-40 as a biomarker in various inflammatory diseases: A review. Biochem. Med. (Zagreb). 34, 42–56

5. Pelkmans, W., Shekari, M., Brugulat-Serrat, A., Sánchez-Benavides, G., Minguillón, C., Fauria, K., et al. (2024) Astrocyte biomarkers GFAP and YKL-40 mediate early Alzheimer’s disease progression. Alzheimer’s & Dementia. 20, 483–493

6. Vázquez-Del Mercado, M., Pérez-Vázquez, F., Márquez-Aguirre, A. L., Martínez-García, E.-A., Chavarria-Avila, E., Ramos-Becerra, C. G., et al. (2023) YKL-40 serum levels are predicted by inflammatory state, age and diagnosis of idiopathic inflammatory myopathies. Sci. Rep. 13, 19172

7. Tizaoui, K., Yang, J. W., Lee, K. H., Kim, J. H., Kim, M., Yoon, S., et al. (2022) The role of YKL-40 in the pathogenesis of autoimmune diseases: a comprehensive review. Int. J. Biol. Sci. 18, 3731–3746

8. Connolly, K., Lehoux, M., O’Rourke, R., Assetta, B., Erdemir, G. A., Elias, J. A., et al. (2023) Potential role of chitinase-3-like protein 1 (CHI3L1/YKL-40) in neurodegeneration and Alzheimer’s disease. Alzheimer’s & Dementia. 19, 9–24

9. Hermansson, L., Yilmaz, A., Axelsson, M., Blennow, K., Fuchs, D., Hagberg, L., et al. (2019) Cerebrospinal fluid levels of glial marker YKL-40 strongly associated with axonal injury in HIV infection. J. Neuroinflammation. 16, 16

10. Outinen, T. K., Mantula, P., Jaatinen, P., Hämäläinen, M., Moilanen, E., Vaheri, A., et al. (2019) Glycoprotein YKL-40 Is Elevated and Predicts Disease Severity in Puumala Hantavirus Infection. Viruses. 10.3390/v11090767

11. Przekora, J., Synowiec, A., Kubiak, J. Z., Gościńska, A., Kalicki, B., and Jobs, K. (2024) Assessment of urine calprotectin and YKL-40 levels in urinary tract infection diagnosis in children under 2 years of age. Sci. Rep. 14, 28695

12. Hedetoft, M., Hansen, M. B., Madsen, M. B., Johansen, J. S., and Hyldegaard, O. (2021) Associations between YKL-40 and markers of disease severity and death in patients with necrotizing soft-tissue infection. BMC Infect. Dis. 21, 1046

13. Kjaergaard, A. D., Vaag, A. A., Jensen, V. H., Olsen, M. H., Højlund, K., Vestergaard, P., et al. (2025) YKL-40 and risk of incident cancer in early type 2 diabetes: a Danish cohort study. Br. J. Cancer. 132, 1019–1026

14. Böckelmann, L. C., Felix, T., Calabrò, S., and Schumacher, U. (2021) YKL-40 protein expression in human tumor samples and human tumor cell line xenografts: implications for its use in tumor models. Cellular Oncology. 44, 1183–1195

15. Junker, N., Johansen, J. S., Hansen, L. T., Lund, E. L., and Kristjansen, P. E. G. (2005) Regulation of YKL-40 expression during genotoxic or microenvironmental stress in human glioblastoma cells. Cancer Sci. 96, 183–90

16. Oh, I. H., Pyo, J.-S., and Son, B. K. (2021) Prognostic Impact of YKL-40 Immunohistochemical Expression in Patients with Colorectal Cancer. Curr. Oncol. 28, 3139–3149

17. Wang, Z., Wang, S., Jia, Z., Hu, Y., Cao, D., Yang, M., et al. (2023) YKL-40 derived from infiltrating macrophages cooperates with GDF15 to establish an immune suppressive microenvironment in gallbladder cancer. Cancer Lett. 563, 216184

18. Magnusdottir, U., Yang Jonatansdottir, Y., Oskarsson, K. R., Hjorleifsson, J. G., Einarsson, J. M., and Thormodsson, F. R. (2025) Characterization of YKL-40 Binding to Extracellular Matrix Glycosaminoglycans. Mar. Drugs. 23, 379

19. Bigg, H. F., Wait, R., Rowan, A. D., and Cawston, T. E. (2006) The Mammalian Chitinase-like Lectin, YKL-40, Binds Specifically to Type I Collagen and Modulates the Rate of Type I Collagen Fibril Formation. Journal of Biological Chemistry. 281, 21082–21095

20. Zhou, Y., He, C. H., Yang, D. S., Nguyen, T., Cao, Y., Kamle, S., et al. (2018) Galectin-3 Interacts with the CHI3L1 Axis and Contributes to Hermansky–Pudlak Syndrome Lung Disease. The Journal of Immunology. 200, 2140–2153

21. Areshkov, P. O., Avdieiev, S. S., Balynska, O. V., LeRoith, D., and Kavsan, V. M. (2012) Two Closely Related Human Members of Chitinase-like Family, CHI3L1 and CHI3L2, Activate ERK1/2 in 293 and U373 Cells but Have the Different Influence on Cell Proliferation. Int. J. Biol. Sci. 8, 39–48

22. Yu, J. E., Yeo, I. J., Han, S.-B., Yun, J., Kim, B., Yong, Y. J., et al. (2024) Significance of chitinase-3-like protein 1 in the pathogenesis of inflammatory diseases and cancer. Exp. Mol. Med. 56, 1–18

23. Lee, Y. S., Yu, J. E., Kim, K. C., Lee, D. H., Son, D. J., Lee, H. P., et al. (2022) A small molecule targeting CHI3L1 inhibits lung metastasis by blocking IL-13Rα2-mediated JNK-AP-1 signals. Mol. Oncol. 16, 508–526

24. Kaur, B., Denzinger, K., Zhang, L., García-Vázquez, N., Wolber, G., and Gabr, M. (2025) Virtual Screening-Guided Discovery of Small-Molecule CHI3L1 Inhibitors with Functional Activity in Glioblastoma Spheroids. ACS Med. Chem. Lett. 16, 2346–2353

25. Czestkowski, W., Krzemiński, Ł., Piotrowicz, M. C., Mazur, M., Pluta, E., Andryianau, G., et al. (2024) Structure-Based Discovery of High-Affinity Small Molecule Ligands and Development of Tool Probes to Study the Role of Chitinase-3-Like Protein 1. J. Med. Chem. 67, 3959–3985

26. Houston, D. R., Recklies, A. D., Krupa, J. C., and van Aalten, D. M. F. (2003) Structure and Ligand-induced Conformational Change of the 39-kDa Glycoprotein from Human Articular Chondrocytes. Journal of Biological Chemistry. 278, 30206–30212

27. Fusetti, F., Pijning, T., Kalk, K. H., Bos, E., and Dijkstra, B. W. (2003) Crystal Structure and Carbohydrate-binding Properties of the Human Cartilage Glycoprotein-39. Journal of Biological Chemistry. 278, 37753–37760

28. Kognole, A. A., and Payne, C. M. (2017) Inhibition of Mammalian Glycoprotein YKL-40. Journal of Biological Chemistry. 292, 2624–2636

29. Ngernyuang, N., Yan, W., Schwartz, L. M., Oh, D., Liu, Y., Chen, H., et al. (2018) A Heparin Binding Motif Rich in Arginine and Lysine is the Functional Domain of YKL-40. Neoplasia. 20, 182–192

30. Magnusdottir, U., Thormodsson, F. R., Kjalarsdottir, L., Filippusson, H., Gislason, J., Oskarsson, K. R., et al. (2025) Heparin-binding of the human chitinase-like protein YKL-40 is allosterically modified by chitin oligosaccharides. Biochem. Biophys. Rep. 41, 101908

31. [Preprint] Großdorf, A., Nabil, S., Imelmann, C., Hellmann, M. J., Froese, J., El-Awaad, E., et al. (2026) Chitinase-3-like protein 1 decodes chitosan acetylation patterns into toll-like receptor 2 signaling through heparan sulfate. 10.64898/2026.06.15.732264

32. Dai, X., Liu, W., Zhou, Q., Cheng, C., Yang, C., Wang, S., et al. (2016) Formal Synthesis of Anticoagulant Drug Fondaparinux Sodium. J. Org. Chem. 81, 162–184

33. Feng, K., Wang, K., Zhou, Y., Xue, H., Wang, F., Jin, H., et al. (2023) Non-Anticoagulant Activities of Low Molecular Weight Heparins—A Review. Pharmaceuticals. 16, 1254

34. Hricovíni, M., and Torri, G. (1995) Dynamics in aqueous solutions of the pentasaccharide corresponding to the binding site of heparin for antithrombin III studied by NMR relaxation measurements. Carbohydr. Res. 268, 159–175

35. Gopalswamy, M., Kroeger, T., Bickel, D., Frieg, B., Akter, S., Schott-Verdugo, S., et al. (2022) Biophysical and pharmacokinetic characterization of a small-molecule inhibitor of RUNX1/ETO tetramerization with anti-leukemic effects. Sci. Rep. 12, 14158

36. Mayer, M., and Meyer, B. (1999) Characterization of Ligand Binding by Saturation Transfer Difference NMR Spectroscopy. Angew. Chem. Int. Ed Engl. 38, 1784–1788

37. Madhavi S., G., Adzhigirey, M., Day, T., Annabhimoju, R., and Sherman, W. (2013) Protein and ligand preparation: parameters, protocols, and influence on virtual screening enrichments. J. Comput. Aided. Mol. Des. 27, 221–234

38. Singh, A., Montgomery, D., Xue, X., Foley, B. L., and Woods, R. J. (2019) GAG Builder: a web-tool for modeling 3D structures of glycosaminoglycans. Glycobiology. 29, 515–518

39. Martínez, L., Andrade, R., Birgin, E. G., and Martínez, J. M. (2009) A package for building initial configurations for molecular dynamics simulations. J. Comput. Chem. 30, 2157–2164

40. Izadi, S., Anandakrishnan, R., and Onufriev, A. V. (2014) Building Water Models: A Different Approach. J. Phys. Chem. Lett. 5, 3863–3871

41. Machado, M. R., and Pantano, S. (2020) Split the Charge Difference in Two! A Rule of Thumb for Adding Proper Amounts of Ions in MD Simulations. J. Chem. Theory Comput. 16, 1367– 1372

42. Case, D. A., et al. (2023) AmberTools. J. Chem. Inf. Model. 63, 6183–6191.

43. Tian, C., Kasavajhala, K., Belfon, K. A. A., Raguette, L., Huang, H., Migues, A. N., et al. (2020) ff19SB: Amino-Acid-Specific Protein Backbone Parameters Trained against Quantum Mechanics Energy Surfaces in Solution. J. Chem. Theory Comput. 16, 528–552

44. Kirschner, K. N., Yongye, A. B., Tschampel, S. M., González-Outeiriño, J., Daniels, C. R., Foley, B. L., and Woods, R. J. (2008) GLYCAM06: A generalizable biomolecular force field. Carbohydrates. J. Comput. Chem. 29, 622–655

45. Sengupta, A., Li, Z., Song, L. F., Li, P., and Merz, K. M. (2021) Parameterization of Monovalent Ions for the OPC3, OPC, TIP3P-FB, and TIP4P-FB Water Models. J. Chem. Inf. Model. 61, 869–880

46. Li, P., Song, L. F., and Merz, K. M. (2015) Systematic Parameterization of Monovalent Ions Employing the Nonbonded Model. J. Chem. Theory Comput. 11, 1645–1657

47. Le Grand, S., Götz, A. W., and Walker, R. C. (2013) SPFP: Speed without compromise—A mixed precision model for GPU accelerated molecular dynamics simulations. Comput. Phys. Commun. 184, 374–380

48. Salomon-Ferrer, R., Götz, A. W., Poole, D., Le Grand, S., and Walker, R. C. (2013) Routine Microsecond Molecular Dynamics Simulations with AMBER on GPUs. 2. Explicit Solvent Particle Mesh Ewald. J. Chem. Theory Comput. 9, 3878–3888

49. Ryckaert, J.-P., Ciccotti, G., and Berendsen, H. J. C. (1977) Numerical integration of the cartesian equations of motion of a system with constraints: molecular dynamics of n-alkanes. J. Comput. Phys. 23, 327–341

50. Hopkins, C. W., Le Grand, S., Walker, R. C., and Roitberg, A. E. (2015) Long-Time-Step Molecular Dynamics through Hydrogen Mass Repartitioning. J. Chem. Theory Comput. 11, 1864–1874

51. Darden, T., York, D., and Pedersen, L. (1993) Particle mesh Ewald: An *N* ⍰log( *N* ) method for Ewald sums in large systems. J. Chem. Phys. 98, 10089–10092

52. Yüksel, S., Bonus, M., Schwabe, T., Pfleger, C., Zimmer, T., Enke, U., et al. (2022) Uncoupling of Voltage-and Ligand-Induced Activation in HCN2 Channels by Glycine Inserts. Front. Physiol. 10.3389/fphys.2022.895324

53. Berendsen, H. J. C., Postma, J. P. M., van Gunsteren, W. F., DiNola, A., and Haak, J. R. (1984) Molecular dynamics with coupling to an external bath. J. Chem. Phys. 81, 3684–3690

54. Loncharich, R. J., Brooks, B. R., and Pastor, R. W. (1992) Langevin dynamics of peptides: The frictional dependence of isomerization rates of *N* -acetylalanyl-*N* ʹ-methylamide. Biopolymers. 32, 523–535

55. Roe, D. R., and Cheatham, T. E. (2013) PTRAJ and CPPTRAJ: Software for Processing and Analysis of Molecular Dynamics Trajectory Data. J. Chem. Theory Comput. 9, 3084–3095

56. Michaud-Agrawal, N., Denning, E. J., Woolf, T. B., and Beckstein, O. (2011) MDAnalysis: A toolkit for the analysis of molecular dynamics simulations. J. Comput. Chem. 32, 2319–2327

57. Humphrey, W., Dalke, A., and Schulten, K. (1996) VMD: Visual molecular dynamics. J. Mol. Graph. 14, 33–38

58. Schrödinger, LLC. (2015) The PyMOL Molecular Graphics System (Version 1.8).

59. Gopalswamy, M., Bickel, D., Dienstbier, N., Tu, J.-W., Vogt, M., Schott-Verdugo, S., et al. (2025) Identification of non-charged 7.44 analogs interacting with the NHR2 domain of RUNX1-ETO with improved antiproliferative effect in RUNX-ETO positive cells. Sci. Rep. 15, 17720

60. Virtanen, P., Gommers, R., Oliphant, T. E., Haberland, M., Reddy, T., Cournapeau, D., et al. (2020) SciPy 1.0: fundamental algorithms for scientific computing in Python. Nat. Methods. 17, 261–272

61. Ozer, G., Quirk, S., and Hernandez, R. (2012) Thermodynamics of Decaalanine Stretching in Water Obtained by Adaptive Steered Molecular Dynamics Simulations. J. Chem. Theory Comput. 8, 4837–4844

62. Ozer, G., Valeev, E. F., Quirk, S., and Hernandez, R. (2010) Adaptive Steered Molecular Dynamics of the Long-Distance Unfolding of Neuropeptide Y. J. Chem. Theory Comput. 6, 3026–3038

63. Park, S., and Schulten, K. (2004) Calculating potentials of mean force from steered molecular dynamics simulations. J. Chem. Phys. 120, 5946–5961

64. Jarzynski, C. (1997) Nonequilibrium Equality for Free Energy Differences. Phys. Rev. Lett. 78, 2690–2693

65. Dittrich, J., Brethauer, C., Goncharenko, L., Bührmann, J., Zeisler-Diehl, V., Pariyar, S., et al. (2022) Rational Design Yields Molecular Insights on Leaf-Binding of Anchor Peptides. ACS Appl. Mater. Interfaces. 14, 28412–28426

66. Govind Kumar, V., Polasa, A., Agrawal, S., Kumar, T. K. S., and Moradi, M. (2022) Binding affinity estimation from restrained umbrella sampling simulations. Nat. Comput. Sci. 3, 59–70

67. Zhu, J., Li, Y., Wang, J., Yu, Z., Liu, Y., Tong, Y., and Han, W. (2018) Adaptive Steered Molecular Dynamics Combined With Protein Structure Networks Revealing the Mechanism of Y68I/G109P Mutations That Enhance the Catalytic Activity of D-psicose 3-Epimerase From Clostridium Bolteae. Front. Chem. 10.3389/fchem.2018.00437

68. Hantz, E. R., and Lindert, S. (2021) Adaptative Steered Molecular Dynamics Study of Mutagenesis Effects on Calcium Affinity in the Regulatory Domain of Cardiac Troponin C. J. Chem. Inf. Model. 61, 3052–3057

69. Allen, C., Bureau, H. R., McGee, T. D., Quirk, S., and Hernandez, R. (2022) Benchmarking Adaptive Steered Molecular Dynamics (ASMD) on CHARMM Force Fields. ChemPhysChem. 10.1002/cphc.202200175

70. Gohlke, H., Hergert, U., Meyer, T., Mulnaes, D., Grieshaber, M. K., Smits, S. H. J., et al. (2013) Binding Region of Alanopine Dehydrogenase Predicted by Unbiased Molecular Dynamics Simulations of Ligand Diffusion. J. Chem. Inf. Model. 53, 2493–2498

71. Kaiser, J., Gertzen, C. G. W., Mann, D., Sachse, C., and Gohlke, H. (2026) Evidence for Epibatidine Binding to the Desensitization Gate in α7 nAChR from Molecular Dynamics Simulations and Cryo-EM. J. Chem. Inf. Model. 66, 1337–1341

72. Janke, J. J., Yu, Y., Pomin, V. H., Zhao, J., Wang, C., Linhardt, R. J., and García, A. E. (2022) Characterization of Heparin’s Conformational Ensemble by Molecular Dynamics Simulations and Nuclear Magnetic Resonance Spectroscopy. J. Chem. Theory Comput. 18, 1894–1904

73. Marcisz, M., and Samsonov, S. A. (2023) Solvent Model Benchmark for Molecular Dynamics of Glycosaminoglycans. J. Chem. Inf. Model. 63, 2147–2157

74. Frieg, B., Gremer, L., Heise, H., Willbold, D., and Gohlke, H. (2020) Binding modes of thioflavin T and Congo red to the fibril structure of amyloid-β(1–42). Chemical Communications. 56, 7589–7592

75. Cardin, A. D., and Weintraub, H. J. (1989) Molecular modeling of protein-glycosaminoglycan interactions. *Arteriosclerosis: An Official Journal of the American Heart Association*, Inc. 9, 21–32

76. Xu, X., Angioletti-Uberti, S., Lu, Y., Dzubiella, J., and Ballauff, M. (2019) Interaction of Proteins with Polyelectrolytes: Comparison of Theory to Experiment. Langmuir. 35, 5373–5391

77. Knight, B. M., Gallagher, C. M. B., Schulz, M. D., Edgar, K. J., McNaul, C. D., McCutchin, C. A., and Dove, P. M. (2025) Thermodynamics of calcium binding to heparin: Implications of solvation and water structuring for polysaccharide biofunctions. Proceedings of the National Academy of Sciences. 10.1073/pnas.2504348122

78. Gentile, R., Modric, M., Thiele, B., Jaeger, K.-E., Kovacic, F., Schott-Verdugo, S., and Gohlke, H. (2024) Molecular Mechanisms Underlying Medium-Chain Free Fatty Acid-Regulated Activity of the Phospholipase PlaF from *Pseudomonas aeruginosa*. JACS Au. 4, 958–973

79. Ho, B. K., and Agard, D. A. (2010) An Improved Strategy for Generating Forces in Steered Molecular Dynamics: The Mechanical Unfolding of Titin, e2lip3 and Ubiquitin. PLoS One. 5, e13068

80. Chen, A., Jiang, Y., Li, Z., Wu, L., Santiago, U., Zou, H., et al. (2021) Chitinase-3-like 1 protein complexes modulate macrophage-mediated immune suppression in glioblastoma. Journal of Clinical Investigation. 10.1172/JCI147552

