## Supporting Information for "Dual Carbohydrate Recognition by the Chitinase-like Protein CHI3L1 Through Distinct Glycosaminoglycan and Chitin-Binding Interfaces"

1. Supporting Tables

**Table S1:** Intial heavy-atom distance between the ligand and CHI3L1 for the starting configurations of the different systems.

**Table S2:** Simulation protocol. Characteristics of the minimization, thermalization, density adaptation, and production phases for the CHI3L1-COS systems.

**Table S3:** Binding events of the different systems.

2. Supporting Figures

**Figure S1:** SPR binding isotherms of the interaction between CHI3L1 with COS.

**Figure S2:** Binding isotherms of the interactions between CHI3L1 and GAGs.

**Figure S3:** Structural variability of CHI3L1 during the fldMD simulations with Fondaparinux

**Figure S4:** Structural variability of CHI3L1 during the fldMD simulations with HS9.

**Figure S5:** Structural variability of CHI3L1 during the fldMD simulations with $HS9_{\Delta 2S/6S}$.

**Figure S6:** Structural variability of HepBmut (R144A, R145A, and K147A) CHI3L1 during the fldMD simulations with HS9.

**Figure S7:** Electrostatic surface potential of CHI3L1.

**Figure S8:** Unbiased MD simulations of Fondaparinux diffusion with CHI3L1.

**Figure S9:** Unbiased MD simulations of HS9 diffusion with CHI3L1.

**Figure S10:** Unbiased MD simulations of $HS9_{\Delta 2S/6S}$ diffusion with CHI3L1.

**Figure S11:** Unbiased MD simulations of $HS9_{\Delta 2S/6S}$ diffusion with HepBmut (R144A, R145A, K147A) CHI3L1

**Figure S12:** Conformations of Fondaparinux during the fldMD simulations with CHI3L1.

**Figure S13:** Conformations of HS9 during the fldMD simulations with CHI3L1

**Figure S14:** Conformations of $HS9_{\Delta 2S/6S}$ during the fldMD simulations with CHI3L1.

**Figure S15:** Conformations of HS9 during the fldMD simulations with HepBmut (R144A, R145A, and K147A) CHI3L1.

**Figure S16:** Clustering analysis of Fondaparinux poses based on interaction fingerprints with CHI3L1.

**Figure S17:** Clustering analysis of HS9 poses based on interaction fingerprints with CHI3L1.

**Figure S18:** Clustering analysis of $HS9_{\Delta 2S/6S}$ poses based on interaction fingerprints with CHI3L1.

**Figure S19:** Clustering analysis of HS9 poses based on interaction fingerprints with HepBmut (R144A, R145A, and K147A) CHI3L1.

**Figure S20:** Probability density of $HS9_{\Delta 2S/6S}$ around CHI3L1.

**Figure S21:** Probability density of HS9 around HepBmut (R144A, R145A, and K147A) CHI3L1.

**Figure S22:** Contacts between CHI3L1 and $HS9_{\Delta 2S/6S}$.

**Figure S23:** Contacts between HepBmut (R144A, R145A, and K147A) CHI3L1 and HS9.

**Figure S24:** Atomic fluctuations calculated from MD simulations.

**Figure S25:** Numbering of the anionic groups of Fondaparinux and HS9 as reference for Figures S26-29.

**Figure S26:** Time-series analysis of Fondaparinux interactions with CHI3L1.

**Figure S27:** Time-series analysis of HS9 interactions with CHI3L1.

**Figure S28:** Time-series analysis of $HS9_{\Delta 2S/6S}$ interactions with CHI3L1.

**Figure S29:** Time-series analysis of HS9 interactions with HepBmut (R144A, R145A, and K147A) CHI3L1 and HS9.

**Figure S30:** MST binding isotherms of the interaction between CHI3L1 and galectin-3.

### Supporting Tables

**Table S1. Initial heavy-atom distance between the ligand and CHI3L1 for the starting configurations of the different systems.**

| Simulation | Distance[a] | | | |
|---|---|---|---|---|
| | CHI3L1/Fondaparinux | CHI3L1/HS9 | CHI3L1/HS9$_{\Delta 2S/6S}$ | HepBmut/HS9 |
| 1 | 23.1 | 16.9 | 20.5 | 20.2 |
| 2 | 20.6 | 21.4 | 18.8 | 21.4 |
| 3 | 27.2 | 17.8 | 28.0 | 21.2 |
| 4 | 25.6 | 15.2 | 19.1 | 19.9 |
| 5 | 19.7 | 23.6 | 25.2 | 19.2 |
| 6 | 16.5 | 17.4 | 25.1 | 18.3 |
| 7 | 23.0 | 20.0 | 17.8 | 19.0 |
| 8 | 18.6 | 19.0 | 20.8 | 22.7 |
| 9 | 16.0 | 18.9 | 23.1 | 20.1 |
| 10 | 15.2 | 17.0 | 17.8 | 20.0 |
| 11 | 19.2 | 16.3 | 19.2 | 20.0 |
| 12 | 21.5 | 14.7 | 19.1 | 18.3 |
| 13 | 12.7 | 15.9 | 16.5 | 22.5 |
| 14 | 22.1 | 19.8 | 20.2 | 22.1 |
| 15 | 25.0 | 18.7 | 22.2 | 18.4 |
| 16 | 19.1 | 21.1 | 19.9 | 22.1 |
| 17 | 16.2 | 18.1 | 20.5 | 19.1 |
| 18 | 18.4 | 21.1 | 22.6 | 20.0 |
| 19 | 14.3 | 18.0 | 17.2 | 18.7 |
| 20 | 17.4 | 27.0 | 21.4 | 17.7 |

[a]In Å; calculated using the “nativecontacts mindist” command in CPPTRAJ.

**Table S2. Simulation protocol. Characteristics of the minimization, thermalization, density adaptation, and production phases for the CHI3L1-COS systems.**

| Process | No. of Steps | Algorithm | Restrained Selection and Force Constant [kcal mol$^{-1}$ Å$^{-2}$] | | |
|---|---|---|---|---|---|
| | | | PBB[a] | PSC[b] | LIG[c] |
| Minimization | 2,500/2,500 | Steepest descent/Conjugate gradient | 10.0 | 5.0 | 2.5 |
| **Process** | **Time [ps]** | **Ensemble/Time Step [fs]** | **Restrained Selection and Force Constant [kcal mol$^{-1}$ Å$^{-2}$]** | | |
| | | | PBB[a] | PSC[b] | LIG[c] |
| Thermalization | 50.0 | NVT/1.0 | 10.0 | 5.0 | 2.5 |
| Density adaptation 1 | 100.0 | NPT/2.0 | 5.0 | 2.5 | 2.5 |
| Density adaptation 2 | 100.0 | NPT/2.0 | 2.5 | 1.0 | 1.0 |
| Density adaptation 3 | 100.0 | NPT/4.0 | 1.0 | 0.5 | 0.5 |
| Density adaptation 4 | 100.0 | NPT/4.0 | 0.5 | 0.1 | 0.1 |
| Density adaptation 5 | 100.0 | NPT/4.0 | 0.1 | – | – |
| Density adaptation 6 | 150.0 | NPT/4.0 | 0.05 | – | – |
| Density adaptation 7 | 300.0 | NPT/4.0 | – | – | – |
| Production | $1.0 \times 10^6$ | NPT/4.0 | – | – | – |

[a]Protein backbone atoms. [b]Protein side-chain heavy atoms. [c]Ligand atoms.

**Table S3. Binding events of the different systems.**

| Simulation | Number of Bound Frames/Binding Events[a] | | | |
|---|---|---|---|---|
| | CHI3L1/Fondaparinux | CHI3L1/HS9 | CHI3L1/HS9$_{\Delta 2S/6S}$ | HepBmut/HS9 |
| 1 | 428 | 746 | 896 | 555 |
| 2 | 364 | 781 | 493 | 73 |
| 3 | 482 | 578 | 471 | 555 |
| 4 | 425 | 646 | 685 | 557 |
| 5 | 664 | 737 | 567 | 311 |
| 6 | 402 | 285 | 618 | 454 |
| 7 | 416 | 713 | 729 | 240 |
| 8 | 306 | 697 | 872 | 238 |
| 9 | 311 | 236 | 164 | 244 |
| 10 | 484 | 907 | 239 | 462 |
| 11 | 660 | 725 | 248 | 377 |
| 12 | 210 | 166 | 520 | 541 |
| 13 | 186 | 317 | 731 | 201 |
| 14 | 297 | 770 | 847 | 231 |
| 15 | 215 | 704 | 460 | 297 |
| 16 | 432 | 787 | 495 | 161 |
| 17 | 646 | 966 | 475 | 227 |
| 18 | 131 | 532 | 262 | 385 |
| 19 | 592 | 469 | 10 | 740 |
| 20 | 270 | 89 | 596 | 66 |
| Total | 7,921 | 11,851 | 10,378 | 6,915 |
| Median | 409 (290-482)[b] | 700 (431-752) | 508 (410-696) | 304 (230-482)** |

[a]One frame corresponds to 1 ns of simulation time. [b]Interquartile range (25$^{th}$-75$^{th}$ percentiles). Statistical significance between the WT and mutant system for HS9 binding was evaluated using a two-sided Mann-Whitney U test (**$p \leq 0.01$).

### Supporting Figures

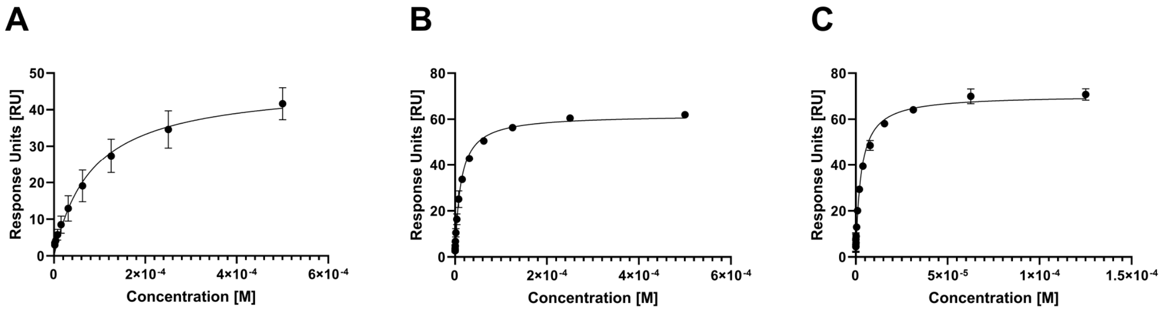


**Figure S1. SPR binding isotherms of the interaction between CHI3L1 and COS**. (A) CHI3L1 wild type (WT) and $(GlcNAc)_4$. (B) CHI3L1 WT and $(GlcNAc)_5$. (C) CHI3L1 WT and $(GlcNAc)_6$.

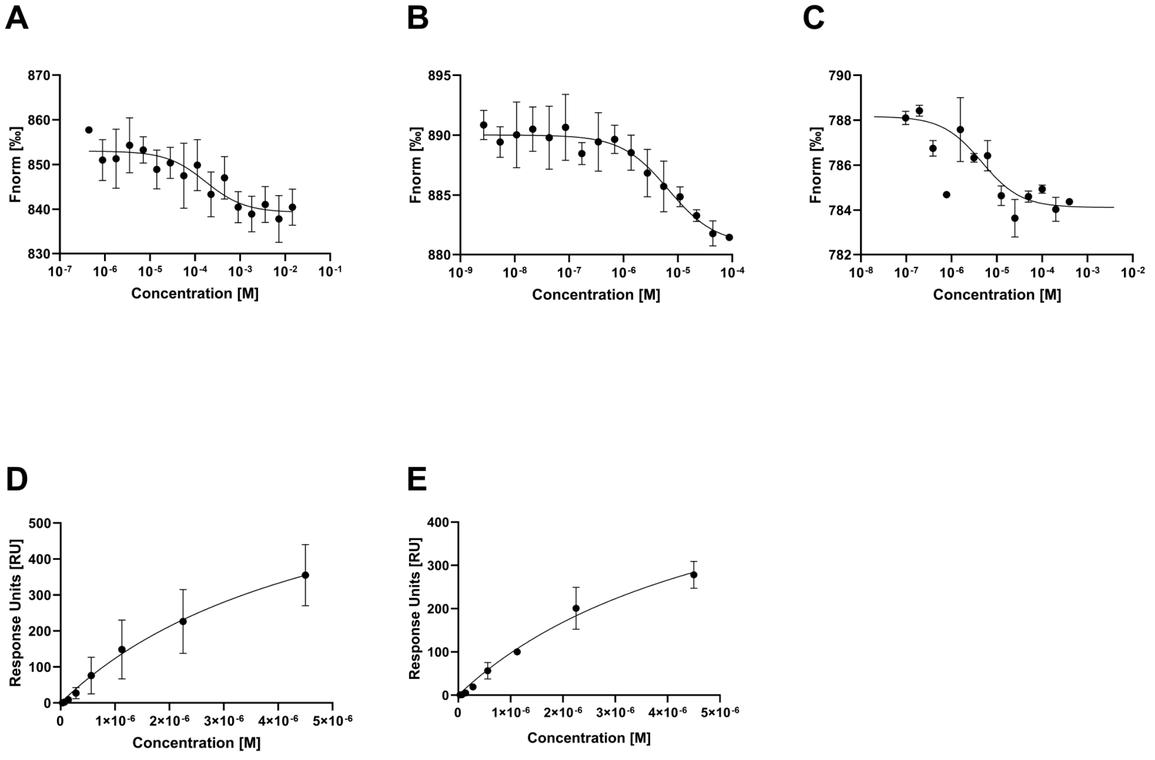


**Figure S2. Binding isotherms of the interactions between CHI3L1 and GAGs**. (A) Fondaparinux binding to CHI3L1 WT (MST). (B) HS9 binding to CHI3L1 WT (MST). (C) Enoxaparin binding to CHI3L1 WT (MST). (D) CHI3L1 WT binding to UFH coated surface (SPR). (E) CBmut CHI3L1 binding to UFH coated surface (SPR).

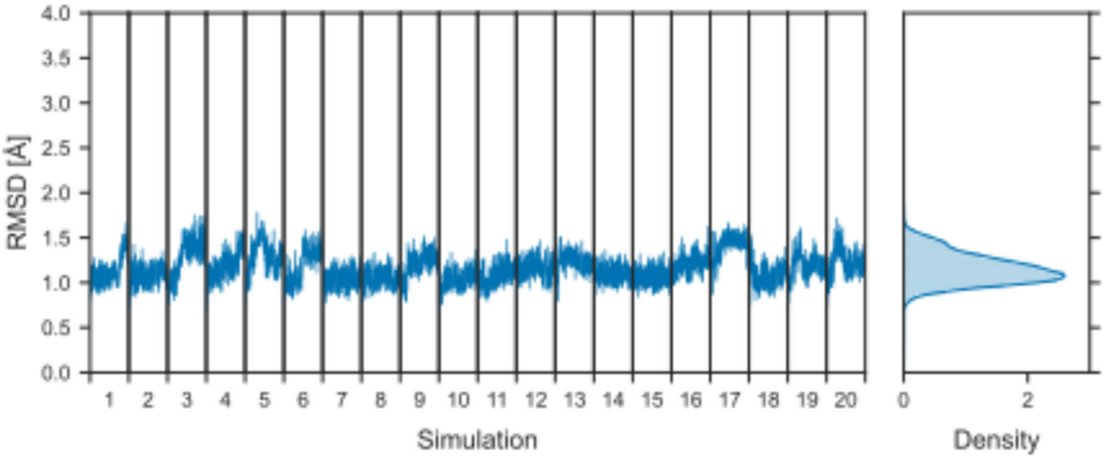


**Figure S3. Structural variability of CHI3L1 during the fldMD simulations with Fondaparinux.** Time courses (left) and probability distributions (right) of the RMSD of CHI3L1 ($C_\alpha$ atoms) shown for all of the 20 simulations (1 µs length). The smoothed distributions (thick, opaque lines) were calculated using a Gaussian kernel density estimator. The underlying data is displayed as a histogram (transparent bars). RMSD values were calculated relative to the crystal structure (PDB ID: 1HJX) after least-squares fitting of the $C_\alpha$ atoms.

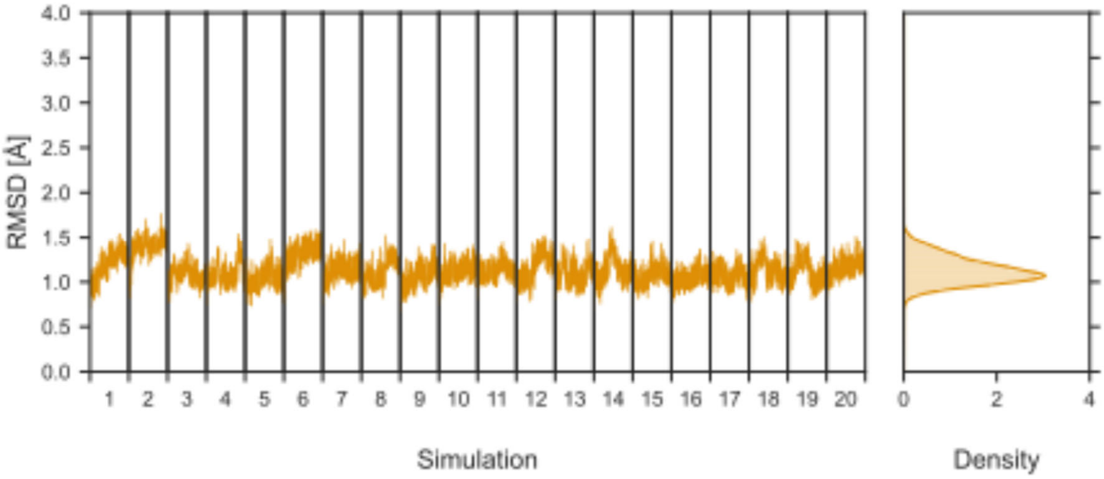


**Figure S4. Structural variability of CHI3L1 during the fldMD simulations with HS9.** Time courses (left) and probability distributions (right) of the RMSD of CHI3L1 ($C_\alpha$ atoms) shown for all of the 20 simulations (1 µs length). The smoothed distributions (thick, opaque lines) were calculated using a Gaussian kernel density estimator. The underlying data is displayed as a histogram (transparent bars). RMSD values were calculated relative to the crystal structure (PDB ID: 1HJX) after least-squares fitting of the $C_\alpha$ atoms.

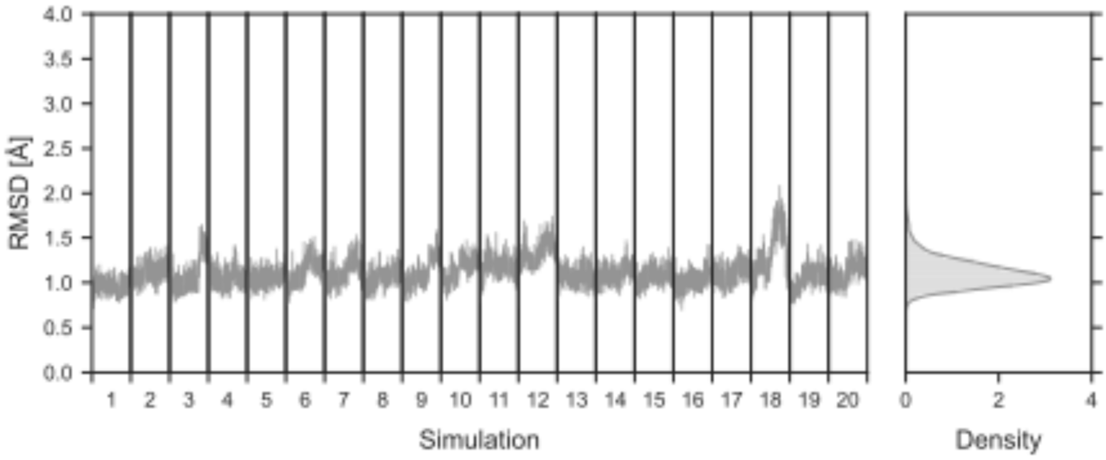


**Figure S5. Structural variability of CHI3L1 during the fldMD simulations with $HS9_{\Delta 2S/6S}$.** Time courses (left) and probability distributions (right) of the RMSD of CHI3L1 ($C_\alpha$ atoms) shown for all of the 20 simulations (1 µs length). The smoothed distributions (thick, opaque lines) were calculated using a Gaussian kernel density estimator. The underlying data is displayed as a histogram (transparent bars). RMSD values were calculated relative to the crystal structure (PDB ID: 1HJX) after least-squares fitting of the $C_\alpha$ atoms.

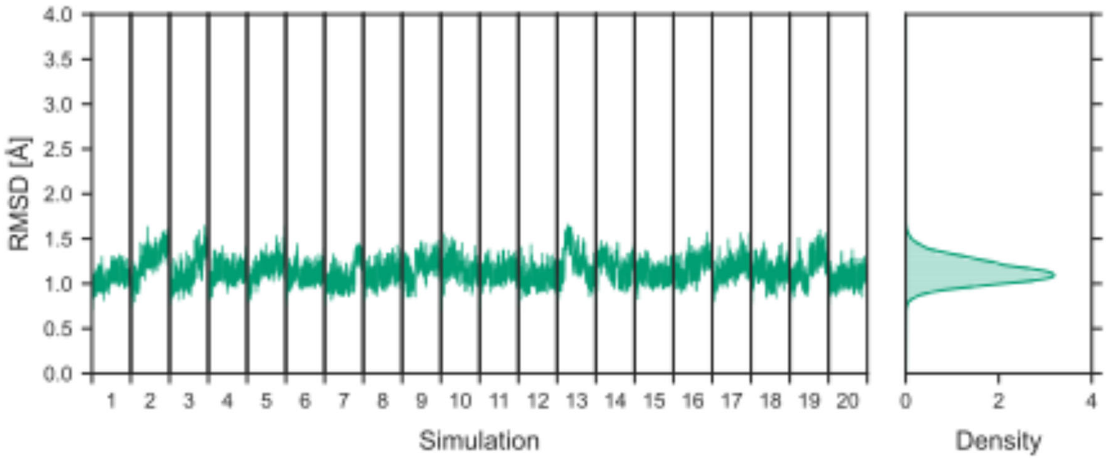


**Figure S6. Structural variability of HepBmut (R144A, R145A, and K147A) CHI3L1 during the fldMD simulations with HS9.** Time courses (left) and probability distributions (right) of the RMSD of CHI3L1 ($C_\alpha$ atoms) shown for all of the 20 simulations (1 µs length). The smoothed distributions (thick, opaque lines) were calculated using a Gaussian kernel density estimator. The underlying data is displayed as a histogram (transparent bars). RMSD values were calculated relative to the crystal structure (PDB ID: 1HJX) after least-squares fitting of the $C_\alpha$ atoms.

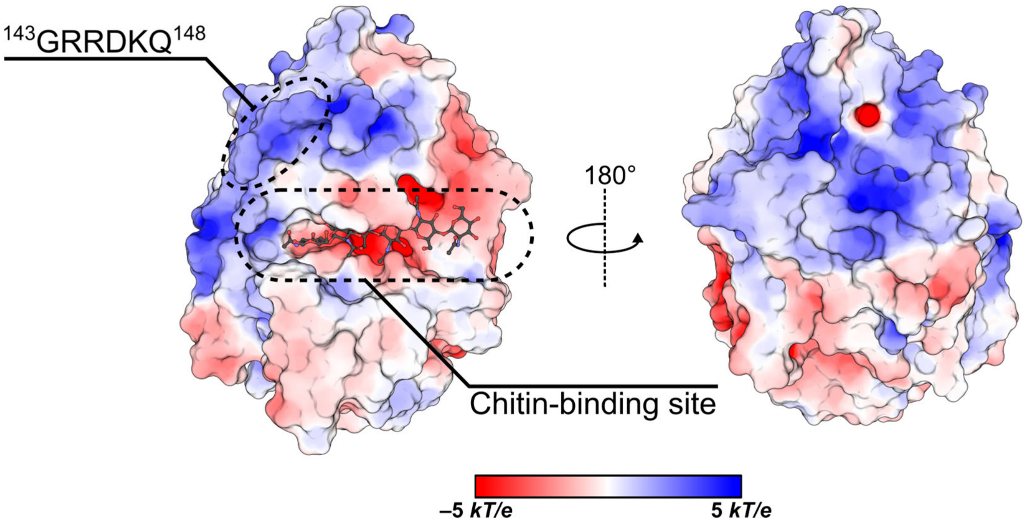


**Figure S7. Electrostatic surface potential of CHI3L1.** The electrostatic surface potential of CHI3L1 (PDB ID: 1HJW) was calculated using the Adaptive Poisson-Boltzmann Solver (APBS) software package (grid spacing: 0.5 Å) in PyMOL and mapped onto the protein surface (color scale in *kT/e*). Dashed circles indicate the $^{143}$GRRDKQ$^{148}$ site and the chitin-binding site ($(GlcNAc)_6$, grey ball-and-stick model).

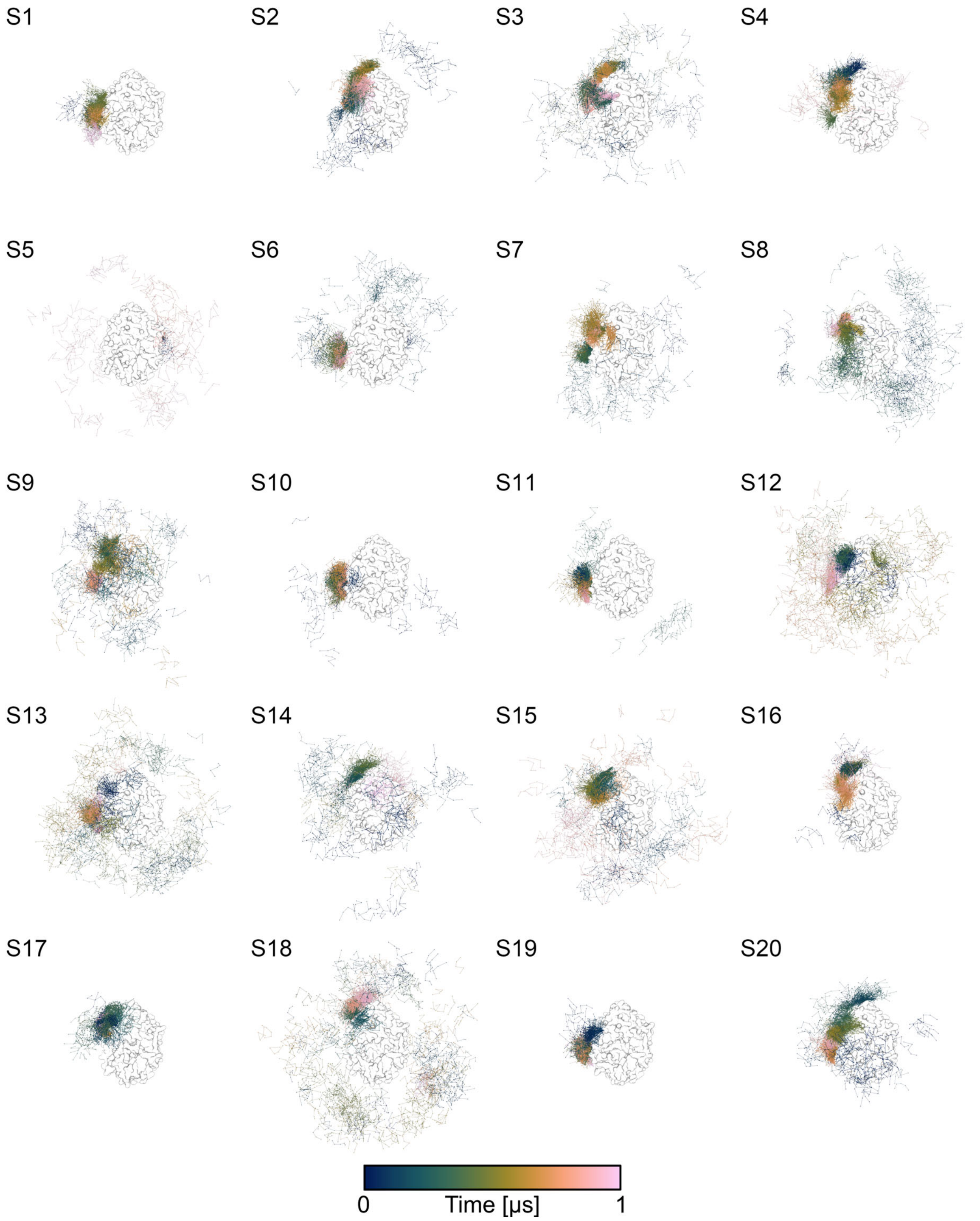


**Figure S8. Unbiased MD simulations of Fondaparinux diffusion with CHI3L1.** The diffusion path taken by the Fondaparinux molecule (ball-and-stick representation) during each of the 20 fldMD simulations, color-coded by the simulation time (gradient: dark blue [0 µs] to light pink [1.0 µs]). Each sphere represents the center of mass (COM) position of the ligand's sugar ring heavy atoms for every tenth trajectory frame. The protein structure is shown as (white) surface representation and the orientation is the same as in Figure S5, left.

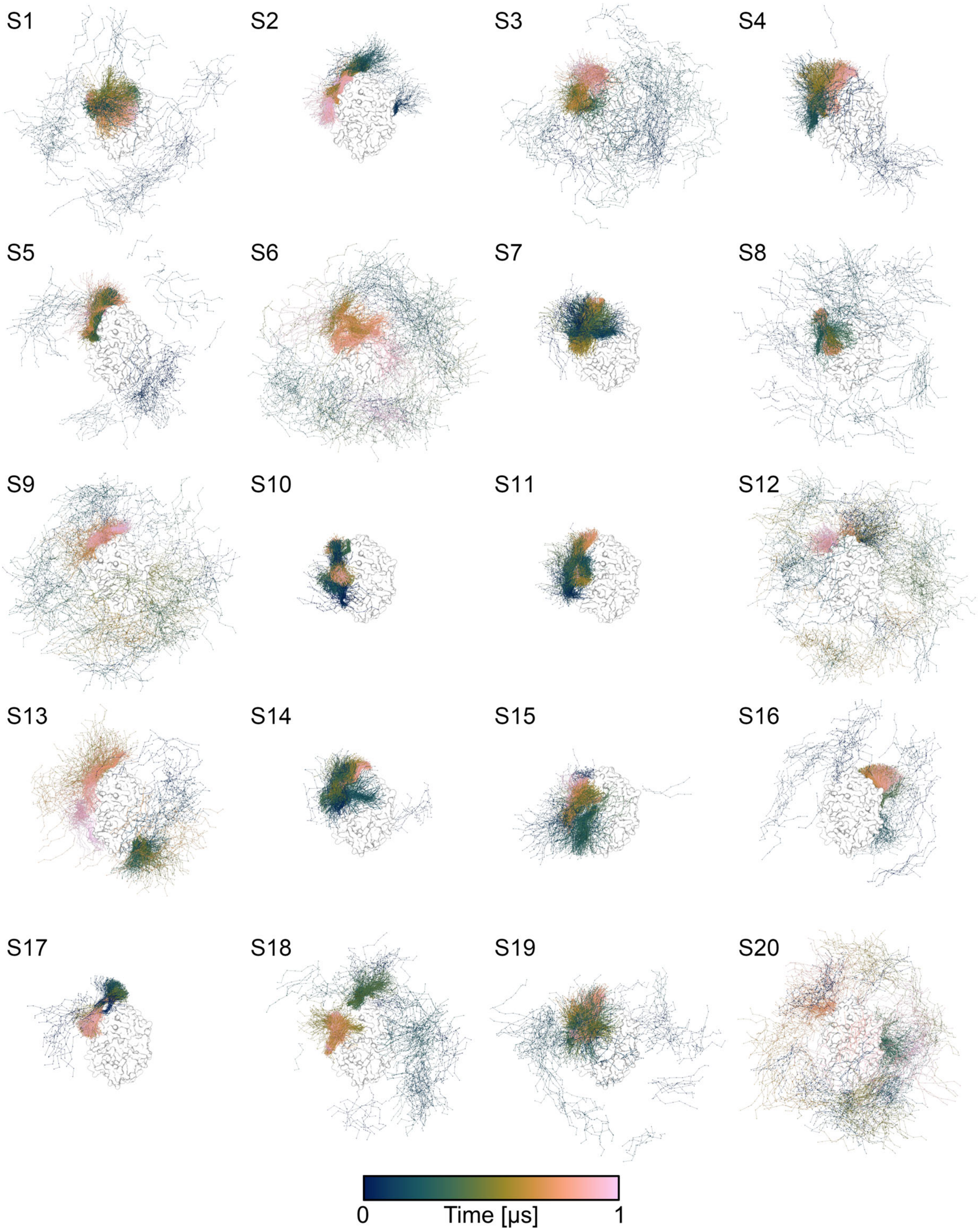


**Figure S9. Unbiased MD simulations of HS9 diffusion with CHI3L1.** The diffusion path taken by the HS9 molecule (ball-and-stick presentation) during each of the 20 fldMD simulations, color-coded by the simulation time (gradient: dark blue [0 µs] to light pink [1.0 µs]). Each sphere represents the center of mass (COM) position of the ligand's sugar ring heavy atoms for every tenth trajectory frame. The protein structure is shown as (white) surface representation and the orientation is the same as in Figure S5, left.

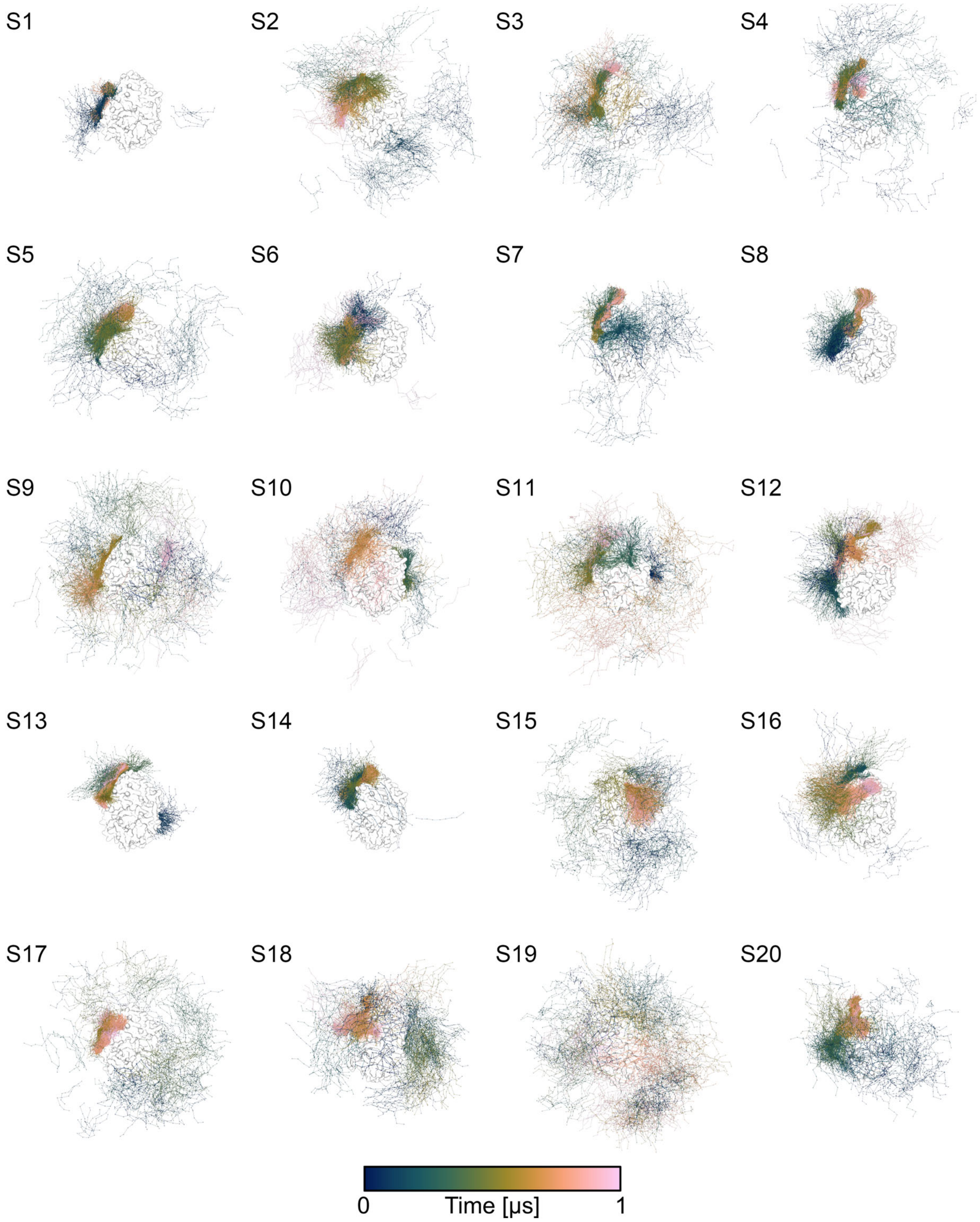


**Figure S10. Unbiased MD simulations of HS9$_{\Delta 2S/6S}$ diffusion with CHI3L1.** The diffusion path taken by the HS9$_{\Delta 2S/6S}$ molecule (ball-and-stick representation) during each of the 20 fldMD simulations color-coded by the simulation time (gradient: dark blue [0 µs] to light pink [1.0 µs]). Each sphere represents the center of mass (COM) position of the ligand's sugar ring heavy atoms for every tenth trajectory frame. The protein structure is shown as (white) surface representation and the orientation is the same as in Figure S5, left.

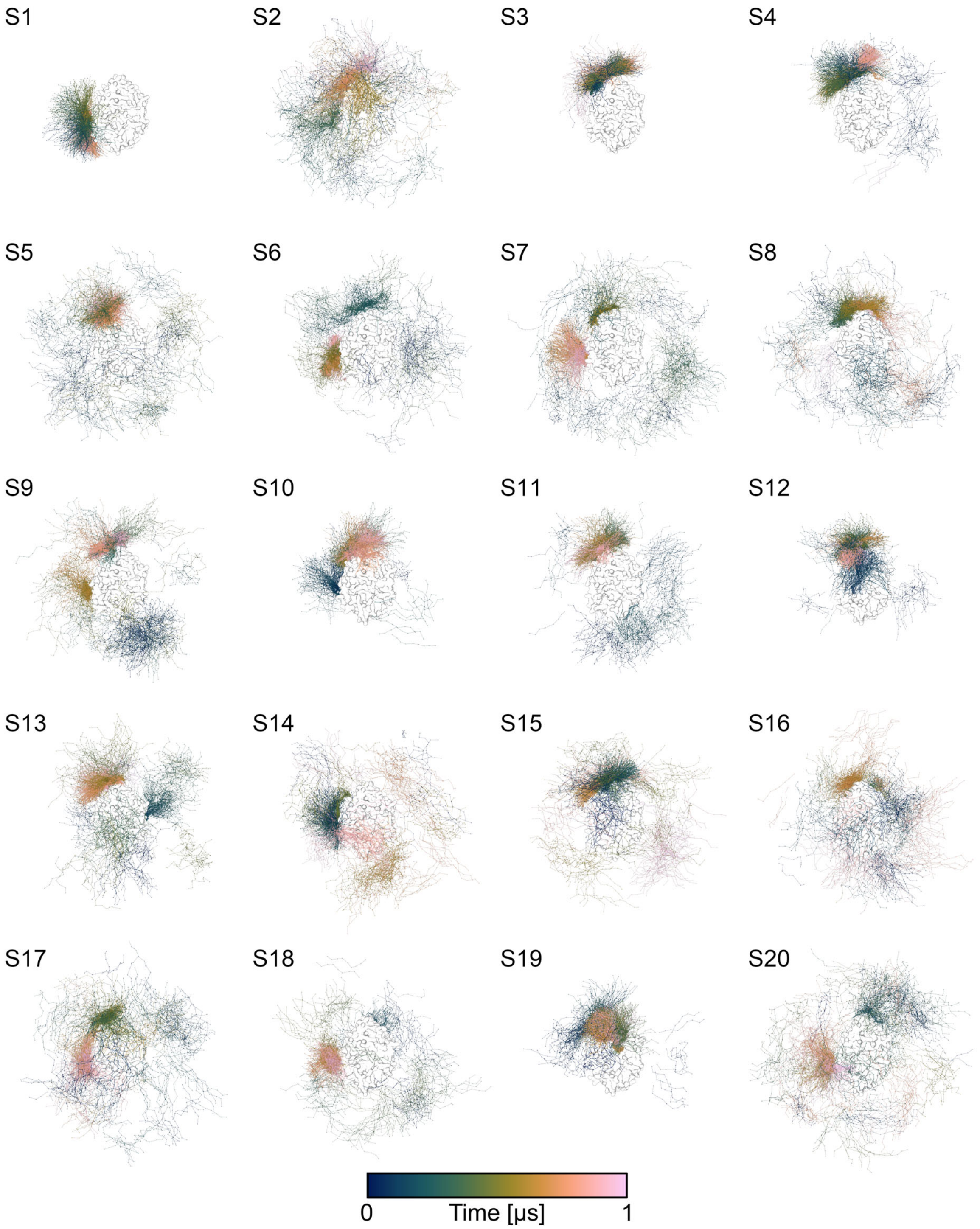


**Figure S11. Unbiased MD simulations of HS9 diffusion with HepBmut (R144A, R145A, and K147A) CHI3L1.** The diffusion path taken by the HS9 molecule (ball-and-stick representation) during each of the 20 fldMD simulations color-coded by the simulation time (gradient: dark blue [0 μs] to light pink [1.0 μs]). Each sphere represents the center of mass (COM) position of the ligand's sugar ring heavy atoms for every tenth trajectory frame. The protein structure is shown as (white) surface representation and the orientation is the same as in Figure S5, left.

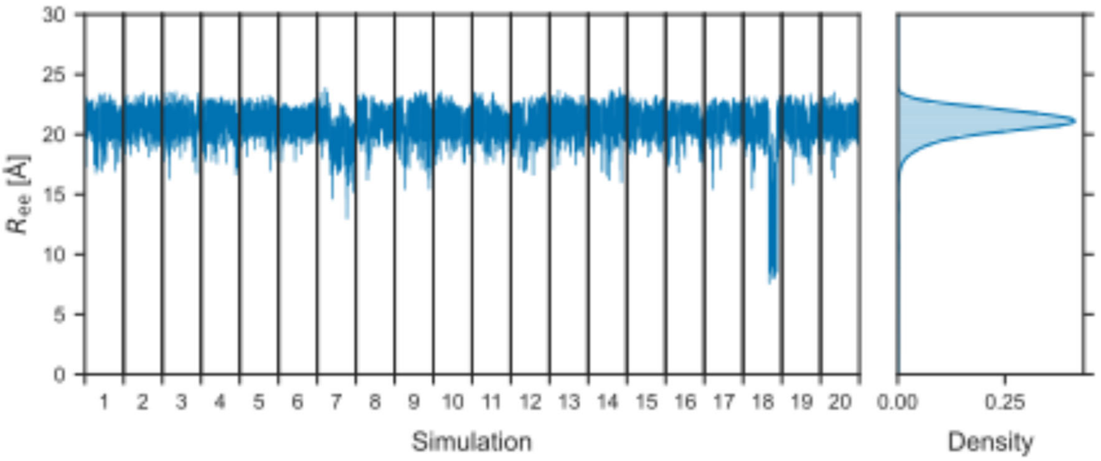


**Figure S12. Conformations of Fondaparinux during the fldMD simulations with CHI3L1.** Time courses (left) and probability distributions (right) of the end-to-end distance ($R_{ee}$) of Fondaparinux shown for all of the 20 simulations (1 µs length). The $R_{ee}$ of Fondaparinux is defined as the distance between the C1 ring carbon of the reducing end to the C4 ring carbon of the nonreducing end. The smoothed distributions (thick, opaque lines) were calculated using a Gaussian kernel density estimator. The underlying data is displayed as a histogram (transparent bars).

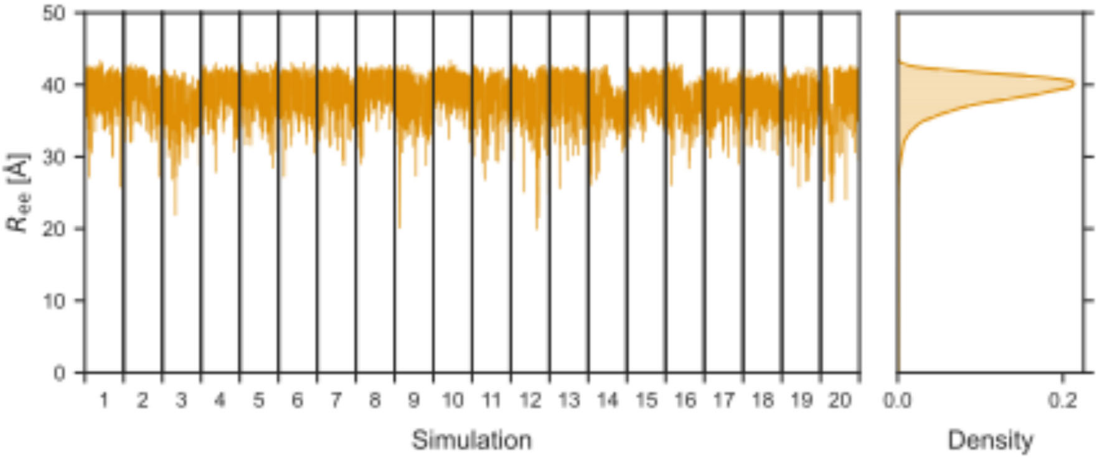


**Figure S13. Conformations of HS9 during the fldMD simulations with CHI3L1.** Time courses (left) and probability distributions (right) of the end-to-end distance ($R_{ee}$) of HS9 shown for all of the 20 simulations (1 µs length). The $R_{ee}$ of HS9 is defined as the distance between the C1 ring carbon of the reducing end to the C4 ring carbon of the nonreducing end. The smoothed distributions (thick, opaque lines) were calculated using a Gaussian kernel density estimator. The underlying data is displayed as a histogram (transparent bars).

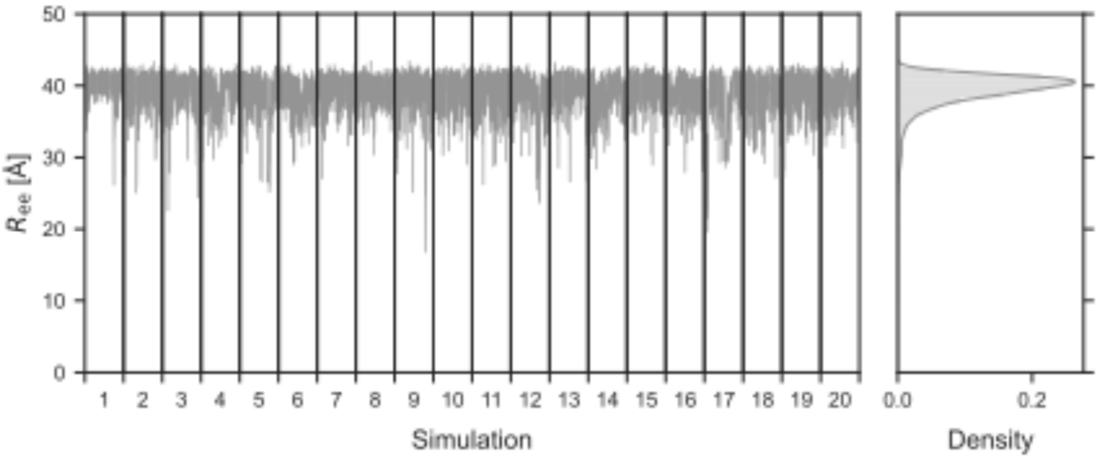


**Figure S14. Conformations of HS9$_{\Delta 2S/6S}$ during the fldMD simulations with CHI3L1.** Time courses (left) and probability distributions (right) of the end-to-end distance ($R_{ee}$) of HS9$_{\Delta 2S/6S}$ shown for all of the 20 simulations (1 µs length). The $R_{ee}$ of HS9$_{\Delta 2S/6S}$ is defined as the distance between the C1 ring carbon of the reducing end to the C4 ring carbon of the nonreducing end.The smoothed distributions (thick, opaque lines) were calculated using a Gaussian kernel density estimator. The underlying data is displayed as a histogram (transparent bars).

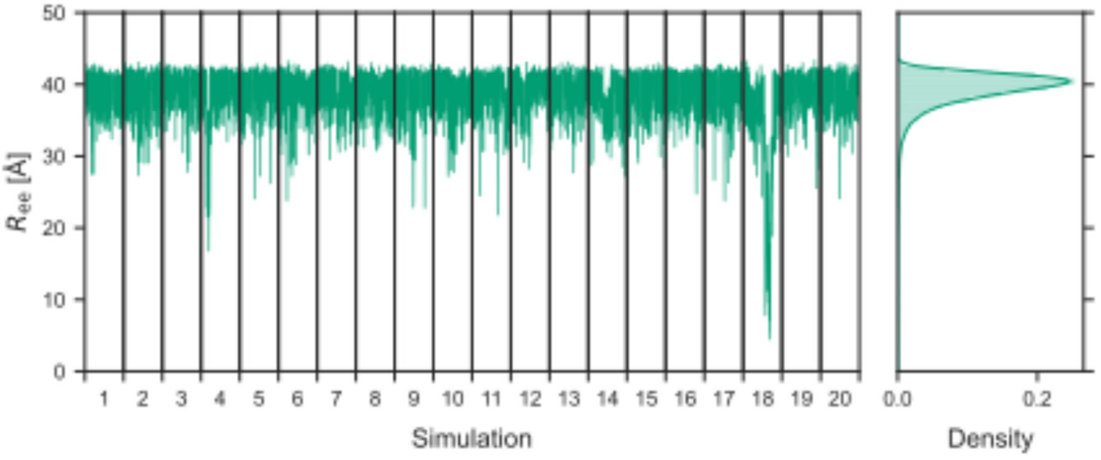


**Figure S15. Conformations of HS9 during the fldMD simulations with HepBmut (R144A, R145A, and K147A) CHI3L1.** Time courses (left) and probability distributions (right) of the end-to-end distance ($R_{ee}$) of HS9 shown for all of the 20 simulations (1 µs length). The $R_{ee}$ of HS9 is defined as the distance between the C1 ring carbon of the reducing end to the C4 ring carbon of the nonreducing end. The smoothed distributions (thick, opaque lines) were calculated using a Gaussian kernel density estimator. The underlying data is displayed as a histogram (transparent bars).

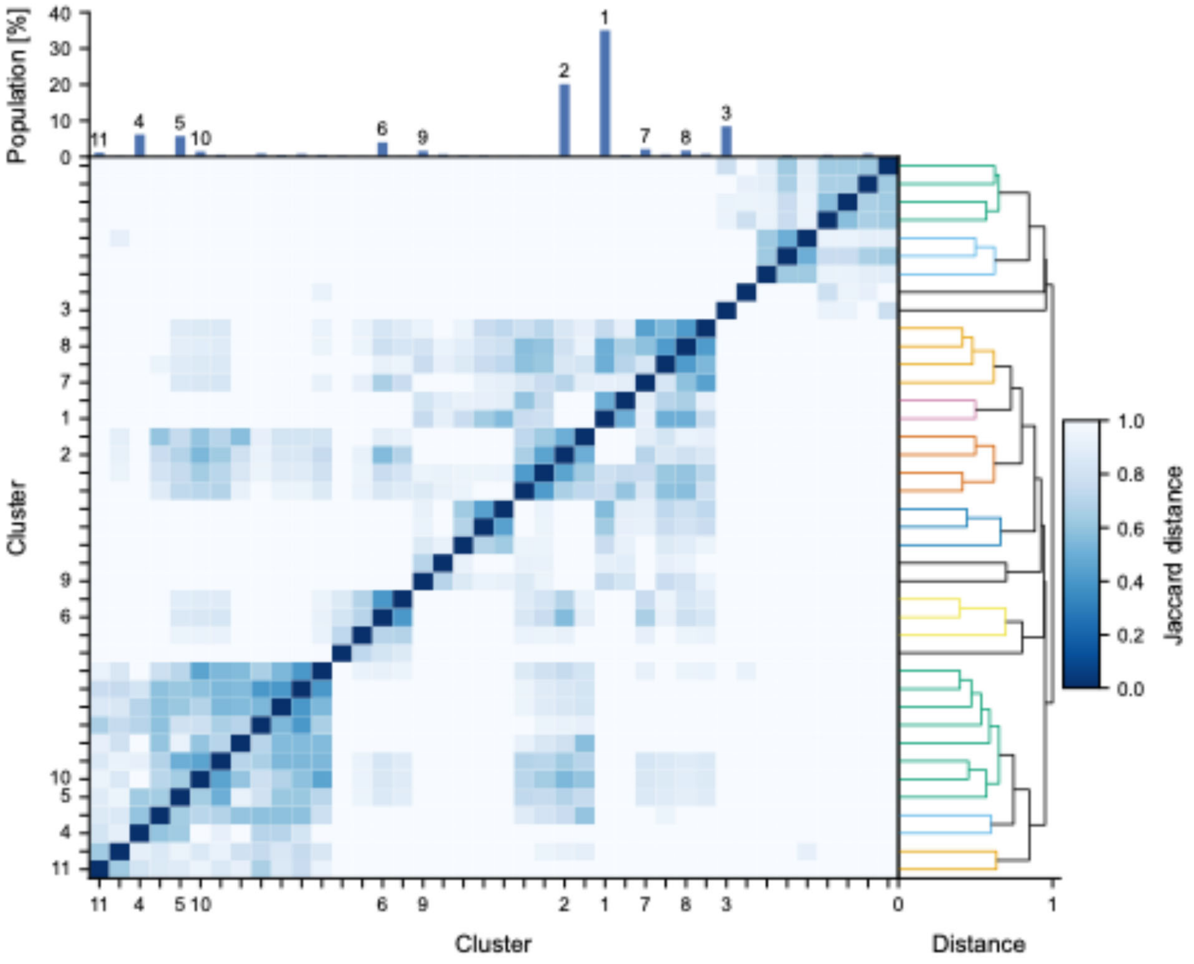


**Figure S16. Clustering analysis of Fondaparinux poses based on interaction fingerprints with CHI3L1.** Similarity and hierarchical relationships are shown for the top 40 clusters (representing populations > 0.1% of total samples). Representative binary profiles for each cluster were generated by applying a 30% interaction frequency threshold. The heatmap illustrates the pairwise Jaccard distance between these representative profiles, where a distance of 0 indicates identical interaction patterns. The corresponding dendrogram was generated via average linkage hierarchical clustering. Clusters are labeled sequentially in the order of decreasing population (1 = largest) for clusters with a population > 1.0%.

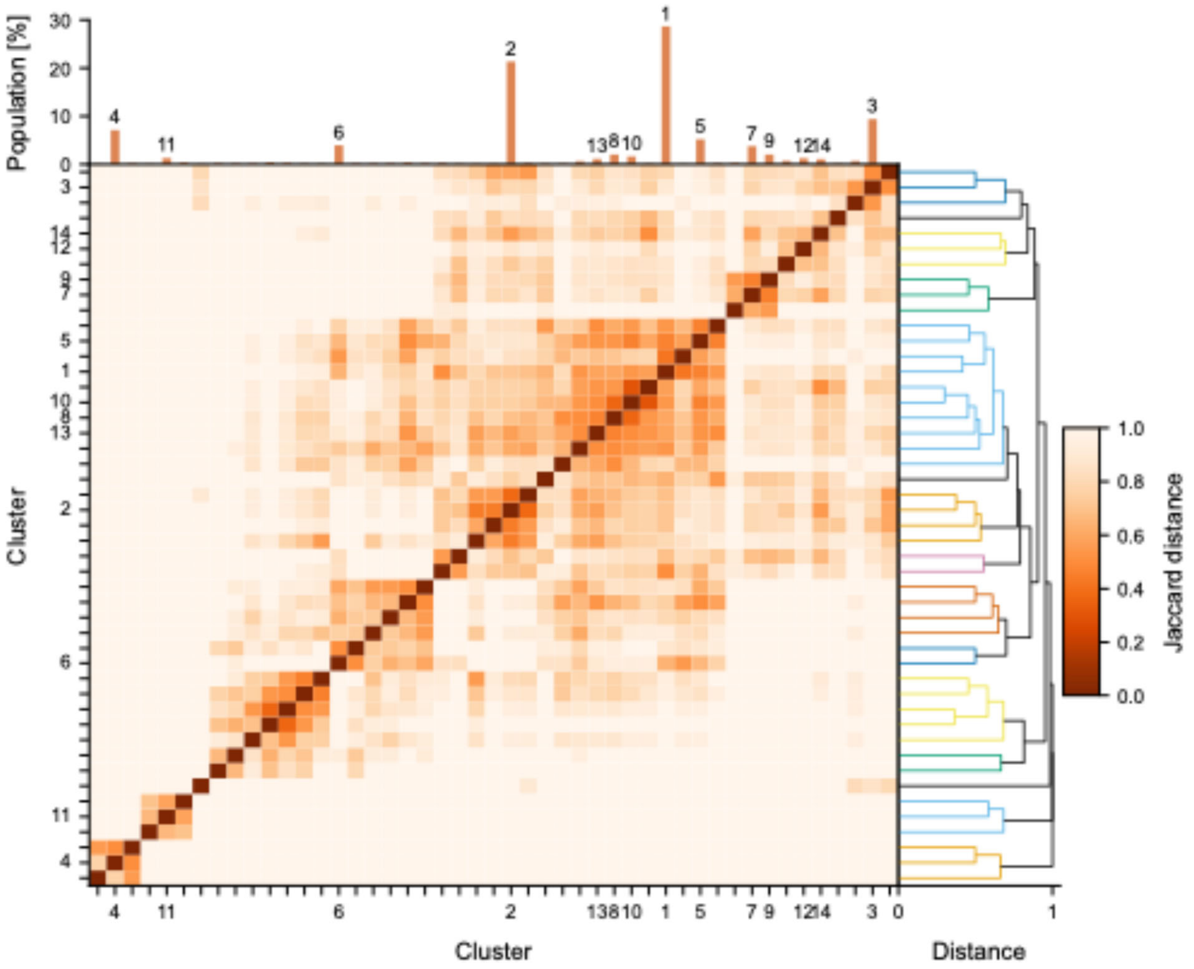


**Figure S17. Clustering analysis of HS9 poses based on interaction fingerprints with CHI3L1.** Similarity and hierarchical relationships are shown for the top 47 clusters (representing populations > 0.1% of total samples). Representative binary profiles for each cluster were generated by applying a 30% interaction frequency threshold. The heatmap illustrates the pairwise Jaccard distance between these representative profiles, where a distance of 0 indicates identical interaction patterns. The corresponding dendrogram was generated via average linkage hierarchical clustering. Clusters are labeled sequentially in the order of decreasing population (1 = largest) for clusters with a population > 1.0%.

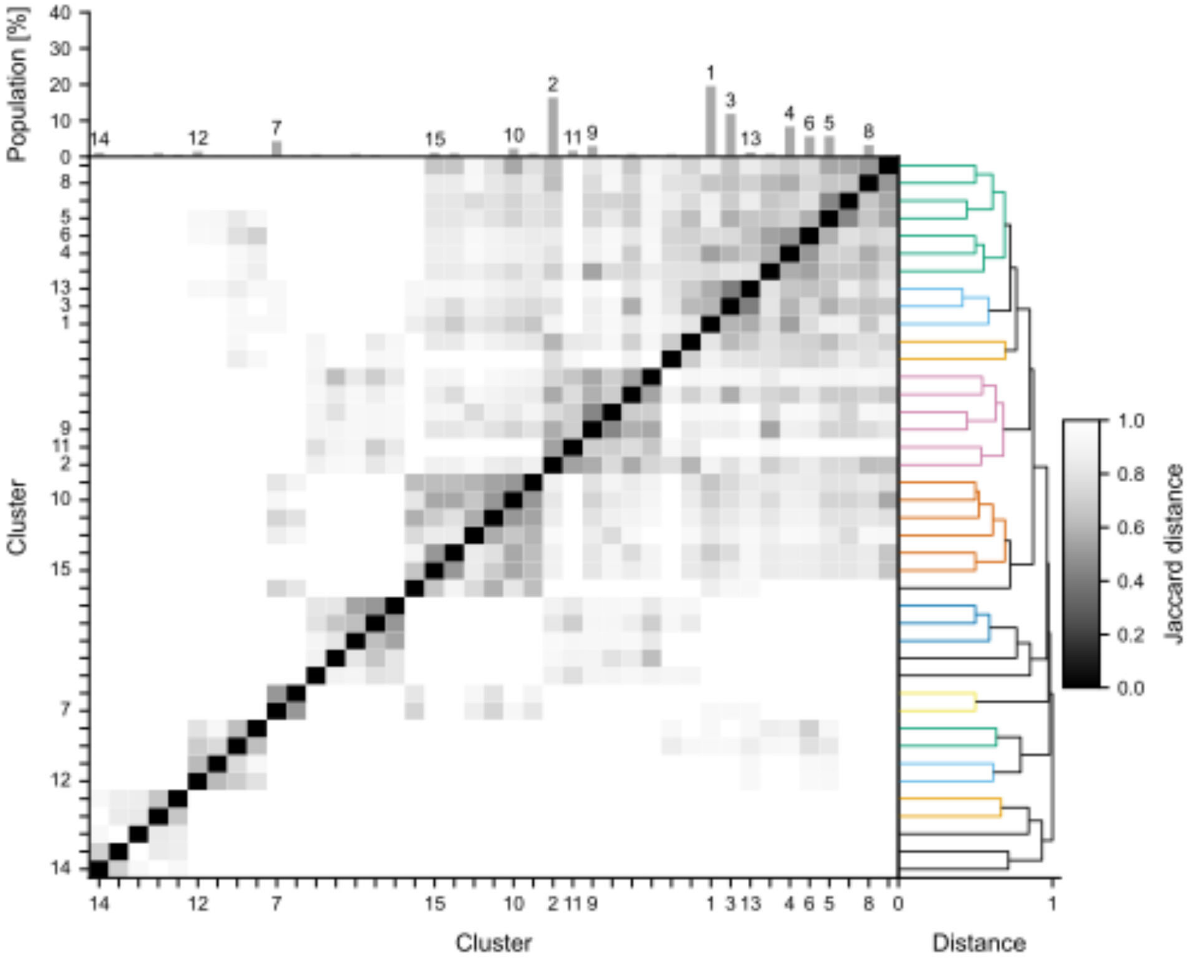


**Figure S18. Clustering analysis of HS9$_{\Delta 2S/6S}$ poses based on interaction fingerprints with CHI3L1.** Similarity and hierarchical relationships are shown for the top 41 clusters (representing populations > 0.1% of total samples). Representative binary profiles for each cluster were generated by applying a 30% interaction frequency threshold. The heatmap illustrates the pairwise Jaccard distance between these representative profiles, where a distance of 0 indicates identical interaction patterns. The corresponding dendrogram was generated via average linkage hierarchical clustering. Clusters are labeled sequentially in the order of decreasing population (1 = largest) for clusters with a population > 1.0%.

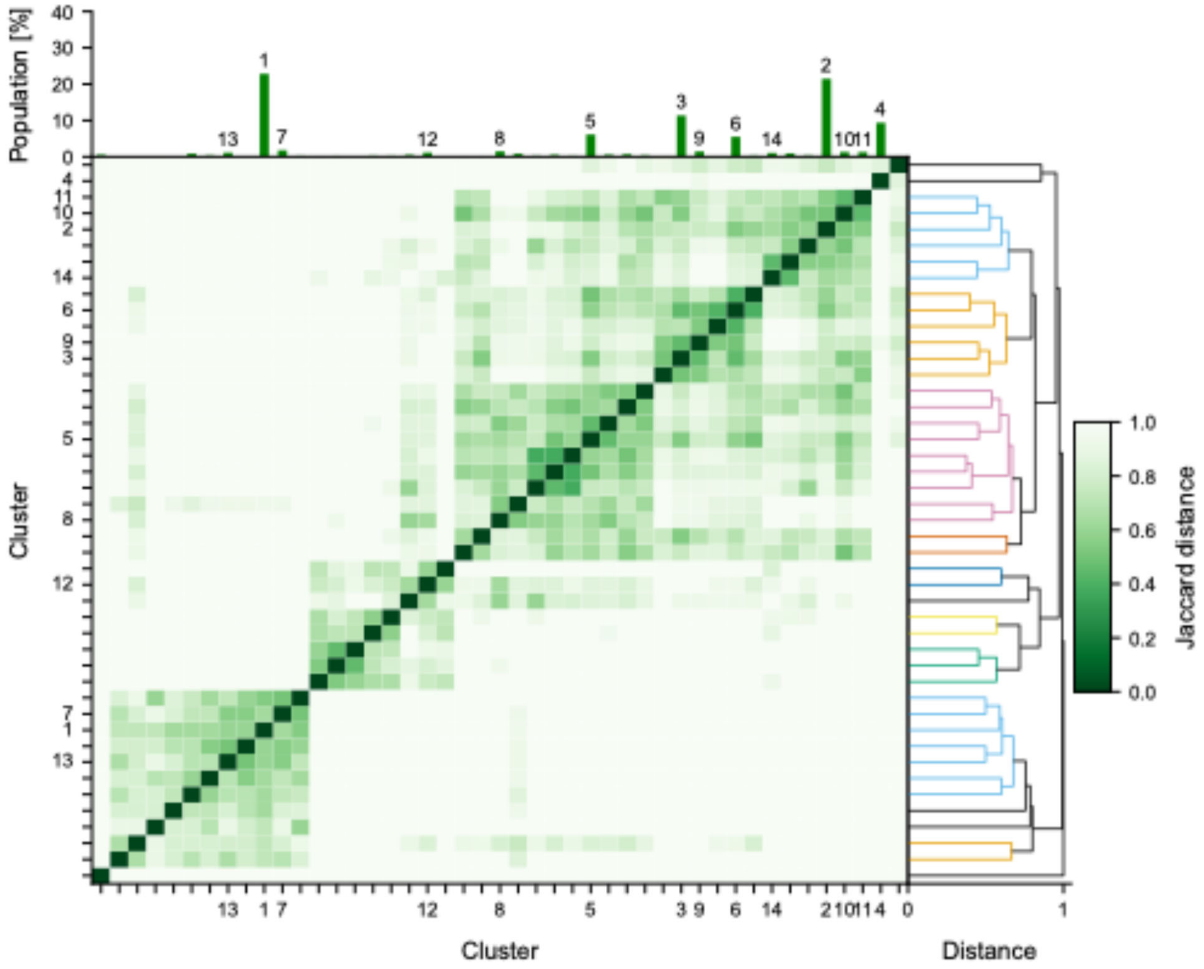


**Figure S19. Clustering analysis of HS9 poses based on interaction fingerprints with HepBmut (R144A, R145A, and K147A) CHI3L1.** Similarity and hierarchical relationships are shown for the top 45 clusters (representing populations > 0.1% of total samples). Representative binary profiles for each cluster were generated by applying a 30% interaction frequency threshold. The heatmap illustrates the pairwise Jaccard distance between these representative profiles, where a distance of 0 indicates identical interaction patterns. The corresponding dendrogram was generated via average linkage hierarchical clustering. Clusters are labeled sequentially in the order of decreasing population (1 = largest) for clusters with a population > 1.0%.

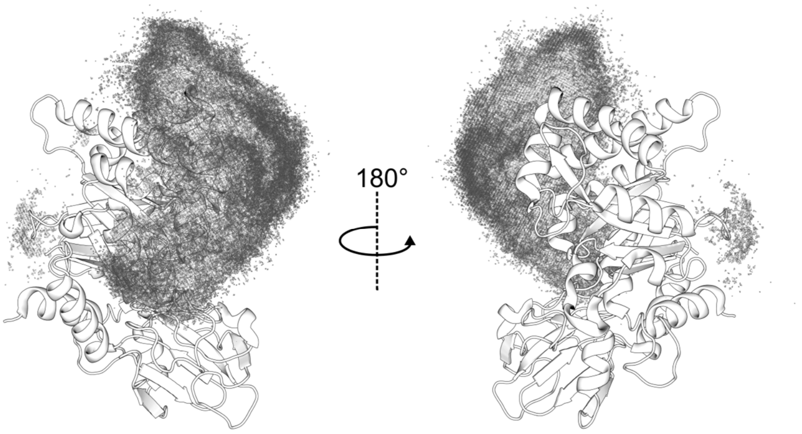


**Figure S20. Probability density of HS9$_{\Delta2S/6S}$ around CHI3L1.** The derived occupancy densities from all fldMD simulations for HS9$_{\Delta2S/6S}$ (red mesh) are mapped around the CHI3L1 crystal structure (white cartoon; left = same orientation as in Figure 1A; PDB ID: 1HJW). For the occupancy density contour level, see the Materials and Methods section.

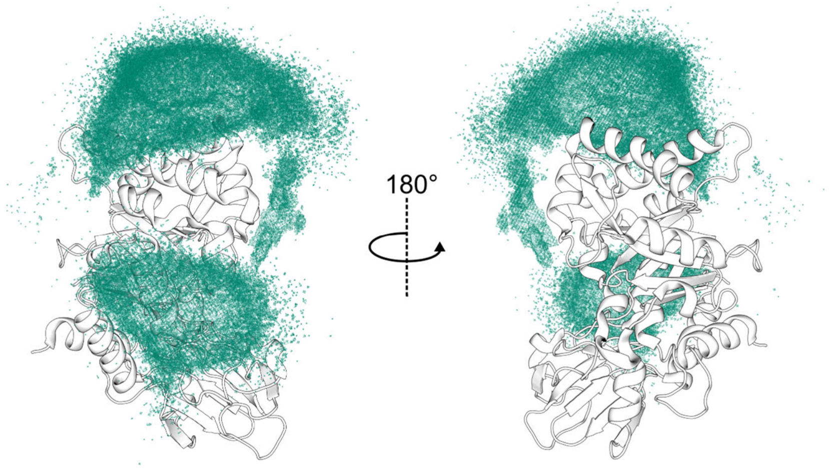


**Figure S21. Probability density of HS9 around HepBmut (R144A, R145A, and K147A) CHI3L1.** The derived occupancy densities from all fldMD simulations for HS9$_{\Delta2S/6S}$ (green mesh) are mapped around the CHI3L1 crystal structure (white cartoon; left = same orientation as in Figure 1A; PDB ID: 1HJW). For the occupancy density contour level, see the Materials and Methods section.

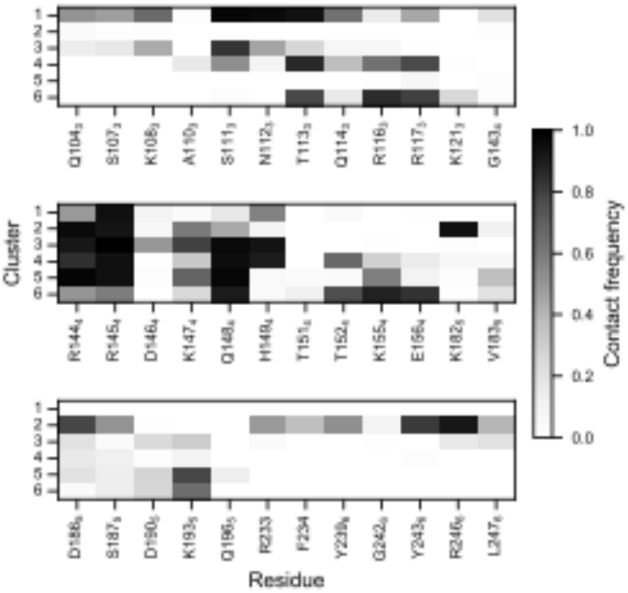


**Figure S22. Contacts between CHI3L1 and $HS9_{\Delta 2S/6S}$.** Heatmap illustrating the residue-wise contact frequencies between CHI3L1 and $HS9_{\Delta 2S/6S}$. Data are shown for the six most populated clusters, ranked by size (1 = highest population) (see also Figure S16). Only residues exhibiting a contact frequency >10% in at least one cluster are shown. Subscript numbers indicate the α-helix to which each residue belongs.

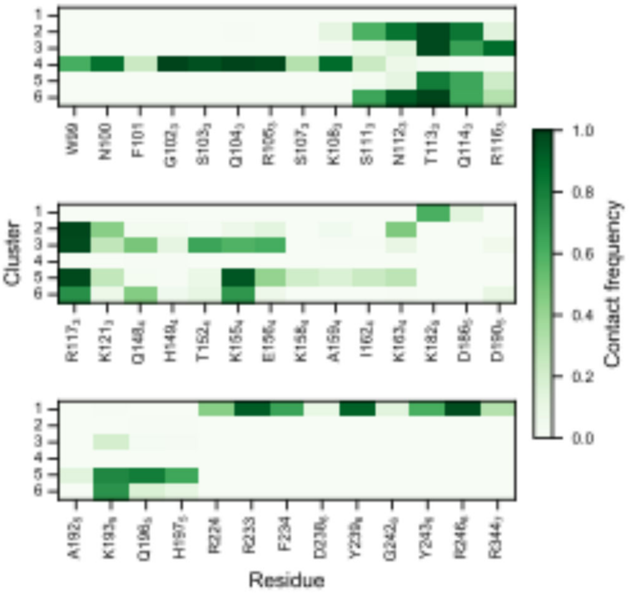


**Figure S23. Contacts between HepBmut (R144A, R145A, and K147A) CHI3L1 and HS9.** Heatmap illustrating the residue-wise contact frequencies between hCHI3L1 (R144A, R145A, and K147A) and HS9. Data are shown for the six most populated clusters, ranked by size (1 = highest population) (see also Figure S17). Only residues exhibiting a contact frequency >10% in at least one cluster are shown. Subscript numbers indicate the α-helix to which each residue belongs.

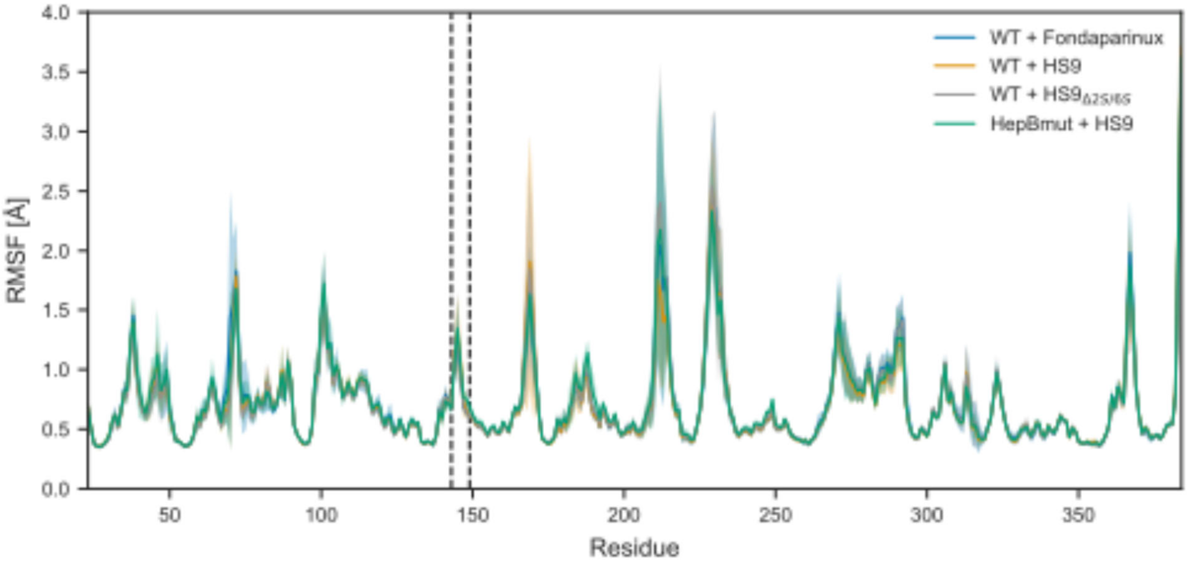


**Figure S24. Atomic fluctuations calculated from MD simulations.** Average per-residue $C_\alpha$ root-mean-square fluctuation (RMSF; opaque lines) for each system over all of the 20 fldMD simulations. Shaded areas represent the standard deviation around the mean. RMSF values were calculated relative to the crystal structure (PDB ID: 1HJX) after least-squares fitting of the $C_\alpha$ atoms. Vertical dashed lines denote the $^{143}$GRRDKQ$^{148}$ motif.

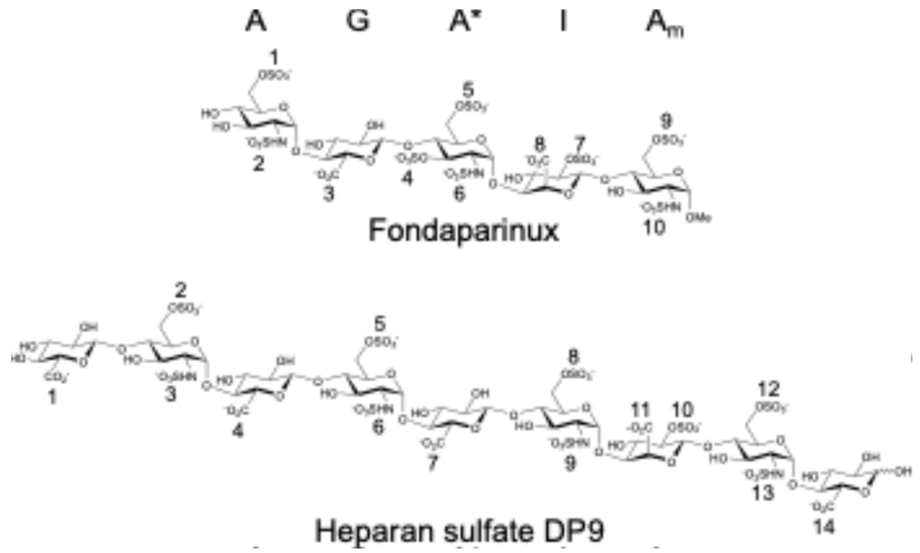


**Figure 25. Numbering of the anionic groups of Fondaparinux and HS9 for reference in Figures S26-29.**

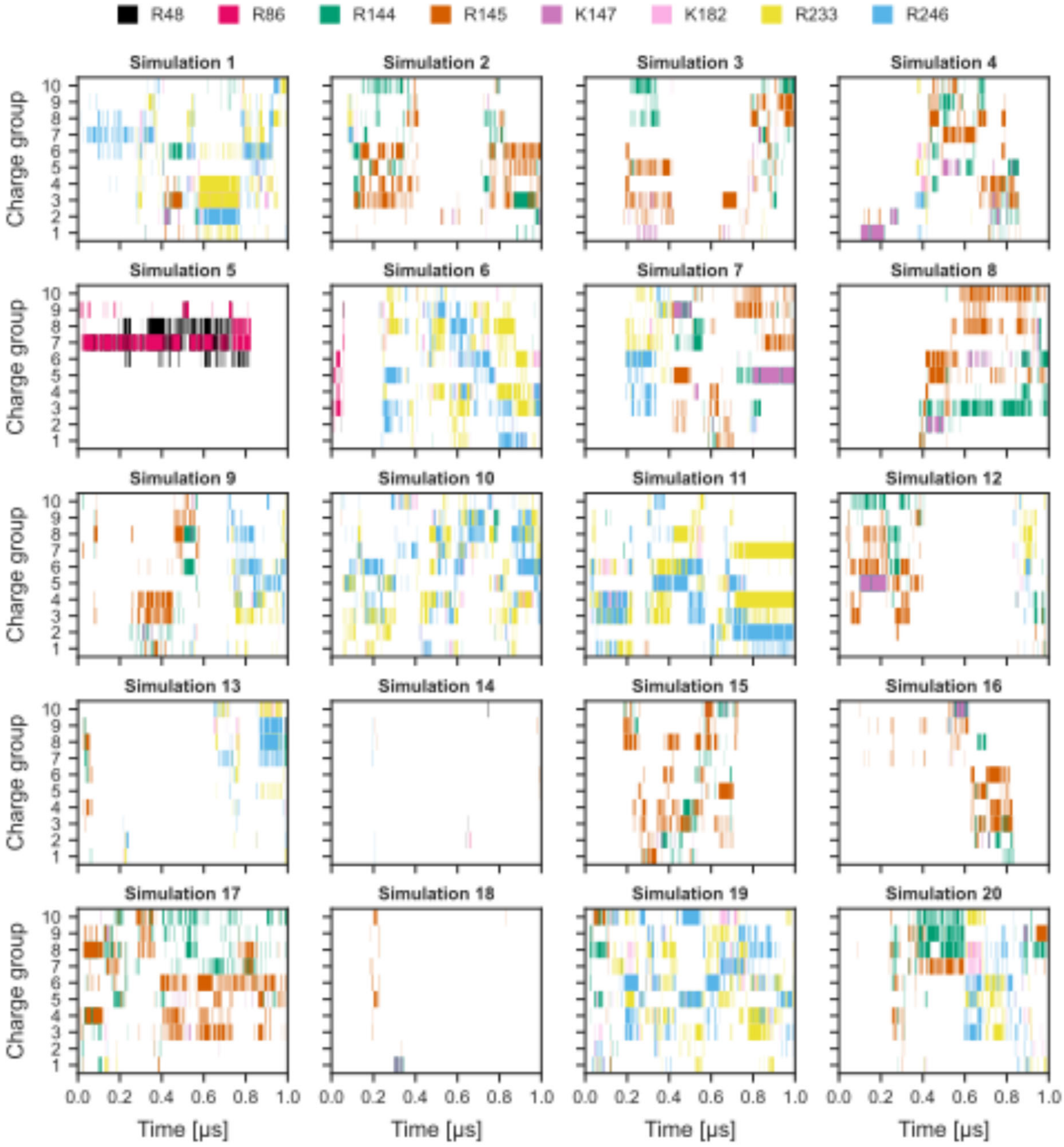


**Figure S26. Time-series analysis of Fondaparinux interactions with CHI3L1.** Time courses of the binding (heavy-atom distance < 4 Å) between the anionic groups of Fondaparinux (sulfate and carboxylate; see Figure S25 for numbering) and specific cationic residues of CHI3L1 identified via clustering analysis. Binding to individual residues is colored as follows: R48 (black), R86 (magenta), R144 (green), R145 (dark orange), K147 (violet), K182 (pink), R233 (yellow), and R246 (light blue).

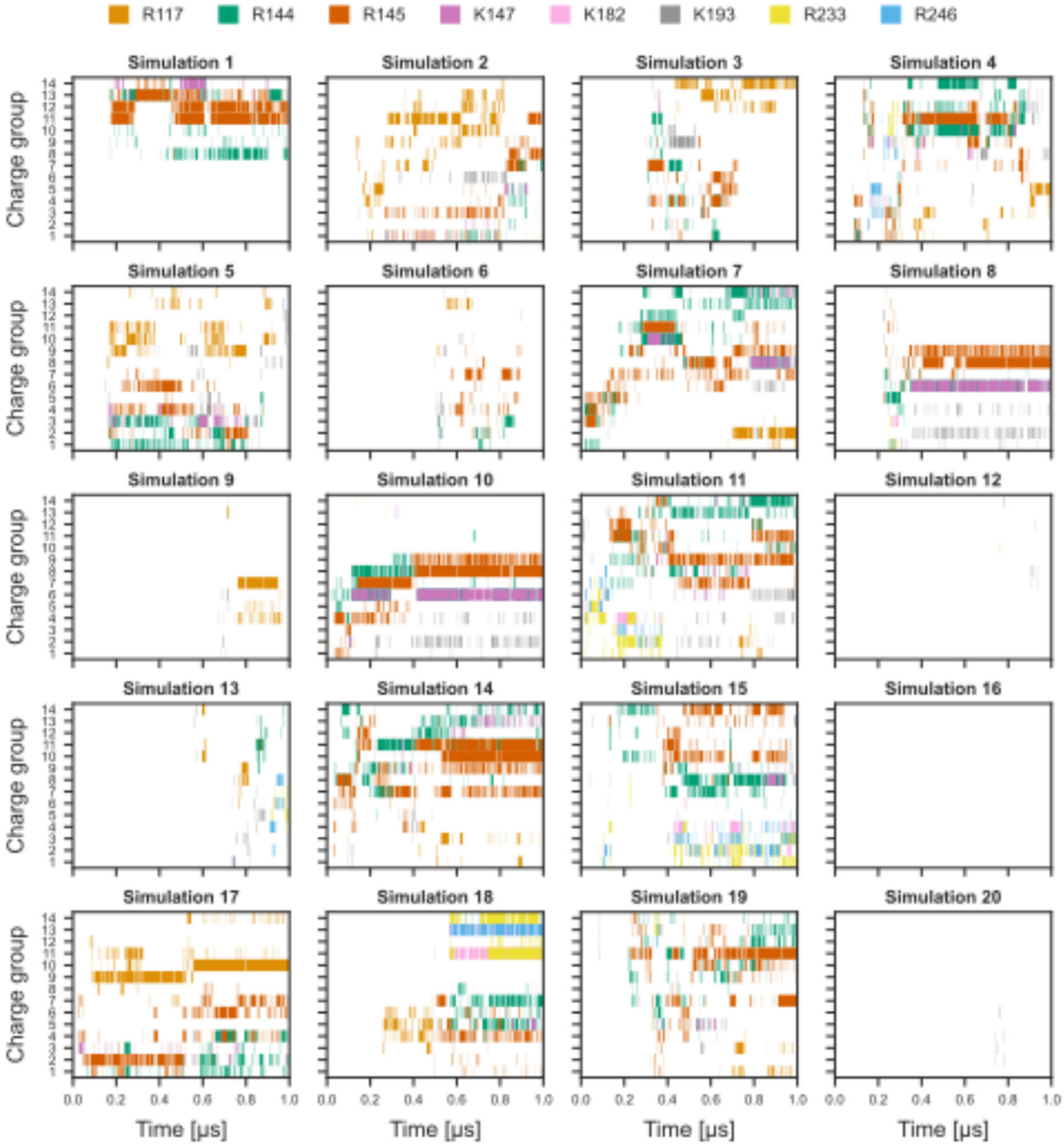


**Figure S27. Time-series analysis of HS9 interactions with CHI3L1.** Time courses of the binding (heavy-atom distance < 4 Å) between the anionic groups of HS9 (sulfate and carboxylate; see Figure S25 for numbering) and specific cationic residues of CHI3L1 identified via clustering analysis. Binding to individual residues is colored as follows: R117 (orange), R144 (green), K145 (dark orange), K147 (violet), K182 (pink), and K193 (gray), R233 (yellow), and R246 (light blue).

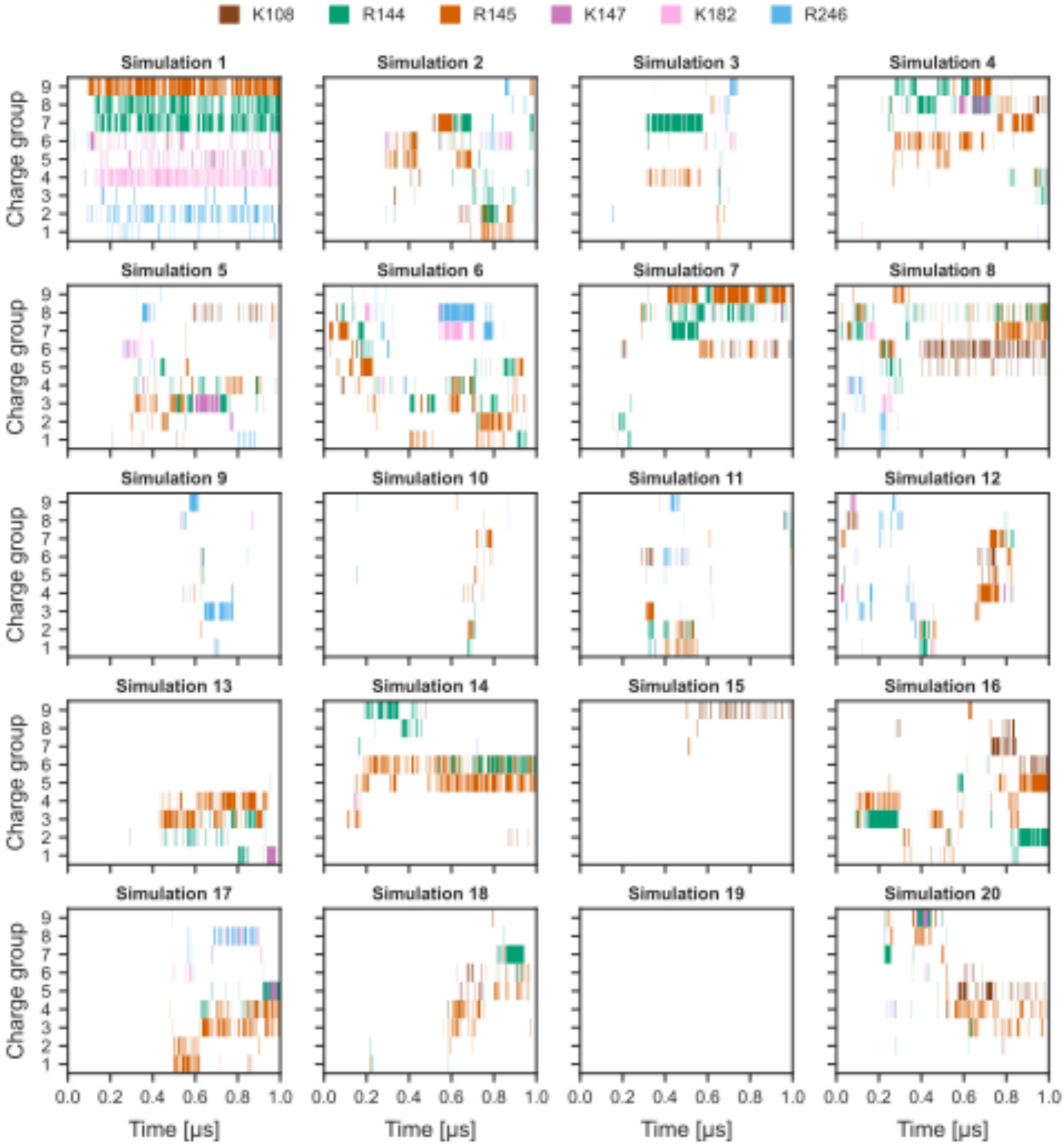


**Figure S28. Time-series analysis of HS9$_{\Delta 2S/6S}$ interactions with CHI3L1.** Time courses of the binding (heavy-atom distance < 4 Å) between the anionic groups of HS9$_{\Delta 2S/6S}$ (sulfate and carboxylate; see Figure S25 for numbering) and specific cationic residues of CHI3L1 identified via clustering analysis. Binding to individual residues is colored as follows: K108 (dark brown), R144 (green), K145 (dark orange), K147 (violet), K182 (pink), and R246 (light blue).

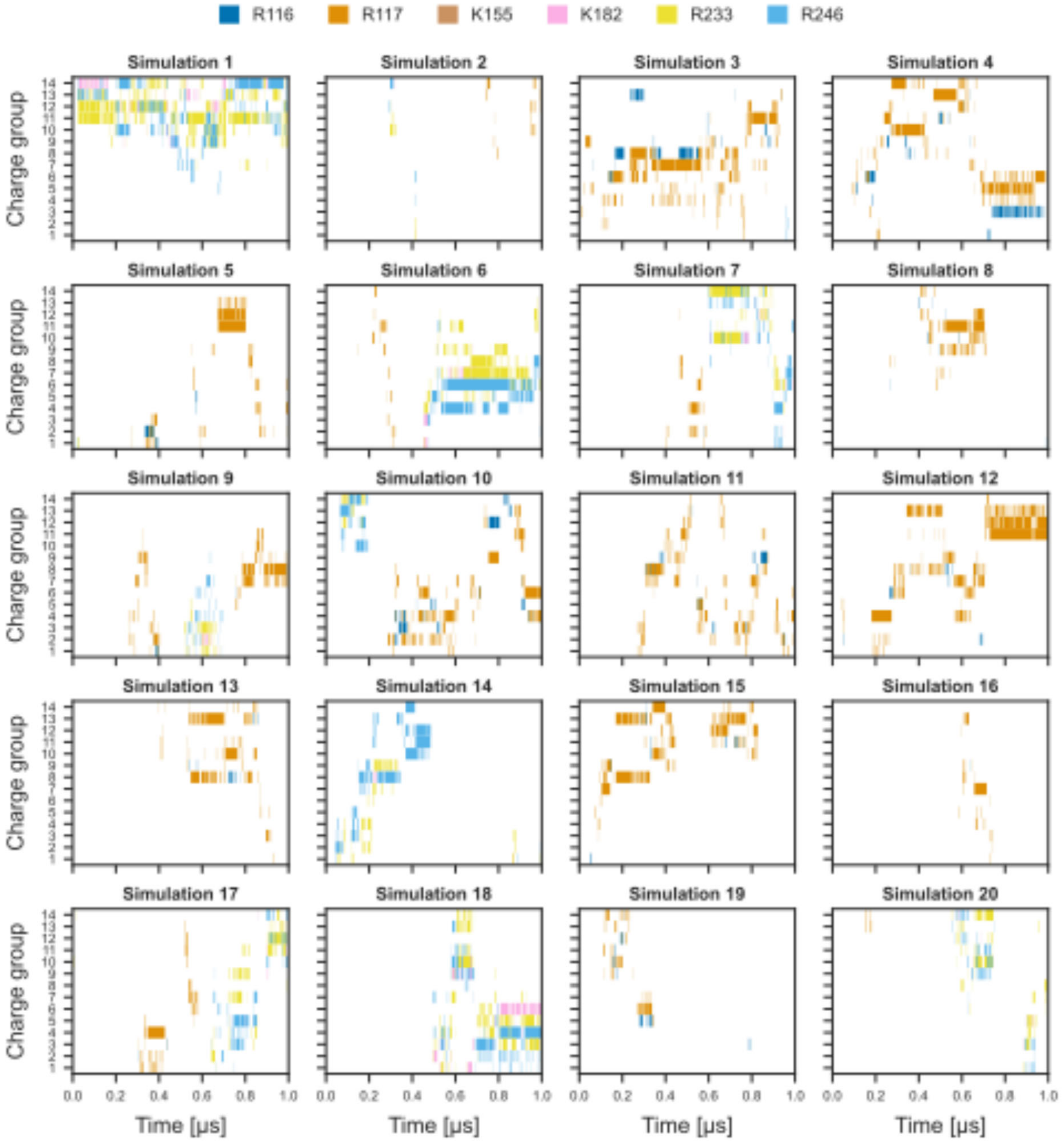


**Figure S29. Time-series analysis of HS9 interactions with HepBmut (R144A, R145A, and K147A) CHI3L1.** Time courses of the binding (heavy-atom distance < 4 Å) between the anionic groups of HS9 (sulfate and carboxylate; see Figure S25 for numbering) and specific cationic residues of CHI3L1 identified via clustering analysis. Binding to individual residues is colored as follows: R116 (blue), R117 (orange), K155 (brown), K182 (pink), R233 (yellow), and R246 (light blue).

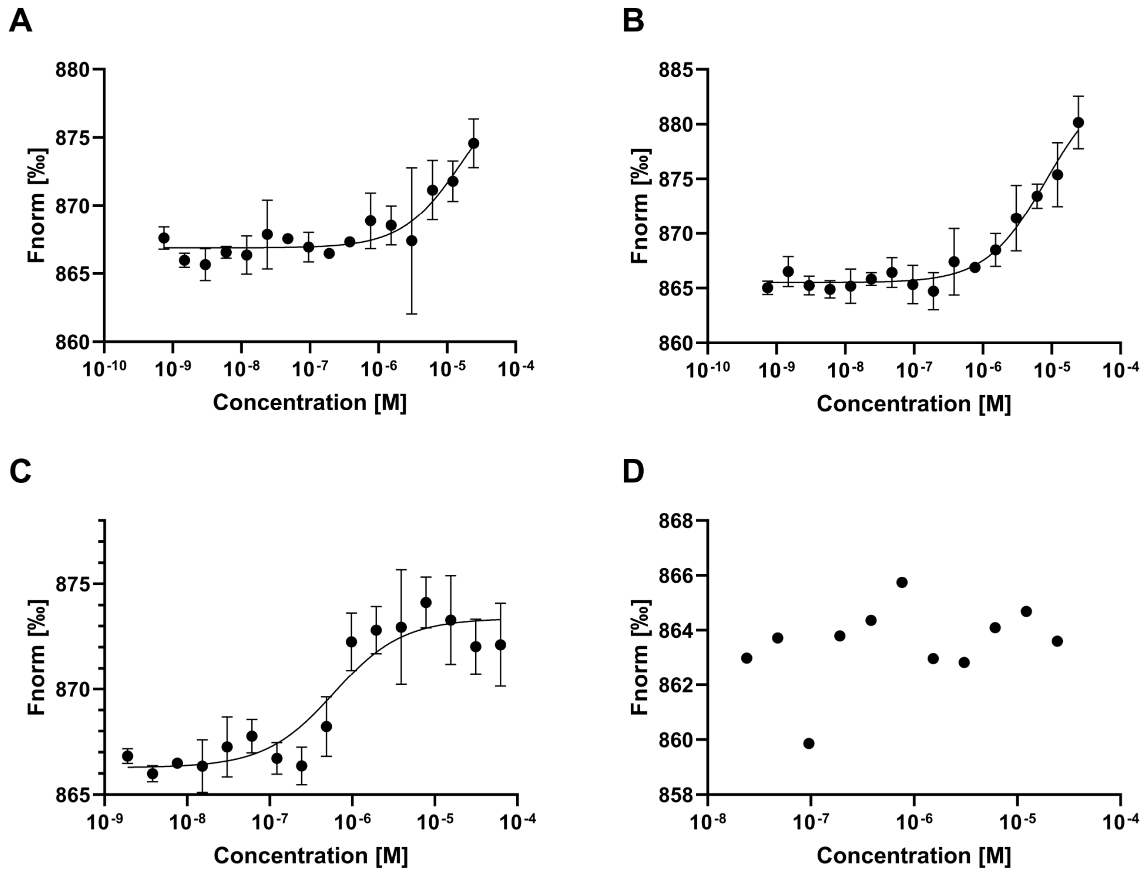


**Figure S30. MST binding isotherms of the interaction between CHI3L1 and galectin-3.** (A) CHI3L1 WT binding to galectin-3, (B) CBmut CHI3L1 binding to galectin-3. (C) CHI3L1 binding to galectin-3 in the presence of 500 µM Fondaparinux. (D) CHI3L1 binding to galectin-3 in presence of 500 µM $(GlcNAc)_6$.
